# Opportunity nest egg: insights on the nutritional ecology, life history and captive management of three species of kiwi (*Apteryx* spp.) chick from Operation Nest Egg zoo hand-rearing records

**DOI:** 10.1101/2023.10.31.564897

**Authors:** L.J. Gray, B.A. Mitchell, I.L. Milner-Bradford, L. Keller, G. Bell, K.J. McGuire, C. Travers

## Abstract

Zoo data collected by keepers while looking after endangered species are increasingly recognised as important scientific resources. As chicks, New Zealand’s threatened kiwi (*Apteryx* spp.) are subject to the protective conservation programme Operation Nest Egg (ONE), during which growth, developmental and life history data are recorded. We have conducted comparative analyses on hand rearing records from 306 chicks, from Coromandel, Eastern and Western populations of Brown kiwi (*A. mantelli*), and rowi (*A. rowi*) and Haast tokoeka (*A. australis* “Haast”). We analysed chick responses to an old diet *vs*. a new kiwi diet, introduced in 2017. Both diets are fixed nutrient ratio mixtures. The old diet was high-protein, low-energy, while the new diet is high-fat, high-energy, with better micronutrient levels. We found Coromandel chicks, who originate from the environmentally variable Kūaotunu Peninsula, grow the most efficiently overall on either diet, indicating they may be nutritional generalists. Western and Eastern chicks’ growth efficiency was intermediate, while the South Island species grew the least efficiently on either diet.

Rowi chicks developed the fastest overall on either diet, while Haast tokoeka chicks developed the most slowly, especially on the new diet. Rowi chicks therefore had to eat large volumes of either diet over a short time to maintain their rapid development, while Haast chicks were required to eat large volumes, especially of the new diet, over a protracted developmental period. This situation may have led South Island chicks to over-consume one or another diet component, with likely health consequences. Neither diet was obviously superior for chick wellbeing overall, though the new diet better supported chicks that needed hand feeding. This work demonstrates different genetic populations of kiwi differ in their physiological responses to nutrition. As ONE is ongoing, tailored diets for chicks from each genetic group should be developed, and we present methods to achieve this. In our life history trait analyses, we found chick starting size (hatch mass) did not significantly influence growth efficiency across kiwi genetic group, nor did chick sex. We identified that chicks malpositioned as embryos were more likely to require extended periods of hand feeding, and that Eastern males produce more malpositioned embryos than other populations. Our study shows that effective zoo records can be used to improve captive care, to stimulate future research to refine species management practices, and to explore fundamental questions of life history evolution in wild and captive populations.

## INTRODUCTION

The meticulous husbandry records collected by zoo keepers on the captive animals in their care are increasing recognised as important research resources. Scientific analysis of zoo keeping data has revealed insights into the evolutionary strategies and life history adaptations of wide-range of animals (Ryder & Feistner 1995) including testudines, primates, and other mammals (Da Silva et al. 2022; Zehr et al. 2014; de Visser et al. 2022). Zoo data have proven particularly useful when studying species that are rare, and when the analysis of their zoo-data can teach us how to improve their captive husbandry and their fundamental biology (Scheun et al. 2020; Farquharson et al. 2021).

Four of the five species of Aotearoa New Zealand’s threatened kiwi are subject to a protective captive conservation management programme called Operation Nest Egg (ONE) from which these kinds of instructive data are generated. Operation Nest Egg started in the mid 1990’s and was one of the earliest intensive endangered species breeding augmentation programmes globally.

Many of the pioneering techniques and approaches used in kiwi ONE have been replicated, adapted for, or have inspired, successful *ex-situ* / *in-situ* augmentation programs in other species in Aotearoa and internationally, for example, kākāpo (DoC 2018), Tasmanian devils (Hogg et al. 2019), and even zebra sharks (Traylor-Holzer 2021).

Brown kiwi (*Apteryx mantelli*) in the North Island, and rowi or Ōkārito brown kiwi (*A. rowi*), tokoeka (*A. australis* “Haast”), and Roroa/Great Spotted kiwi (*A. haastii*) in the South Island are part of ONE. Kiwi are threatened in Aotearoa New Zealand primarily because of introduced mammalian predators like cats, dogs, and mustelids. Introduced stoats (*Mustela erminea*) are a particular problem for kiwi as they target juvenile birds (McLennan et al. 1996, 2004; Murphy et al. 2008). The current population of all kiwi species combined is approximately 68,000, and kiwi populations not under any conservation management are still declining at least 2% per annum (Germano et al. 2018). In these unmanaged populations, with no predator control or ONE, up to 95% of juvenile kiwi are predated (McLennan et al. 1996).

Operation Nest Egg works by removing young kiwi, while they are fertile eggs, from within the nests of monitored sires. The eggs are then incubated and hatched in captive facilities. Hatchling chicks are hand-raised for approximately one month in indoor brooders, and then released outside into semi- or fully captive predator free crèches. Juvenile kiwi are returned to the wild, usually to their parent’s forest (though translocations often occur [Jahn et al. 2022]), when they reach a “predator proof” weight of approximately 1 kg (Colbourne et al. 2005; Bassett 2012; Colbourne et al. 2020). Programme ONE is considered acceptable from a behavioural point of view as kiwi are precocial, with limited (Craig et al. 2023) or no parental care, and juveniles can survive post-release using instinct (Bassett 2012; Colbourne et al. 2020).

As part of ONE, data pertaining to husbandry and life history is kept by zoo keepers. Each breeding season data collected includes sire identification, egg size, egg clutch number and egg number from within a clutch. Whether the embryo is malpositioned and required an assisted hatch is recorded, as is hatch weight, hatch date, chick daily weight change, food intake, illness incidence, veterinary interventions, and chick behaviour (Bassett 2012). Multiple centres across Aotearoa conduct ONE, and at least two, the National Kiwi Hatchery (NKH) in Rotorua, and the West Coast Wildlife Centre (WCWC) in Franz Josef, use near-identical incubation and husbandry protocols adapted from Basset (2012). The NKH incubates and hatches Brown kiwi from genetically distinct and formally recognised North Island *A. mantelli* genetic groups (Burbidge et al. 2003; Germano et al. 2018; White et al. 2018), including birds of “Eastern”, “Western”, and “Coromandel” provenance. The WCWC has, until recently, hatched two separate South Island species, rowi and Haast tokoeka. Operation Nest Egg institutions own decade’s worth of hand-written ONE records that have been infrequently examined scientifically, and only in a limited way (Gray 2011, Prier et al. 2013, Vieco-Galvez et al. 2021).

During the 2016/2017 and 2017/2018 breeding seasons, a new kiwi maintenance diet was introduced at both NKH and WCWC. This “new diet” recipe was formulated by the Australasian Zoo and Aquarium Association (ZAA) and Massey University (Barlow 2018) and replaced the “old diet” recipe that was used by NKH and WCWC for feeding chicks and adults. The new diet was developed by bird nutrition experts, and was intended as a maintenance diet for adults, to improve their health outcomes. In this study we take advantage of the congruence in husbandry methods and data recorded from both facilities, and the timing of the introduction of the new diet, to analyse hatchling kiwi chick responses to the “old” vs. “new” kiwi diets. We have extracted ONE data from over 300 chick’s handwritten records, and use these to examine chick food intake and growth. We review the nutritional suitability of each diet for chicks, and determine whether the responses differ among *A. mantelli* populations “Eastern”, “Western”, and “Coromandel”, and *A. rowi* and *A. australis* “Haast”.

The use of ONE, and associated chick translocation, are planned to continue for the next five to ten years (Kiwis for kiwi 2016; Undin et al. 2021; Jahn et al. 2022), especially in *A. mantelli* populations (Kiwis for kiwi 2016). Quantitative analysis of chick captive nutrition and the new ZAA diet is therefore timely and important (Germano et al. 2018; Barlow 2018). It is well established that early life experiences (pre- and post-natal), especially nutritional experiences, impact adult physiology and fitness, including trans-generationally (Nettle et al. 2013; Gruber et al. 2018; Aristizabal et al. 2020). Ensuring that the captive husbandry methods we use as keepers best supports kiwi survival in the wild post-release is critical.

The comprehensiveness of the ONE data collected also allowed for preliminary investigations of fundamental kiwi egg and chick life history characteristics. Kiwi exhibit extreme evolutionary and ecological novelty (Calder 1979; Reid & Williams 1975) and have an unusual breeding strategy that involves burrow nesting, laying proportionally enormous eggs for their body size, and providing very limited parental care (Castro 2011). The evolutionary context for this strategy is still poorly understood, but being secretive and nocturnal, kiwi are difficult to study in nature, limiting research progress (Cunningham & Castro 2011). Examining ONE data and relating these to other bird species life history strategies could help us to better understand *Apteryx* adaptations. Here we examine whether genetic groups differ in their mass loss prior to first feeding, and whether egg size and hatchling size differ between: genetic groups; clutches; within clutches; and between the sexes. We also investigate whether certain kiwi populations and ONE sires are more prone to producing “problem” offspring that require more intensive captive management, and how all of these responses may interact with rearing diet.

This our work demonstrates that zoo-based hand rearing records are a largely up-tapped, yet rich scientific resource for improving the captive care and our basic knowledge of young kiwi.

## METHODS

### Genetic groups

We extracted data from 196 Brown kiwi chick (*A. mantelli*) records raised at NKH from across the 2007/2008, 2008/2009, 2009/2010, 2015/2016, 2017/2018, and 2018/2019 breeding seasons. Records were from the following North Island *A. mantelli* genetic groups: “Coromandel” (*n* = 52 records extracted), “Eastern” (*n* = 79), and “Western” (*n* = 65). Coromandel chicks were sourced as eggs from forest on the Kūoatunu Penisula. Eastern eggs came from forests in Omataroa, Maungataniwha, and Ōhope, and Western eggs were collected from Tongariro and Taranaki forests (Figure 1).

**Figure 1.**
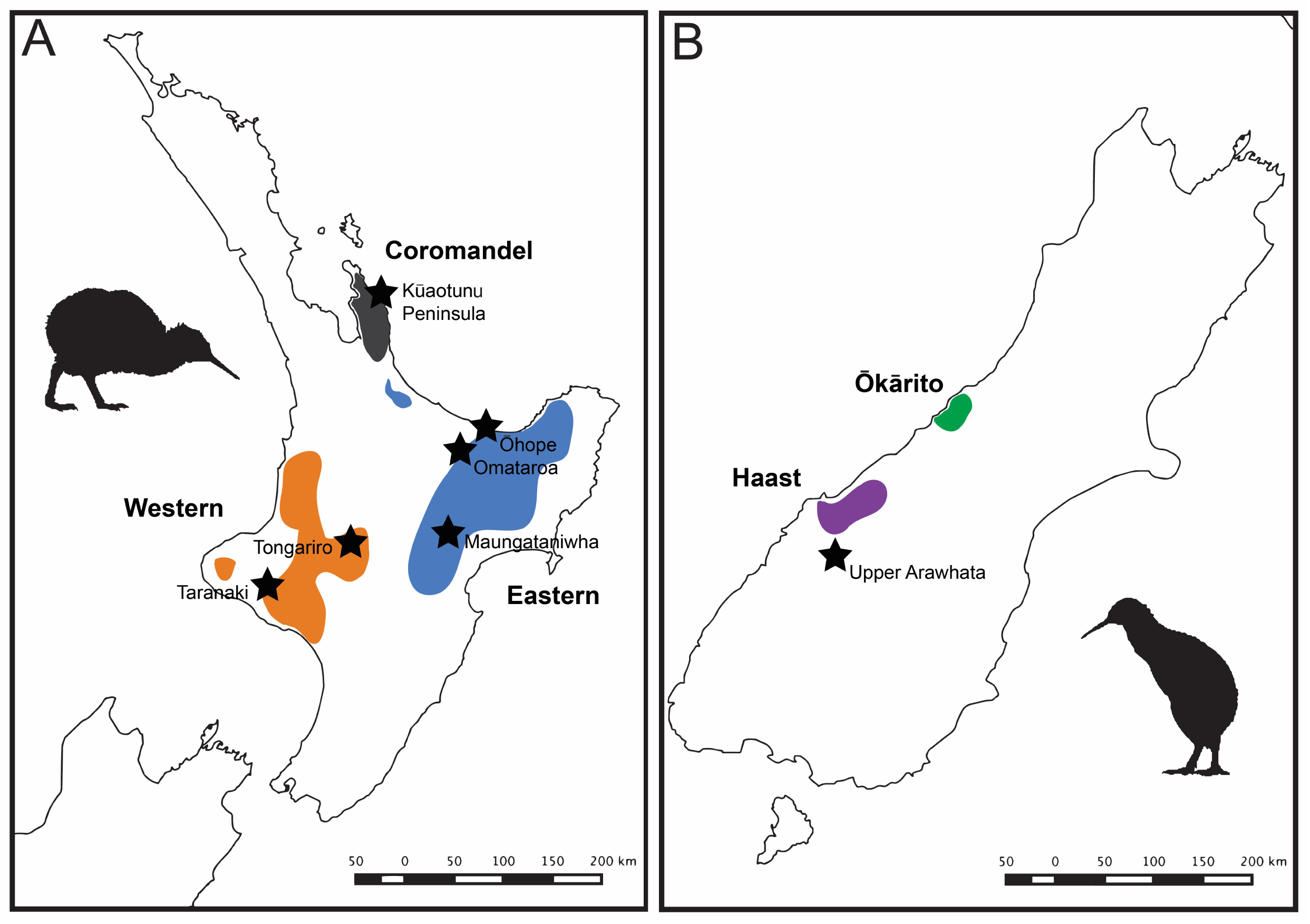
Maps of the North Island (A) and South Island (B) of Aotearoa/New Zealand showing where the Operation Nest Egg (ONE) chicks from this study were collected as eggs.

**Figure 2.**
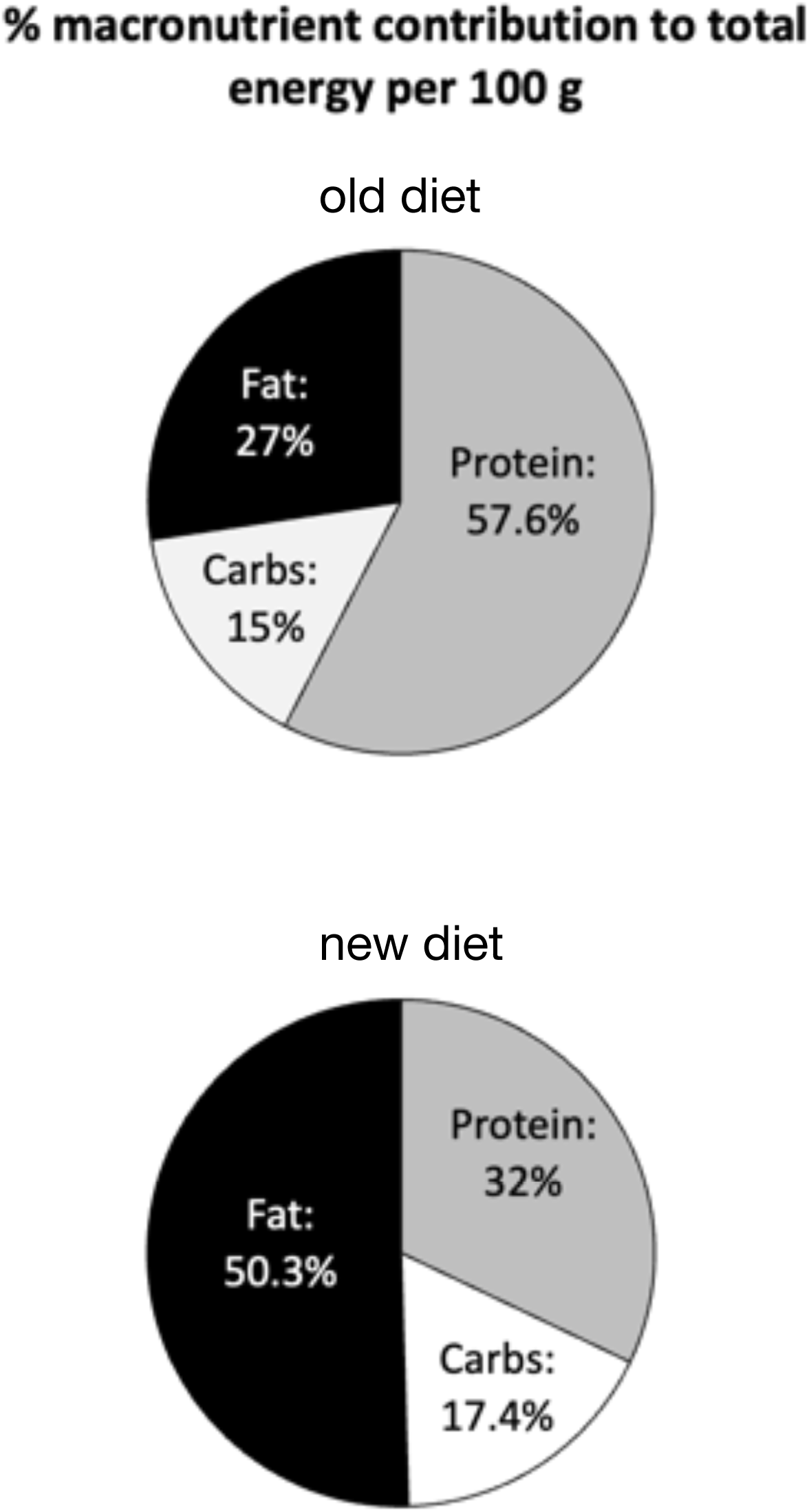
The percentage of energy provided by each macronutrient in the old (upper pie chart) and new Massey ZAA (lower pie chart) kiwi maintenance diets. Note that the majority of old diet’s energy comes from dietary protein, while fat supplies most of the energy to kiwi chicks feeding on the new diet (also see Table 2).

From the WCWC we extracted data from 105 records from *A. rowi* (*n* = 57) and *A. australis* “Haast” (*n* = 48) from across the 2012/2013, 2013/2014, 2015/2016, 2016/2017, 2017/2018, and 2019/2020 seasons. Ōkārito brown/rowi chicks were collected as eggs from the Ōkārito forest, and Haast eggs from both the Haast forest and the upper reaches of the Arawhata river on the West Coast of the South Island (Figure 1). Extracted records, including the specific collection forests of individual chicks and their parentage, are available in data supplement SM1.

### Operation Nest Egg rearing protocol

Kiwi eggs and chicks were raised at NKH and WCWC following Bassett (2012). Minor variations in these methods occur when rearing Haast tokoeka and rowi/ Ōkārito kiwi. For Haast tokoeka, incubation relative humidity (RH) is maintained at ∼45% until the chick externally pips during hatch, whereas other taxa eggs are incubated at 65 – 75% RH until external pip. Haast tokoeka are also offered food for the first time over-night on day 7 post hatch, rather than over-night on day 6 for Brown kiwi, and overnight on day 5 for rowi. Otherwise, the rearing protocols and data recorded at WCWC and NKH are the same.

Briefly, eggs with viable embryos were sanitised in “Incusan” alkyl benzyl dimethyl ammonium chloride solution (Chook Manor, New Zealand) to remove micro-organisms, and then artificially incubated following Bassett (2012). Embryonic development and chicks’ progress through to hatching were monitored using egg candling. Candling, and other regular viability checks (like embryonic movement checks) identify malpositioned embryos, allowing zoo keepers to “assist-hatch” chicks if required. Kiwi keepers are non-interventionist, however, due to the threatened status of kiwi and the challenging ethics of not assisting, malpositioned chicks struggling to hatch are assisted either through: the keeper manually internally-pipping the membranes for the chick; placing lateral cracks on either side of the egg shell to assist the chick to complete its hatch; or, fully-assisting the chick to hatch by completely peeling off the eggshell. Having performed any of these procedures is considered an assisted hatch in this study. Most chicks hatched un-assisted.

Two days after hatch, chicks were moved from individual hatchers into wooden brooder boxes which have a ∼ 1200 x 600 mm run area and a ∼ 400 x 600 mm bed-box. Brooder boxes were kept together in temperate controlled brooder rooms maintained at ∼18°C and at ambient humidity. Overhead florescent lights were turned on at 8:00 AM and off at 4:00 PM. The NKH also has windows that allow natural light in, while WCWC does not. Bed-boxes were fitted with ceramic heat-lamps set at 27°C that are gradually reduced to room temperate and turned off on day 6. A shallow, wide dish of tap water was provided in the run area and changed daily. The run area was covered in moisten, composted peat-moss, filled to approximately 100 mm depth (just deeper than “bill depth”) for chicks to probe in. Peat was dug-over daily and food-scraps and faeces removed. Chicks were weighed daily in the mornings, and their weight, and notes on health and behaviour recorded.

When food was first offered for overnight self-feeding, 10 g was placed in a plastic dish in the run area before lights off. If chicks did not eat this food voluntarily, on the morning of day 6 for rowi, day 7 for Brown kiwi, and day 8 for Haast tokoeka, 2 g of diet was offered and hand-fed to the chicks as an “introduction” to the captive diet. Many chicks commenced self-feeding after the introduction feed and are offered initially 5 g and then 10 g of food more than they ate the previous night, until a maximum of ∼70 g was offered overnight. Chicks that failed to self-feed after the introductory feed were given an assist-feed by hand each morning, which increased 2 g / day. Chicks who failed to self-feed after 10 days were given an additional hand-feed around noon daily. These second feeds commenced at 4 g and increase by 2 g / day until the chicks began to eat independently. Most chicks were self-feeding by day 10. The total amount of food eaten each day was recorded. In most cases, once chicks were established on the artificial diet/s and had re-gained their hatch weight, they were released from brooders to outdoor crèches. Please note, these rearing methods were correct as of 2019 and have since been updated to reflect best-practice. Contact the ONE Best Practice Group for current methodology.

### Diets

Both diets were prepared in the same way, and their recipes are shown in Table 1. All meat was pre-trimmed and kept frozen until needing to be used, when it was defrosted over-night in a refrigerator. Portions of wheat germ, oats, cat food or currants were blanched in boiling water, and then set aside to cool to room temperature. Frozen corn, peas and carrots were thawed at room temperature prior to use. Fresh broccoli, apple, pear, banana, zucchini and silver beat were used. All fruit and vegetables were blended to small (∼2-8 mm) pieces using a standard domestic food processor. A proprietary insectivore vitamin mix “Kiwi Pre-mix” (Vetpak, New Zealand) was added to both diets at the rate of 2 g/100 g of food (see Table S1 for pre-mix composition). For the new diet only, the oils and calcium carbonate powder where blended together into a smooth paste. All prepared ingredients were then thoroughly mixed to a uniform consistency using a large clean spoon, spatula or by hand. Diet was fed out to chicks fresh or following a maximum of 24 h refrigeration.

**Table 1.** The old kiwi maintenance diet recipe (left), based on ox-heart, used until recently by the West Coast Wildlife Centre, Franz Josef, and the National Kiwi Hatchery, Rotorua. The new Massey ZAA kiwi maintenance diet recipe (right) now used across at kiwi hatcheries in New Zealand since 2016.

| *Old Kiwi Diet*<br>Ingredient | Amount (g/100 g) | *New ZAA diet*<br>Ingredient | Amount (g/100 g) |
| --- | --- | --- | --- |
| Oxheart | 78 | Lean beef (fat trimmed) | 40.5 |
| Rolled oats | 2.5 | Oxheart | 13 |
| Wheat germ | 0.78 | Wheat germ | 1 |
| Cat biscuits (Hillspet) | 3.25 | Banana | 7 |
| Apple | 0.78 | Apple | 3.5 |
| Banana (skin removed) | 2.6 | Pear | 3.5 |
| Broccoli (florets) | 1.82 | Carrot | 7 |
| Zucchini | 0.78 | Cat biscuits (Hillspet) | 7 |
| Silver beet (leaf only) | 0.39 | Currants | 4 |
| Carrot (peeled) | 3.03 | Peas | 3 |
| Corn | 3.03 | Corn | 5 |
| Peas | 3.03 | Canola oil | 1 |
| - | - | Corn oil | 1 |
| - | - | Calcium Carbonate<br>(powder) | 1.5 |

Following Gosby et al. (2010). We determined the nutritional composition of the kiwi diets by retrieving each ingredient’s nutrient profile from the Food Standards Australia New Zealand Food Composition Database (FSANZ 2021). For the proprietary cat biscuits, we used the nutritional table provided on the manufacturer’s website (see: https://www.hillspet.co.nz/cat-food/sd-feline-adult-perfect-weigh-dry).

The new ZAA kiwi diet was introduced at NKH during the 2017/2018 season. The new diet was introduced during the 2016/2017 season at WCWC. Below we compare the nutrient composition of these two artificial diets to the best estimated wild kiwi diet, as reported in Potter et al. (2010).

### Data extraction

We aimed to extract records from at least 10 female and 10 male chicks reared on each diet from each genetic group. We avoided extracting records from multiple clutches from the same sire, though this was not always possible. The ONE programme involving *A. rowi* and *A. australis* does not always determine chick sex, so we did not conduct sex based comparisons for these species. For these taxa, we extracted as many records as we could in the time we had available.

Hand-rearing data were transcribed to Microsoft Excel, and were: chick ID, chick genetic provenance/taxa, breeding season, sire, clutch number (from within a breeding season), egg number (from within a clutch), egg length and width (at longest and widest parts of the egg, respectively), whether the embryo was malpositioned, hatch mass, hatch date, hatch type (normal *vs.* assisted hatch), chick sex, chick mass (per day), diet type (“old” or “new”), and food intake. Food intake was separated into the amount self-fed per day and the amount assist-fed/or hand-fed per day.

Sometimes, at the peak of hatching season, two chicks the same age were housed together in the same brooder. In these instances, we averaged the self-feeding daily food intake recorded for the brooder across both resident chicks. We also recorded the date they were released from the brooder room into a juvenile crèche. Please refer to SM1 for a copy of the extracted records.

### Statistical comparisons and data analysis Relative growth efficiency on the new and old diets

The most intensive part of ONE ends the day chicks are released from their brooders. It is a major ONE milestone. Here we have assessed how well each diet (new and old) supported this milestone’s attainment by evaluating chick growth efficiency on each diet from the day they commenced feeding to the day they were released. This time variable is referred to as “days” below, for “days to release”.

We used two separate generalised linear mixed models (GLMMs), using R (ver. 3.4.0) package lme4 (ver. 1.1-15) to evaluate growth efficiency, and in subsequent GLMMs. One GLMM compared relative performance among the five kiwi genetic groups on the new diet. The second compared relative performance on the old diet. The response variable was a daily growth index. The index was determined as the proportional daily mass change per unit food intake. Proportional daily mass was calculated by dividing a chick’s mass, on a given day, by the chick’s hatch weight. This ensured the chick’s daily mass was always proportional to their starting mass (hatch mass). We did this to standardise mass changes between and among chicks of different starting size. To calculate the daily growth index, the proportional daily mass for a chick on a given day was divided by the chick’s food intake on that day, Growth index values decreased over time. This was because the chicks’ mass changes became smaller over time as their food intake stabilised, and their growth per day slowed. To improve our GLMM interpretation, we inverted the growth index (by calculating: 1/the growth index value). This allowed us to interpret index values that increased over time, which we found more intuitive. Note, inverting the index did not impact the meaning of our analysis, as the gradient of our GLMM slopes, and a slopes’ position relative to one another were not altered. As kiwi chicks gain relatively more mass per unit food intake in the first week of feeding, the index distribution was highly right-skewed. We log-transformed the growth index to normalise the distribution before the conducting GLMMs.

We followed model specification methods presented in Harrison et al. (2018). We used the growth index as the response variable, regressed against “days”. To investigate the relationship between how long the chicks spent in the brooder room, their genetic background, and their growth efficiency on each diet, we included the interaction term of chicks’ genetic group and “days” in the models. The model was specified as: growth efficiency ∼ genetic group*days (1 | days). Time (days) was fitted as a random factor, though with random intercepts only, not random slopes. This was due to insufficient degrees of freedom, and previous models failing to converge with days fitted fully randomly. This model fitting still allowed for the intercept position of each variable’s slope to differ (see Harrison 2018), allowing us to determine the relative growth efficiency of chick genetic groups.

To communicate the chick daily growth index visually in our figures we plotted individual chick’s daily food intake (cumulative) on the *x*-axis and their proportional mass change for each day on the *y*-axis. Individual points on these plots provide a visual representation of the growth index values for each chick on each day while still allowing total food intake and mass change to be visualised. A log-scale was used on the *x*-axis.

Next, we tested for within genetic group differences in the growth efficiency of chicks raised on the new *vs*. old diets. We used individual GLMMs for each genetic group and used the same growth efficiency index as above. Models were specified as: growth efficiency ∼ diet*days (1 | days). We used a Bonferroni corrected alpha of 0.01 (α =0.05/5 comparisons) for these tests.

### Sex and diet differences in growth efficiency in Brown kiwi

Among the *A. mantelli* genetic populations, we tested whether there where sex related differences in growth efficiency on the new or old diet. The response variable was the growth index described above, and we used the same modelling approach.

### Mass loss over the first eight days

We also tested whether the kiwi chick genetic groups and/or sexes differed in how much mass chicks lost over the first eight days after hatching. Mass loss is largely due to yolk absorption, and by day eight most chicks have commenced self-feeding and ceased losing mass (Prinziner & Dietz 2002). The first GLMM tested whether chick proportional daily mass differed over time due to chick genetic background. We included the interaction term between days and genetic group in the model. The next GLMM analysed Brown kiwi (*A. mantelli*) populations only (i.e., Eastern, Western and Coromandel) and determined whether sex, genetic group, and/or time (days), and their interaction significantly influenced mass loss.

### Relationship between hatchling mass, egg size and release day

We used general linear models (GLMs) to determine whether there was a significant relationship between the total number of days it took a chick to be released from the brooder room and the chick’s hatch mass. First, we tested whether there was a significant relationship between time (i.e., the number of days to release from the brooder) and hatch mass, rearing diet, and interactions hatch mass*diet, and hatch mass*diet*genetic group. The second GLM analysed data from Brown kiwi only, and examined whether sex interacted with hatch mass, diet and genetic background. We then repeated these analyses for egg size. Egg size was estimated as egg length + width.

### Days to regain hatch mass: developmental time

In ONE, regaining hatch mass indicates a chick has fully completed yolk-absorption, is self-feeding and is ready for release. In the wild, this is likely to be the time the chick commences dispersal, or longer nocturnal foraging trips. We have analysed how long it took chicks from each population to regain hatch mass to determine whether there might be subtle life history differences between the populations, and whether the timing of this developmental milestone interacted with diet type. We used ANOVA (R “Stats” package ver. 3.4.0). to determine whether *Apteryx* spp. differed in their development time, and whether this interacted with rearing diet. We used multiple post-hoc *t-*tests (with a Bonferroni corrected alpha of 0.005) to identify significant differences between genetic populations and/or diets.

### Relationships between developmental time, assist-hatching, malpositioned embryos and assist feeding

As practitioners, we are occasionally required to intervene to save the life of an individual kiwi. Interventions can include assisting malpostioned embryos to hatch, and hand-feeding chicks who refuse to voluntarily consume the artificial diet. We investigated whether these high-needs ONE chicks also took longer to develop. Using ANOVA we tested whether being an assisted-hatch, a malpositioned embryo, or being assist-fed (effectively force-fed) for more than four consecutive days caused slower development, and whether these interacted with genetic group. To test whether the diet used to hand-feed influenced a high-needs chick’s response to assist-feeding, we conducted an additional ANOVA to test for an interaction between diet type (the old *vs*. new maintenance diets) and developmental time (Bonferroni adjusted alpha of 0.025).

### ***Brown Kiwi (*Apteryx mantelli*) life history data*** Hatch mass & egg size differences due to sex, clutch number, genetic group and egg ***number***

We performed four separate Kruskal-Wallis analyses to test whether hatch mass significantly differed among the three genetic groups of Brown kiwi (*A. mantelli*) due to either chick sex; the clutch number the chick was from (either the 1st or 2nd clutch of a breeding season), egg number (egg one or two from either clutch from any season), or chick genetic group (Eastern, Western or Coromandel). As we conducted four analyses on the same response variable, we used a Bonferroni corrected α of 0.0125. To determine whether genetic groups differed in their hatch mass, we conducted three additional planned, post-hoc Kruskal-Wallis comparisons. We used a Bonferroni corrected α of 0.0071. We used non-parametric methods as sample number and variance were uneven. We repeated these analyses with egg size as the response variable.

### Differences in developmental time due to clutch number and egg number

Using Kruskal-Wallis analyses, we determined whether a chick’s clutch number (either the 1st of the 2nd clutch from any given season) or egg number from within a clutch, influenced their developmental time. We used the number of days it took chicks to regain hatch mass as the response variable. Unfortunately, we did not have sufficient replication to analyse for differences due to genetic group.

### Associations between sex, clutch, and egg number

We conducted a Chi-squared test of independence to determine whether there was a significant association between *A. mantelli* chicks’ sex and their clutch number. We also used a Chisquared analysis to test whether the first or second egg from within a clutch (i.e., egg one or two) was more likely to produce a female or male chick.

### Relationships between genetic groups, sires, Malpositioned embryos, assisted hatching and hand-feeding

Using Chi-squared tests of independence we tested whether Eastern, Western and Coromandel Brown kiwi populations were more likely to produce malpositioned embryos. Next tested whether certain males were associated with siring more malpositioned embryos. We also tested for a relationship between being a malposition embryo and requiring an assisted hatch.

## RESULTS

### Diets

Our FSANZ analyses of diet compositions found the old and new ZAA Massey kiwi maintenance diets differed in their macronutrient composition and concentration. Both diets also differed from the best published estimate of the adult *A. mantelli* wild diet (see Potter et al. 2010) in their macronutrient composition. Here we compare the macronutrient composition of all three.

The protein composition of new and old maintenance diets was similar (16.7 g *vs.* 16.1 g), with the new diet offering only 0.6 g more protein per 100 g than the old diet. The estimated wild diet offers a lot more protein per hundred grams however and is estimated to contain 19.9 g of protein per 100 g (Table 2a). The new kiwi maintenance diet contains over twice as much carbohydrate per 100 g (C = 9.06 g / 100 g) than the old diet (C = 4.2 g / 100 g), and the estimated carbohydrate content of the wild diet (C = 4.13 g / 100 g, Table 2a). It should be noted though that the estimated wild Brown kiwi diet of Potter et al. (2010) did not include plant food sources, so the wild diet carbohydrate estimate is likely artificially low. The new and old diets differed most in their fat content. The fat content of the new diet, at 11.6 g per 100 g, is closer match to the estimated fat content of the wild kiwi diet of 7.19 g per 100 g. The fat content of the old diet was only 3.4 g per 100 g (Table 2a).

**Table 2.** A) The amount (in grams) of each macronutrient available per 100 g in the Massey ZAA new kiwi maintenance diet, the old kiwi diet, and the estimated “wild” kiwi diets. B) The percentage contribution each macronutrient makes to the total energy available in each diet. C) The total energy density (kcal/100 g) of each diet.

| a. Nutrient g per 100 g<br>(wet weight) | Kiwi diet |  |  |
| --- | --- | --- | --- |
|  | new | old | Potter<br>et al.<br>(2010)<br>“wild<br>diet” |
| Protein | 16.7 | 16.1 | 19.9 |
| Carbohydrate | 9.06 | 4.2 | 4.13* |
| Fat | 11.6 | 3.4 | 7.2 |
| b. % each macronutrient<br>contributes to total diet<br>energy | Kiwi diet |  |  |
|  | new | old | Potter<br>et al.<br>(2010)<br>“wild<br>diet” |
| Protein | 32.2 | 57.6 | 49.6 |
| Carbohydrate | 17.47 | 15.03 | 10.2 |
| Fat | 50.33 | 27.37 | 40.2 |
| c. Energy Density (kcal)<br>per 100 g (wet weight) | Kiwi diet |  |  |
|  | new | old | Potter<br>et al.<br>(2010)<br>“wild<br>diet” |
|  | 207.44 | 111.8 | 160.83 |

\*Plant food sources not included in calculations, and the carbohydrate content is likely higher in wild diet.

The macronutrient composition of the artificial diets was fixed per unit intake because the diets were homogeneous mixtures of ingredients. This meant that the blend of macronutrients chicks were able to source energy from was fixed. Protein and carbohydrate each offer 4 kcal of energy per g, while fat offers 9 kcal per hundred g. Protein was by far the most plentiful macronutrient in old kiwi maintenance diet, with over half (57.6%) of the energy available in the old diet derived from protein. Protein supplied 49.6% of the energy in the estimated wild kiwi diet but only 32.2% of the energy in the new kiwi maintenance diet (Table 2b). Carbohydrate supplied a similar amount of energy to chicks feeding on either the new or the old kiwi maintenance diets, providing 17.47% and 15% of total energy respectively. The percentage of total available energy supplied by carbohydrate in the wild diet was 10.2% (Table 2b), though, as stated above, this may be inaccurate. Fat was an important source of energy for both the new maintenance diet and the estimated wild diet, with fat supplying 50.3% of energy of the new diet, and 40.2% of the estimated wild diet. Only 27.4% of the old artificial diet’s energy came from fat (Table 2b).

The diet energy densities also differed. Protein offers fewer kcal per unit intake than fat, and because protein was the major contributor to dietary energy on the old diet (rather than fat), the old diet contained far less energy per unit intake than the new or estimated wild diet. The old kiwi maintenance diet only contained 111.8 kcal per 100 g, while the new diet contains 207.44 kcal per hundred grams. The estimated wild diet for kiwi contains at least 160.83 kcal per hundred grams (Table 2c, note, all values are presented per 100 g wet weight).

The old and new kiwi maintenance diets also differed in their micronutrient composition (Table 3). While there were many differences, we have chosen to only report on, and subsequently discuss, the micronutrients that markedly differ in availability between the two diets and whose possible intake deficits are not likely compensated for by the addition of kiwi pre-mix (please refer Table S1). We would like to future investigate the micro nutritional needs of kiwi, as these remain unquantified.

**Table 3.** Micronutrient content of the new Massey ZAA kiwi maintenance diet compared to the “old” maintenance diet used at the National Kiwi Hatchery and the West Coast Wildlife Centre. Note, a proprietary micronutrient supplement powder called “kiwi pre-mix” was added to all helpings of the old and new kiwi maintenance diets at 2 g / 100 g of prepared diet.

| Nutrient per 100 g | Kiwi diet |  |
| --- | --- | --- |
|  | new | old |
| Calcium (mg) | 33.73 | 8.9011 |
| Copper (mg) | 0.08558 | 0.0161 |
| Fluoride (ug) | 5.6828 | 3.952 |
| Iron (mg) | 1.6498 | 4.1772 |
| Magnesium (mg) | 21.76644 | 21.446 |
| Manganese (mg) | 0.22697 | 0.1658 |
| Phosphorus (mg) | 120.19* | 22.935* |
| Adjusted Phosphorus (mg) | 131.5\$ | 90.79\$ |
| Potassium (mg) | 279.0825 | 272.364 |
| Selenium (ug) | 8.003 | 0.38324 |
| Sodium (mg) | 21.260315 | 3.0531 |
| Sulphur (mg) | 0.02 | NR |
| Zinc (mg) | 2.9826 | 1.7266 |
| Thiamin (B1) (mg) | 0.11545 | 0.42697 |
| Riboflavin (B2) (mg) | 0.25014 | 1.18179 |
| Niacin (B3) (mg) | 2.4355 | 16.7856 |
| Niacin equivalent (mg) | 4.5318 | 8.1672 |
| Pantothenic acid (B5) (mg) | 0.15935 | 0.0295 |
| Pyridoxine (B6) (mg) | 0.11375 | 0.0307 |
| Biotin (B7) (ug) | 0.161 | 0.26195 |
| Folate (ug) | 22.455 | 10.19252 |
| Alpha carotene (ug) | 2.21 | 0.9893 |
| Beta carotene (ug) | 518.85 | 244.218 |
| Cryptoxanthin (ug) | 20.07 | 10.3805 |
| Retinol (ug) (Vitamin A) | 1.3 | 7.8 |
| Retinol equivalent (Vitamin A equivalent)(ug) | 117.65 | 61.5823 |
| Taurine (% dry matter of ingredient) | 0.0224 | 0.0104 |
| Vitamin C (mg) | 2.685 | 3.661 |
| Alpha tocopherol (mg) | 0.4085 | 0.02011533 |
| Vitamin E (mg) | 0.7863 | 0.079637 |
| Malic acid (g) | 0.106 | 0.01976 |
| Tryptophan (mg) | 6.75 | 8.3515 |
| Gamma tocopherol<br>(mg) | 0.32 | 0.007488 |
| Linoleic acid % | 4.7687 | 1.86193 |
| Alpha-linoleic acid % | 2.91 | 1.391351 |

\*It is likely the old diet contains more phosphorus than indicated here. We were unable to find phosphorus content reported for Ox heart by Food Standards Australia and New Zealand (FSANZ). <sup>§</sup> We have used the phosphorus level reported by FSANZ for minced beef, to estimate the phosphorus level of beef/ox heart and to provide the adjusted phosphorus value.

### Relative growth efficiency on the new and old diets Growth on the new diet

The first model compared daily growth efficiency on the new diet (Table 4, and Figure 3). The data from Coromandel chicks was, by default, fitted as the group “Days”, and “Days” represents growth for the Coromandel chicks (i.e., Coromandel*days). The remaining genetic groups’ growth efficiency is interpreted relative to the Coromandel results.

**Figure 3.**
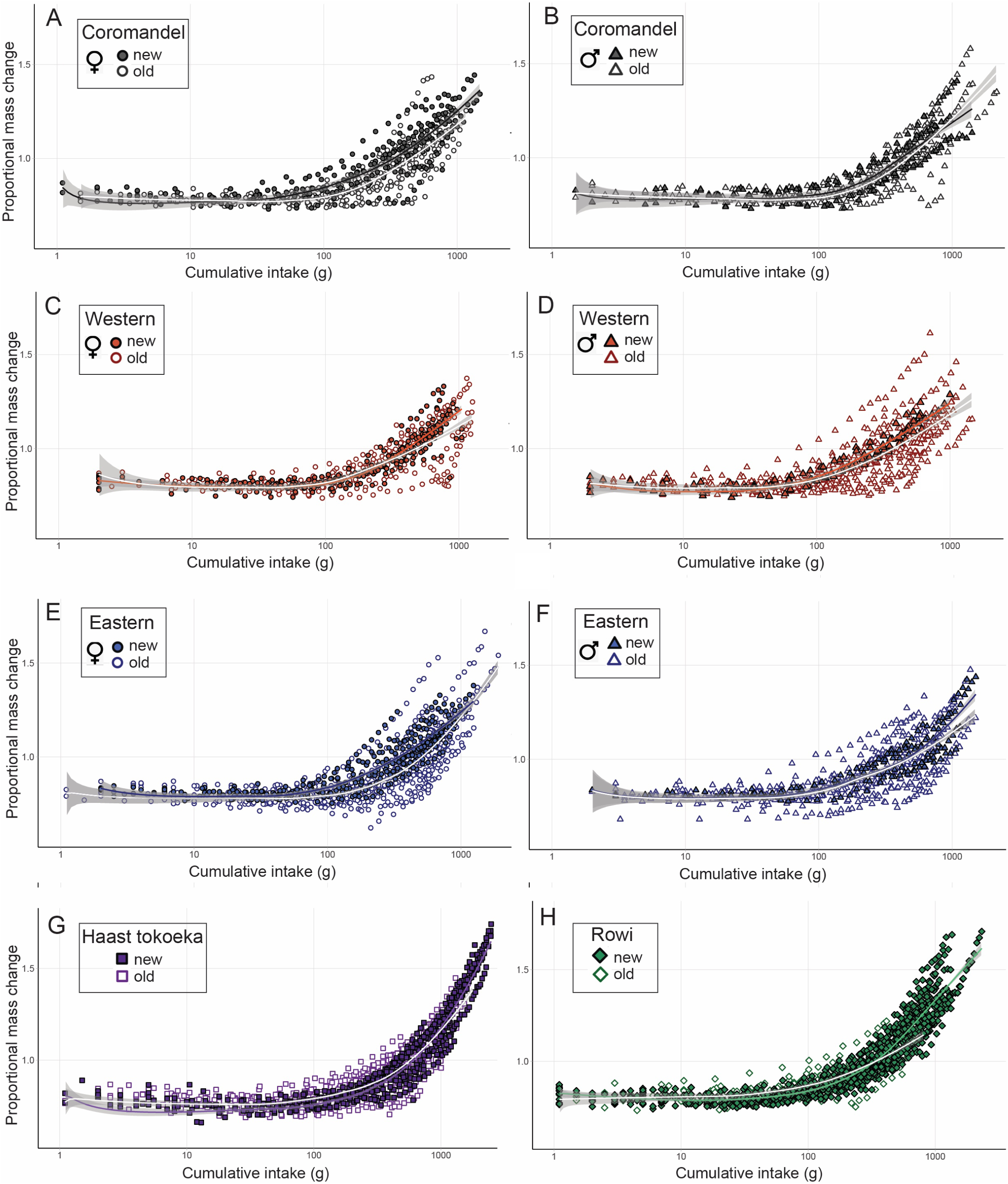
Growth efficiency of kiwi chicks from five genetic groups fed either the new Massey ZAA kiwi maintenance diet (filled symbols and solid colour lines), or the old maintenance diet (open symbols and white lines). Growth efficiency is plotted here as proportional mass change over daily food intake (shown as cumulative). A log scale on the *x* axis was used to allow better visualisation of the food intake data. The lines of best fit are smoothed conditional means ± s.e.m (grey shading). New diet sample numbers: Coromandel *N* = 19 (12 female, 7 male), Eastern *N* = 24 (17 female, 7 male), Western *N =* 20 (13 female, 7 male), Haast *N* = 16, rowi *N* = 19. Old diet: Coromandel *N* = 33 (18 female, 15 male), Eastern *N* = 55 (29 female, 26 male), Western *N* = 45 (17 female, 28 male), Haast *N* = 32, rowi *N* = 38.

**Table 4.**
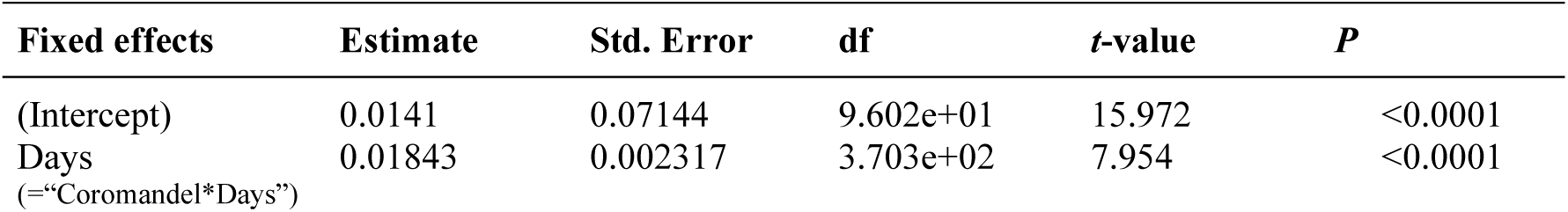
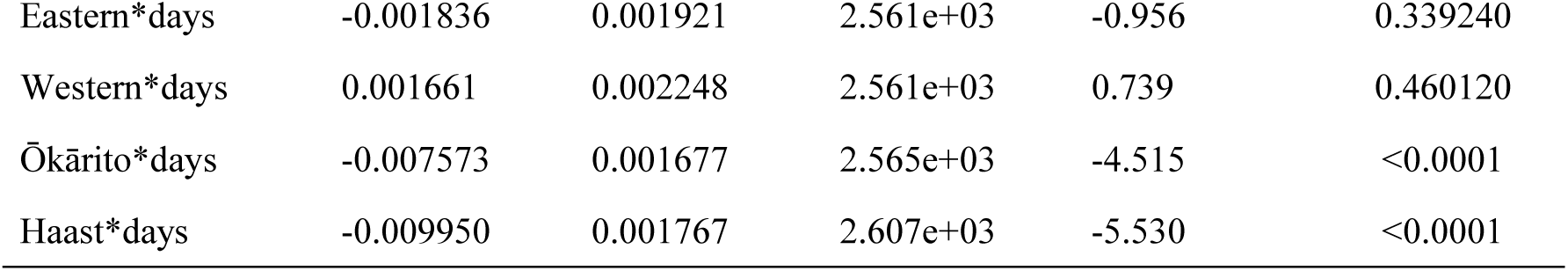
Results from a generalised linear mixed model analysing kiwi chick growth efficiency (mass change per unit daily food intake) for chicks from five different genetic groups raised on the new Massey ZAA kiwi maintenance diet. The negative sign of the Eastern, Ōkārito (rowi) and Haast coefficient estimates indicates these genetic groups grew less efficiently than the Coromandel birds. All the North Island *Apteryx mantelli* genetic groups were statistically equivalent in their growth per day on the new diet, while the South Island species gained significantly less mass per day. Haast chicks grew the least efficiently overall on the new diet.

The Coromandel and Western chick’s growth index variable had a positive coefficient, while the remaining genetic groups had negative coefficients (Table 4). This indicates that on the new diet, Coromandel and Western chicks gained more mass per day, on less food, than the other genetic groups (Figure 3). The GLMM did not detect statistical difference in growth between Western brown chicks and Coromandel chicks (*Western*days*: *t*_(0.00256_) = 0.739, *P* = 0.4601, Table 4). There was also no statistical difference between the Eastern and Coromandel chicks (*Eastern*days*: *t*_(0.00256_) = -0.956, *P* = 0.33924, Table 4), despite the Eastern birds gaining slightly less mass each day when feeding on the new diet than the Coromandel chicks (as indicated by the negative value of the Eastern bird’s beta coefficient, Table 4).

Compared to the Coromandel birds, both the rowi (Ōkārito) and Haast tokoeka chicks grew less efficiently per day feeding on the new diet (*Ōkārito*days*: *t*_(0.002565_) = -4.515, *P* = <0.0001) with the Haast chicks growing the least efficiently on the new diet overall (*Haast*days*: *t*_(0.00261_) = -5.630, *P* <0.0001, Table 4). In other words, the Haast tokoeka grew more slowly and have to eat more over time of the new diet to reach “release” for the brooder room than the other genetic groups.

### Growth on the old diet

The next GLMM investigated chick growth on the old diet. Coromandel grew more efficiently than all other genetic groups when feeding on the old diet (Table 5). The GLMM coefficient estimates indicate that the growth efficiency for the remaining genetic groups was in the following order from most efficient to least: Western browns (*Western*days*: *t*(_0.00346_) = -2.378, *P* =0.0174, Table 5); Eastern browns (*Eastern*days*: *t*_(0.003464_) = -4.895, *P* <0.0001, Table 5); the Haast tokoeka (*Haast*days*: *t*(_0.003479_) = -7.707, *P* <0.0001, Table 5), and lastly the Ōkārito (rowi) chicks (*Ōkārito*days*: *t*_(0.003329_) = -8.621, *P* <0.0001, Table 5).

**Table 5.** Generalised linear mixed model results comparing the growth efficiency of five genetic groups of kiwi chicks raised on the old kiwi maintenance diet. Coromandel chicks grew significantly more efficiently on the old diet than all the other genetic groups. The South Island species grew the least efficiently on the old diet, with the Ōkārito brown kiwi (rowi) gaining the least mass for a unit of food eaten each day.

| Fixed effects | Estimate | Std. Error | df | t-value | P |
| --- | --- | --- | --- | --- | --- |
| (Intercept) | 0.7854 | 0.0965 | 3.604e+01 | 8.101 | <0.0001 |
| Days<br>(=“Coromandel*Days”) | 0.03662 | 0.004194 | 4.247e+01 | 8.732 | <0.0001 |
| Eastern*days | -0.007327 | 0.001497 | 3.464e+03 | -4.895 | 0.000782 |
| Western*days | -0.004148 | 0.001744 | 3.458e+03 | -2.378 | 0.0174 |
| Ōkārito*days | -0.02870 | 0.003329 | 3.479e+03 | -8.621 | <0.0001 |
| Haast*days | -0.01292 | 0.001676 | 3.479e+03 | -7.707 | <0.0001 |

Counter to the results for the new diet, Haast chicks did not grow the least efficiently. Rowi chicks gained the least mass per day per unit intake on the old diet (note the shallowness of the rowi conditional means line of best fit, Figure 3).

Overall, when fed the old diet, there was greater spread among the coefficient estimates for each genetic group than when fed the new diet. This result is visually apparent too (compare open symbols to closed in Figure 3).

### Within genetic group growth efficiency on the new *vs.* old diet

Among the North Island *A. mantelli* chicks, there were no statistical differences between Coromandel chick’s growth efficiency on new versus the old diet (*t*_(0.001064_) = 0.316, *P* = 0.752, Table S2, Figure 3a,b), or Eastern chick’s (*t*(_0.00159_) = -2.474, *P* = 0.0135, Table S4, Figure 3e,f), though Eastern’s approached significant difference. On the new diet the Western chicks grew significantly more each day, on less food (*t*_(0.001986_) = -3.045, *P* = 0.00238, Table S3, Figure 3c,d). For the South

Island species, GLMMs showed the Ōkārito brown chicks also grew more efficiently on the new diet (*t*_(0.00107_) = -3.924, *P* < 0.0001, Table S5, Figure 3h), while the Haast tokoeka GLMM found no statistical difference in chick growth efficiency on the new *vs*. the old diet (*t*_(0.001177_) = -0.9844, *P* = 0.325, Table S6), this was despite some chicks evidently continuing to lose mass following the commencement of feeding on the new diet (see Figure 3g).

Overall, there was a slight visual trend for North Island chicks to grow more per day on the new diet rather than the old, and the reverse being true for the South Island taxa.

### Diet and sex differences in growth efficiency in Brown kiwi (*A. mantelli*)

We found no difference in growth between the sexes of Brown kiwi chicks (*t*_(0.0038_) = -1.271, *P* = 0.204, Table 6), or any significant interactions between the two kiwi diets and sex (*t*_(0.0038_) = 0.177 *P* = 0.8593, Table 6). This was despite the subtle trend for female Coromandel and Eastern chicks to grow more efficiently on the new diet compared to the old (see above and figures 3a & e).

**Table 6.** Results from a generalised linear mixed model testing for sex-based difference in growth efficiency for brown kiwi chicks (*Apteryx mantelli*) feeding on either the old or new kiwi maintenance diet. There were no significant differences due to sex or the interaction of sex and diet.

| Fixed effects | Estimate | Std. Error | df | t-value | P |
| --- | --- | --- | --- | --- | --- |
| (Intercept) | 0.9239 | 0.09098 | 3.319e+01 | 10.155 | <0.0001 |
| Days | 0.03085 | 0.003957 | 3.824e+01 | 7.797 | <0.0001 |
| Diet*days | -0.00357 | 0.001364 | 3.797e+03 | -2.623 | 0.00874 |
| Sex*days | -0.002302 | 0.001812 | 3.793e+03 | -1.271 | 0.20386 |
| Diet*sex*days | -0.000386 | 0.002176 | 3.799e+03 | 0.177 | 0.85927 |

### Mass loss over the first eight days ***Mass loss differences among all genetic groups***

Patterns of mass loss among the kiwi genetic groups were very similar with most chicks losing ∼20% of their hatch mass within eight days post hatch. Overall, Haast tokoeka chicks lost the most mass, and lost mass faster than the other genetic groups. Most had lost 20% of their hatch mass by day six, whereas the other genetic groups did not lose this amount until day eight (Figure 4). The Coromandel chicks, lost the second greatest amount of mass over the eight days (*Coromandel*: *t*_(7.931_) = -15.962, *P* < 0.0001; *Haast*: *t*_(0.00026_) = -4.7, *P* < 0.0001, Table 7). They were followed by the rowi (Ōkārito), which were statistically equivalent in their mass loss to the Coromandel chicks according to GLMM (*t*_(0.00026_) = 1.39, *P* = 0.16533, Table 4). The Eastern, and then Western chicks lost the least mass over the first 8 days (*Eastern*: *t*_(0.0026_) = 3.687, *P* <0.001; *Western*: *t*_(0.0026_) = 3.55, *P* <0.001, Table 7, Figure 4).

**Figure 4.**
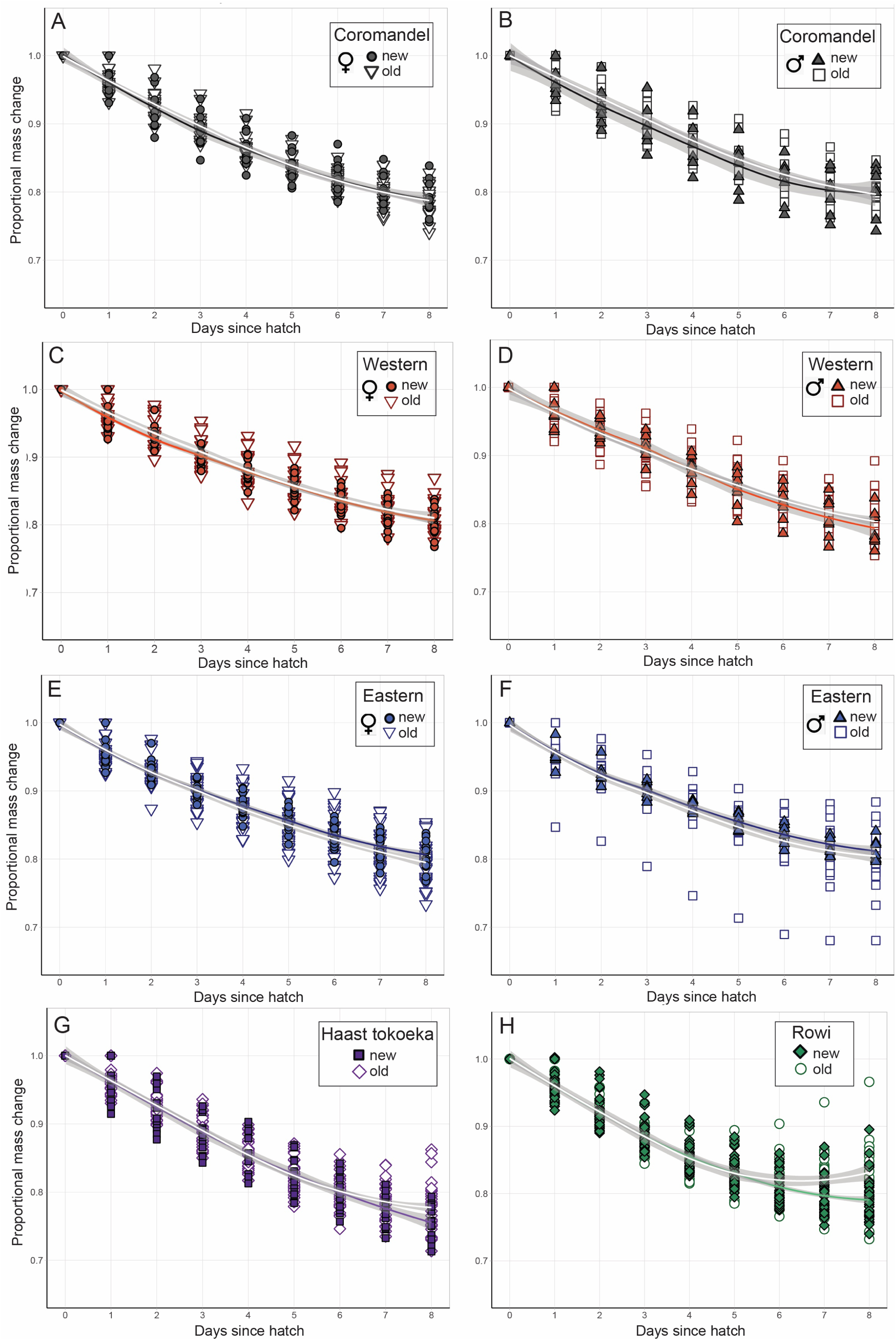
Proportional mass loss in kiwi chicks from five genetic groups over the first eight days of life. While not statistically analysed, the rearing diets used are shown. The new Massey ZAA kiwi maintenance diet is represented by filled symbols and solid colour lines, and the old maintenance diet by open symbols and white lines. The trend lines plotted are for data visualisation only and are smoothed conditional means ± s.e.m. (grey shading). Sample numbers: Coromandel *N* = 54 (31 female, 23 male), Eastern *N* = 79 (45 female, 34 male), Western *N* = 65 (30 female, 35 male), Haast *N* = 48, Ōkārito *N* = 54.

**Table 7.** Generalised linear mixed model analysis determining which kiwi genetic group lost the most mass per day, from hatch to day eight post-hatch. Coromandel and Haast tokoeka chicks lost the most mass over the first eight days of life, with Haast tokoeka chicks losing the most.

| Fixed effects | Estimate | Std. Error | df | t-value | P |
| --- | --- | --- | --- | --- | --- |
| (Intercept) | 0.9858 | 0.007930 | 7.955 | 124.308 | <0.0001 |
| Coromandel*days | -0.02657 | 0.001664 | 7.931 | -15.962 | <0.0001 |
| Eastern*days | 0.002147 | 0.000582 | 2.654e+03 | 3.687 | 0.00023 |
| Western*days | 0.002161 | 0.000608 | 2.654e+03 | 3.550 | 0.00039 |
| Ōkārito*days | 0.000877 | 0.000632 | 2.654e+03 | 1.388 | 0.16533 |
| Haast*days | -0.003024 | 0.0006591 | 2.654e+03 | -4.599 | <0.0001 |

### Mass loss by sex in Brown kiwi

Among *A. mantelli* genetic groups, the GLMM showed no differences between females and males in patterns of mass loss (Table 8, Figure 4).

**Table 8.**
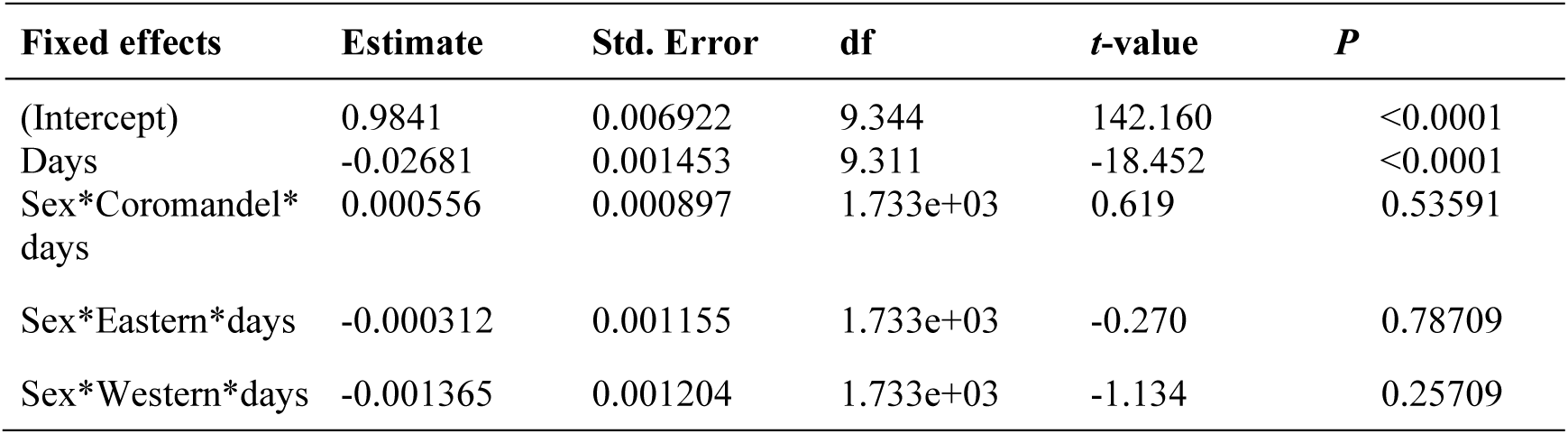
Generalised liner mixed model analysing data from Brown kiwi (*Apteryx mantelli*), demonstrating that there is no difference due to sex and its interaction with genetic group, in the amount of mass lost during the first eight days of life.

### Relationship between hatchling mass and egg size with release day Hatchling mass

We did not identify a relationship between hatch mass and the number of days chicks spent in the brooder room prior to release (*hatch weight*: *t*_(282_) = -0.77, *P* = 0.4403, Table 9, Figure 5). Nor did we identify any significant interactions between the number of days they spent in the brooder room and their hatch mass, genetic group, or rearing diet (see Table 9 and Table 10, all *P* > 0.025).

**Figure 5.**
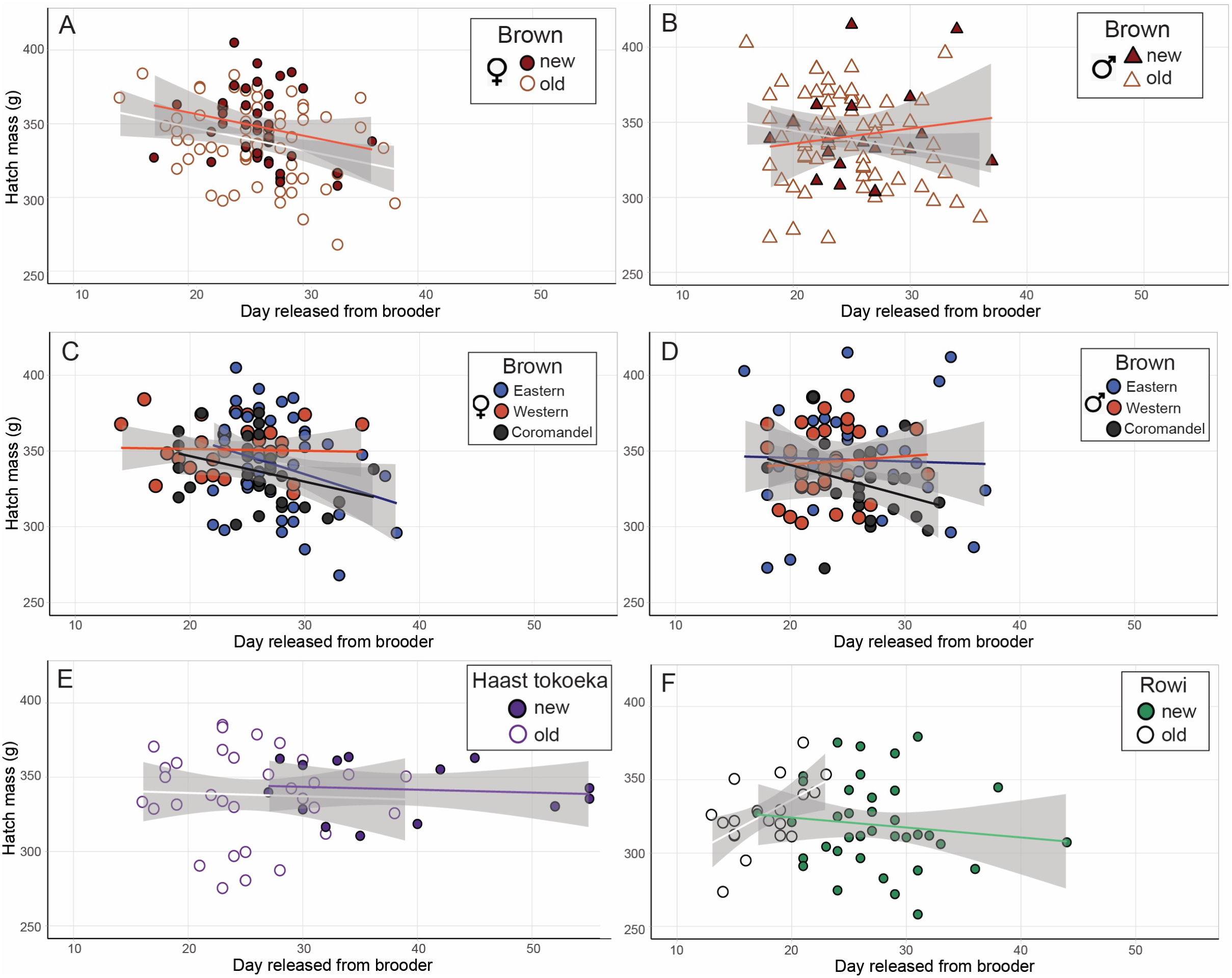
Hatch mass did not significantly influence the number of days it took kiwi chicks of five different genetic groups to reach release day from the brooder room. (New diet = filled symbols and coloured lines; old diet = open symbols and white lines). Trend lines are to assist data visualisation and are smoothed conditional means ± s.e.m. (grey shading). Replicate numbers for new diet: Coromandel *N* = 19 (12 female, 7 male), Eastern *N* = 23 (16 female, 7 male), Western *N* = 20 (13 female, 7 male), Haast *N* = 15, Ōkārito *N* = 38. *N* for old diet: Coromandel *N* = 35 (19 female, 16 male), Eastern *N* = 56 (29 female, 27 male), Western *N* = 45 (17 female, 28 male), Haast *N* = 32, Ōkārito *N* = 19.

**Table 9.** General linear model analysis demonstrating that variation in hatch mass did not significantly influence the number of days it took a kiwi chick to reach release from the brooder room. Hatch mass did not interact significantly with genetic group or diet, in determining days to release either.

| Variable | Estimate | Std. Error | df | t-value | P |
| --- | --- | --- | --- | --- | --- |
| (Intercept) | 44.9327 | 23.3046 | 301 (null) | 1.928 | 0.0549 |
| Hatch weight (h.w.) | -0.05452 | 0.07055 | 282 | -0.773 | 0.4403 |
| Diet | -2.19433 | 26.1030 | 282 | -0.084 | 0.9331 |
| Coromandel*h.w.*diet | 0.00351 | 0.07881 | 282 | 0.045 | 0.9645 |
| Eastern*h.w.*diet | -0.0340 | 0.08947 | 282 | -0.380 | 0.7041 |
| Western*h.w.*diet | -0.06227 | 0.11135 | 282 | -0.559 | 0.5765 |
| Ōkārito*h.w.*diet | 0.08874 | 0.09999 | 282 | 0.887 | 0.3756 |
| Haast*h.w.*diet | 0.09078 | 0.11471 | 282 | 0.791 | 0.4294 |

**Table 10.**
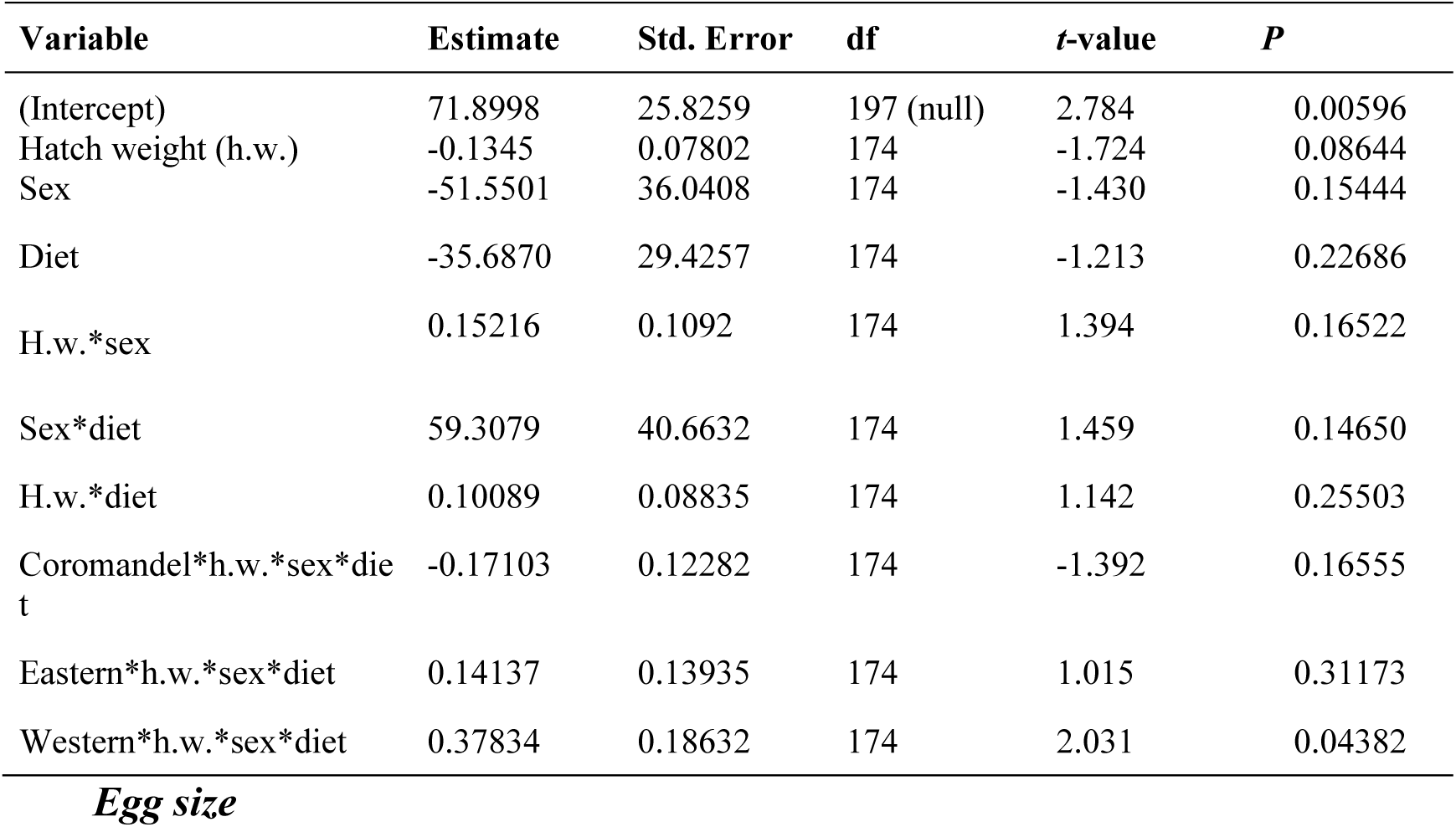
General linear model showing hatch mass in Brown kiwi (*A. mantelli*) does not significantly influence the number of days it takes a kiwi chick to reach release from the brooder room. Hatch mass does not significantly interact with sex, or diet in determining release day.

| Variable | Estimate | Std. Error | df | t-value | P |
| --- | --- | --- | --- | --- | --- |
| (Intercept) | 71.8998 | 25.8259 | 197 (null) | 2.784 | 0.00596 |
| Hatch weight (h.w.) | -0.1345 | 0.07802 | 174 | -1.724 | 0.08644 |
| Sex | -51.5501 | 36.0408 | 174 | -1.430 | 0.15444 |
| Diet | -35.6870 | 29.4257 | 174 | -1.213 | 0.22686 |
| H.w.*sex | 0.15216 | 0.1092 | 174 | 1.394 | 0.16522 |
| Sex*diet | 59.3079 | 40.6632 | 174 | 1.459 | 0.14650 |
| H.w.*diet | 0.10089 | 0.08835 | 174 | 1.142 | 0.25503 |
| Coromandel*h.w.*sex*diet | -0.17103 | 0.12282 | 174 | -1.392 | 0.16555 |
| Eastern*h.w.*sex*diet | 0.14137 | 0.13935 | 174 | 1.015 | 0.31173 |
| Western*h.w.*sex*diet | 0.37834 | 0.18632 | 174 | 2.031 | 0.04382 |

We observed an interesting pattern within the South Island species data (Figure 5, explored empirically below). When raised on the new diet, the day that Haast tokoeka and Ōkārito brown chicks were released from the brooder room occurred much later than when on the old diet. There was also great variability in the South Island species release day when raised on the new diet. This variability was not displayed by South Island species chicks raised on the old diet, and this pattern did not occur among the North Island genetic groups (compare Figure 5a and b to 5e and f). In female *A. mantelli*, there was an apparent trend for heavier hatchlings to be released earlier (Figure 5a & b). However this was not supported statistically by the GLM, which found no differences in hatch mass between the sexes among the North Island Brown kiwi genetic groups (*Hatch weight*sex*: 1.394 : *t*_(174_) = 1.94, *P* = 0.1652, Table 10). While not analysed here (see below), there is also an observable pattern for male *A. mantelli* to have greater variation in their hatch mass than females (Figure 5a & b).

### Egg size

As for hatch mass, there were no significant relationships between kiwi egg size and the number of days it took chicks to be released from the brooder room (*Egg size*: *t*_(238_) = -0.375, *P* = 0.708, Table 11, Figure 6). There were no significant relationships between the number of days it took to be released from the brooder room and egg size, genetic group, diet, or sex (Tables 11 and 12, all variables and interaction *P* > 0.025). As for the hatch mass results, there was a subtle trend for larger female (but not male) *A. mantelli* eggs to take fewer days to be released from the brooder room than smaller eggs (compare Figure 6a to b).

**Figure 6.**
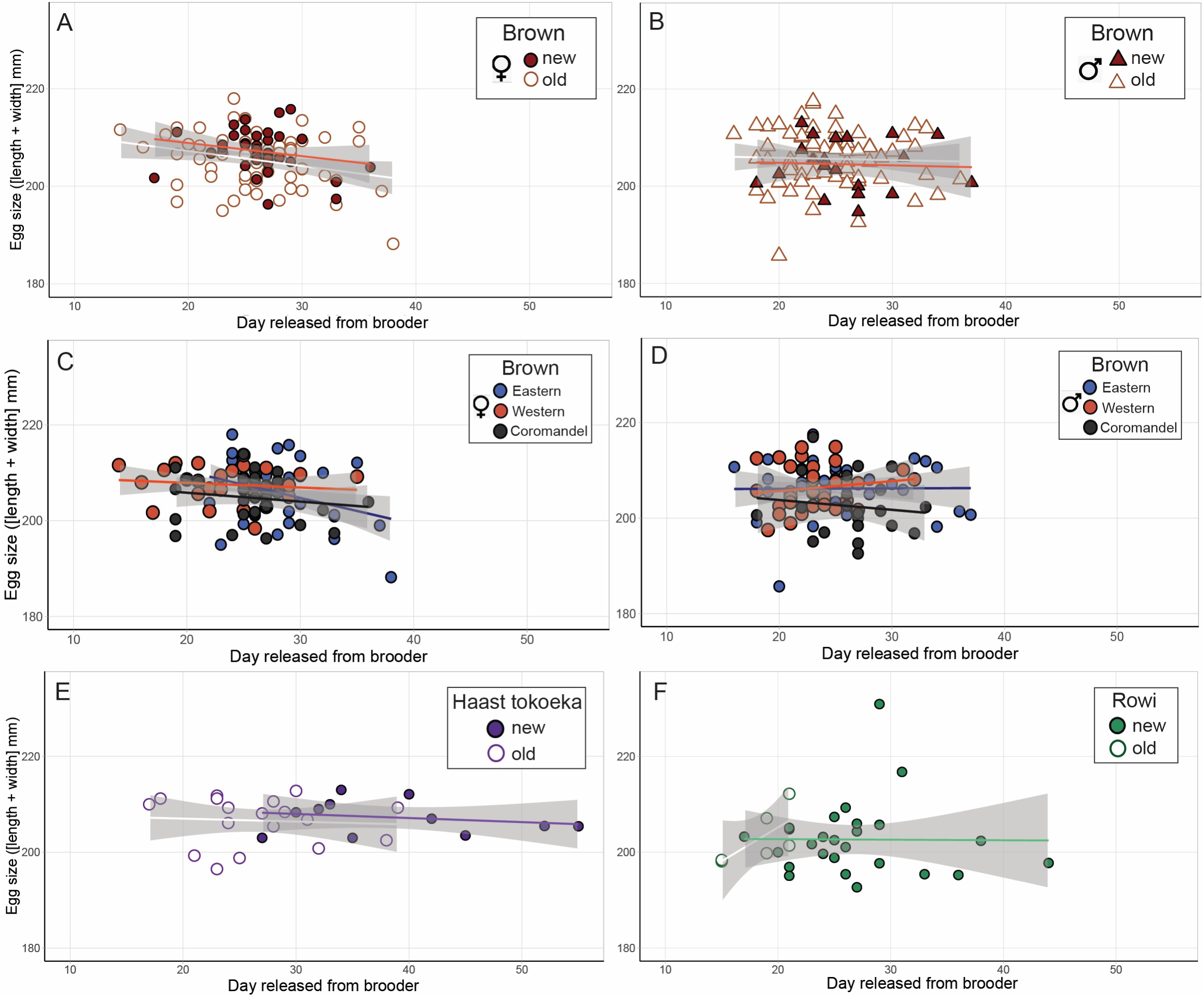
Across five genetic groups of kiwi (*Apteryx* spp.), egg size does not influence how long it takes a chick to reach release from the brooder room. Trend lines are smoothed conditional means ± s.e.m. (shown as grey shading). Chicks reared on the new diet = filled symbols and coloured lines; old diet = open symbols and white lines. Sample *N* for new diet: Coromandel *N* = 18 (11 female, 7 male), Eastern *N* = 23 (16 female, 7 male), Western *N* = 20 (13 female, 7 male), Haast *N* = 12, Ōkārito *N* = 28. Old diet sample *N*: Coromandel *N* = 34 (19 female, 15 male), Eastern *N* = 55 (28 female, 27 male), Western *N* = 44 (16 female, 28 male), Haast *N* = 18, Ōkārito *N* = 6.

**Table 11.** Results from a general liner model showing that egg size does not predict the number of days it takes a kiwi chick to achieve its release day from the brooder room. There were no significant interactions between egg size, genetic group and/or maintenance diet (new *vs*. old) and release day either.

| Variable | Estimate | Std. Error | df | t-value | P |
| --- | --- | --- | --- | --- | --- |
| (Intercept) | 45.47226 | 49.5077 | 257 (null) | 0.918 | 0.359 |
| Egg size (e.s.) | -0.09163 | 0.24404 | 238 | -0.375 | 0.708 |
| Diet | -3.78175 | 58.1851 | 238 | -0.065 | 0.948 |
| Coromandel*e.s.*diet | 0.01251 | 0.28639 | 238 | 0.044 | 0.965 |
| Eastern*e.s.*diet | 0.15891 | 0.43767 | 238 | 0.363 | 0.717 |
| Western*e.s.*diet | -0.15935 | 0.45980 | 232 | -0.347 | 0.729 |
| Ōkārito*e.s.*diet | 0.29591 | 0.52543 | 232 | 0.563 | 0.574 |
| Haast*e.s.*diet | 0.59700 | 0.61394 | 232 | 0.972 | 0.332 |

**Table 12.**
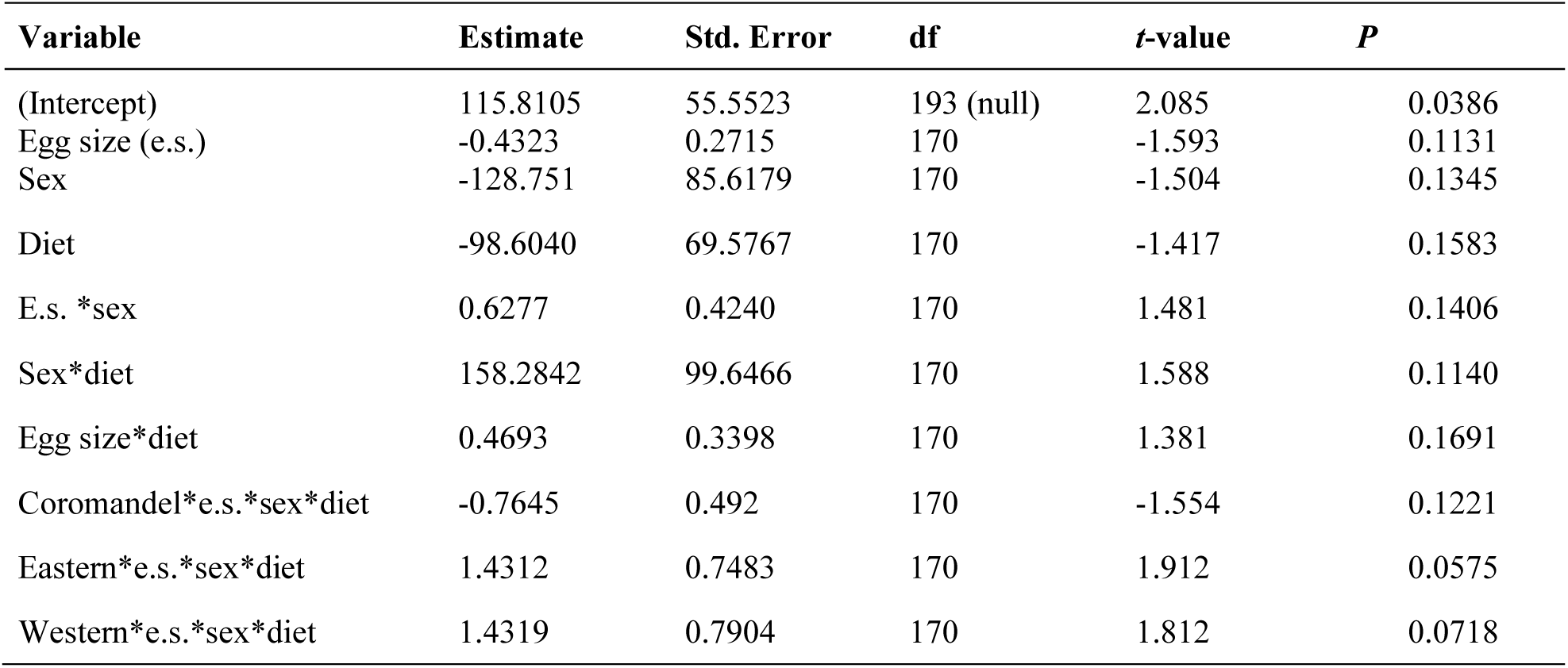
General linear model showing egg size does not influence the number of days it takes *A. mantelli* chicks (Coromandel, Western and Eastern) to reach their release day. Egg size does not interact significantly with sex or diet to influence the time it takes kiwi chicks to be released from the brooder room.

| Variable | Estimate | Std. Error | df | t-value | P |
| --- | --- | --- | --- | --- | --- |
| (Intercept) | 115.8105 | 55.5523 | 193 (null) | 2.085 | 0.0386 |
| Egg size (e.s.) | -0.4323 | 0.2715 | 170 | -1.593 | 0.1131 |
| Sex | -128.751 | 85.6179 | 170 | -1.504 | 0.1345 |
| Diet | -98.6040 | 69.5767 | 170 | -1.417 | 0.1583 |
| E.s. *sex | 0.6277 | 0.4240 | 170 | 1.481 | 0.1406 |
| Sex*diet | 158.2842 | 99.6466 | 170 | 1.588 | 0.1140 |
| Egg size*diet | 0.4693 | 0.3398 | 170 | 1.381 | 0.1691 |
| Coromandel*e.s.*sex*diet | -0.7645 | 0.492 | 170 | -1.554 | 0.1221 |
| Eastern*e.s.*sex*diet | 1.4312 | 0.7483 | 170 | 1.912 | 0.0575 |
| Western*e.s.*sex*diet | 1.4319 | 0.7904 | 170 | 1.812 | 0.0718 |

### Days to regain hatch mass (developmental time) ***Genetic group based differences***

We conducted ANOVA on the number of days it took chicks from each genetic provenance group to regain their hatch mass (previous comparisons between genetic groups where comparing growth efficiency). Genetic groups differed significantly in the number of days it took to regain their hatch mass (*genetic provenance*: *F*(4,289) =18.898, *P* <0.0001, Table 13, Figure 7). Discounting the influence of diet, post-hoc *t*-tests (adjusted α = 0.005) showed rowi (Ōkārito) chicks regained their hatch mass significantly faster than all other genetic groups (Table 14), followed by Western chicks who regained hatch mass significantly faster than Eastern and Haast tokoeka birds (Table 14). There were no significant differences between Coromandel, Eastern, and Haast tokoeka chicks, or between Western and Coromandel chicks (Table 14, Figure 7). Figure 7 shows a general pattern for Ōkārito chicks to regain hatch mass the fastest and Haast tokoeka chicks the slowest, with the *A. mantelli* genetic groups being intermediate.

**Figure 7.**
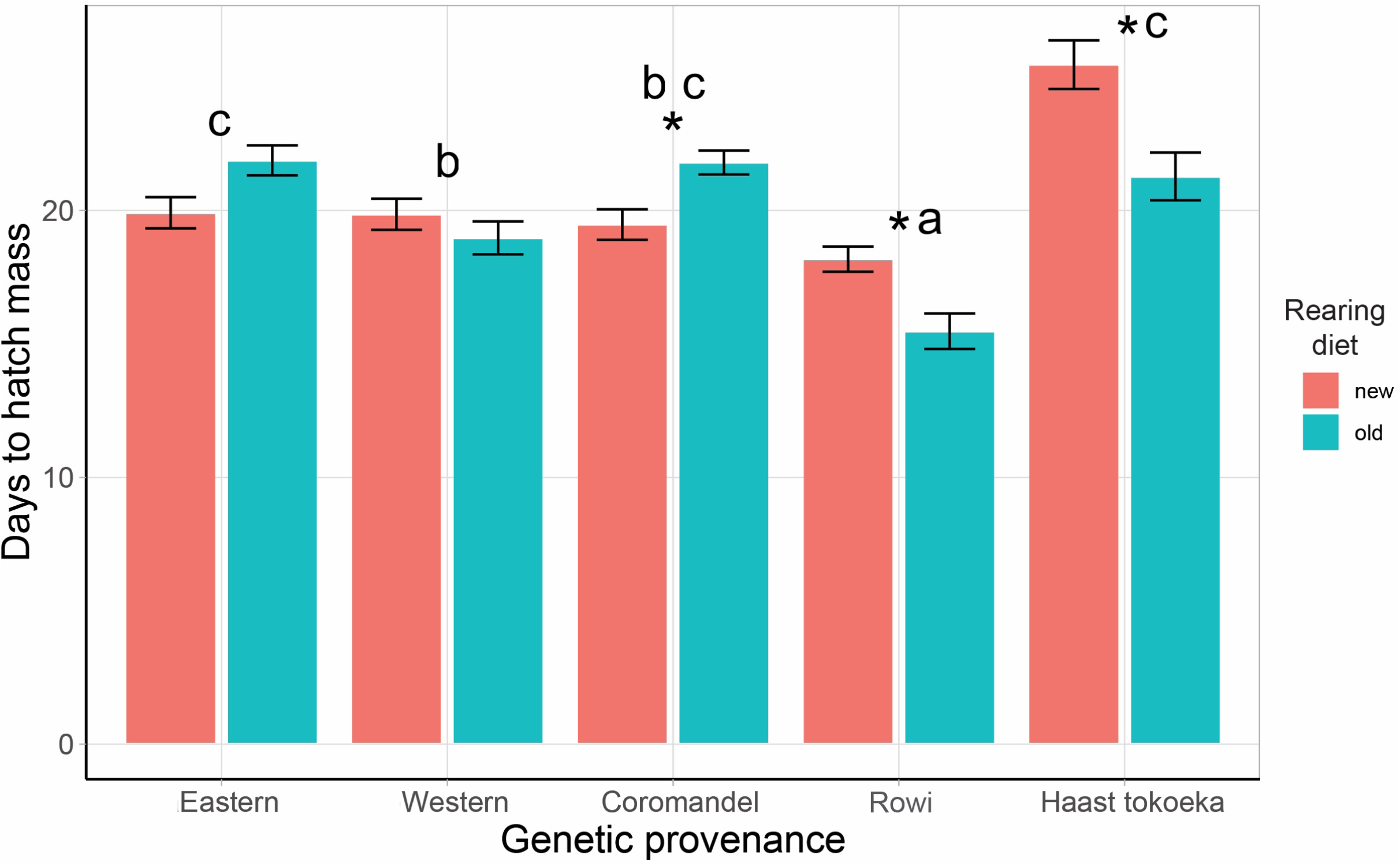
Bar-chart showing the number of days it took kiwi chicks (*Apteryx* spp.) from five genetic groups to regain their hatch mass when reared on either the new Massey ZAA kiwi maintenance diet (orange bars), or the “old” maintenance diet (teal bars). Values are the mean ± s.e.m. Sample *N* for new diet: Coromandel *N* = 19, Eastern *N* = 23, Western *N* = 21, Haast *N* = 15, Ōkārito *N* = 40. *N* for old diet samples: Coromandel *N* = 35, Eastern *N* = 58, Western *N* = 40, Haast *N* = 29, Ōkārito *N* = 19.

**Table 13.** Our analysis of variance showed the five genetic groups of *Apteryx* spp. chicks significantly differed in the number of days it took to regain hatch mass. The ANOVA also showed a significant interaction between rearing diet and the genetic group. This demonstrates that there are differences among the genetic groups in how they respond developmentally to either the “old” or the “new” Massey ZAA kiwi maintenance diets.

| Variable | d.f. | Sum sq | Mean sq | F-value | P |
| --- | --- | --- | --- | --- | --- |
| Diet | 1 | 18 | 18.14 | 1.444 | 0.23 |
| Genetic group | 4 | 949 | 237.36 | 18.898 | <0.0001 |
| Diet*genetic group | 4 | 397 | 99.30 | 7.906 | <0.0001 |
| Residuals | 289 | 3630 | 12.56 |  |  |

**Table 14.**
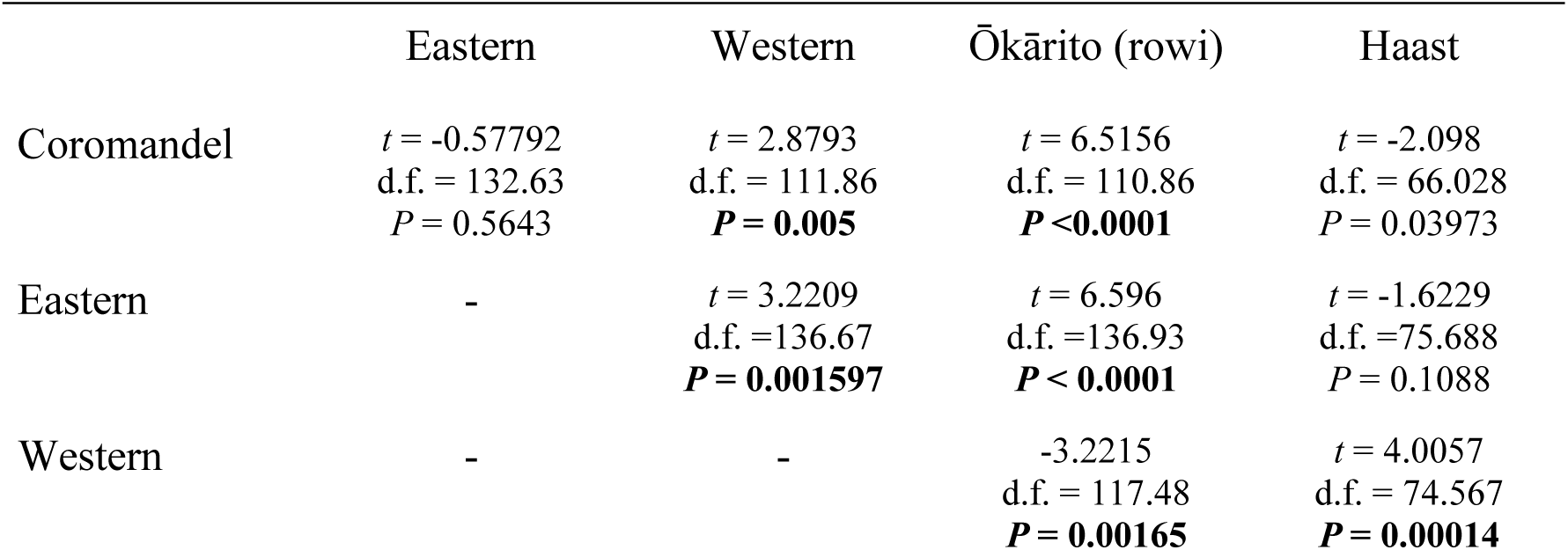
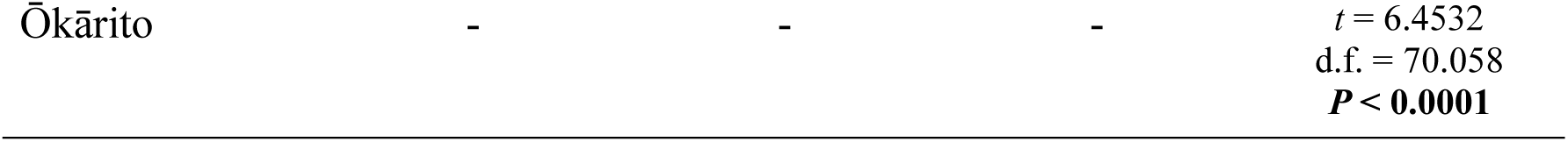
Multiple *t*-tests comparing the number of days it took kiwi chicks from each genetic group to regain their hatch mass (developmental time). We used a Bonferroni corrected alpha of 0.005.

Our ANOVA also demonstrated there were difference among the genetic groups in how long the chicks took to reach hatch mass on the old versus new rearing diets (*genetic provenance*diet*: *F*(4,289) = 7.906, *P* <0.0001, Table 13, Figure 7). Post-hoc within genetic group *t*-tests showed this pattern was driven by the Ōkārito brown, Coromandel and Haast tokoeka chicks. As indicated in Figure 7, Ōkārito and Haast chicks both developed faster on the old kiwi diet (*Ōkārito*: *t*(36.3) = 3.3223, *P* = 0.00204; *Haast*: *t*(36.9) = 3.2825, *P* = 0.00225, Table 14), while Coromandel chicks developed faster on the new diet (*Coromandel*: *t*(39) = -3.1925, *P* = 0.00278, Table 14). There were no statistical differences between the number of days to regain hatch mass on the new or old diets for Eastern or Western *A. mantelli* chicks (*Eastern*: *t*(61.3) = -2.4181, *P* = 0.01859; *Western*: *t*(54.6) = 1.0409, *P* = 0.3025, Table 14).

### Relationships between assist-hatching, malpositioned embryos and assist feeding

We found no differences in developmental time due to being an assist hatch chick or a malpositioned embryo (*assist hatch*: *F*(1,289) = 0.03, *P* = 0.862, Table 15a, Figure 8; *malposition*: *F*(1,289) = 0.278, *P* = 0.278, Table 15b, Figure 9), and neither variable interacted significantly with chick genetic group (*assist hatch*genetic group*: *F*(4,289) = 7.906, *P* = 0.255, Table 15a, Figure 8; *malposition*genetic group*: *F*(4,289) = 0.398, *P* = 0.810, Table 15b, Figure 9).

**Figure 8.**
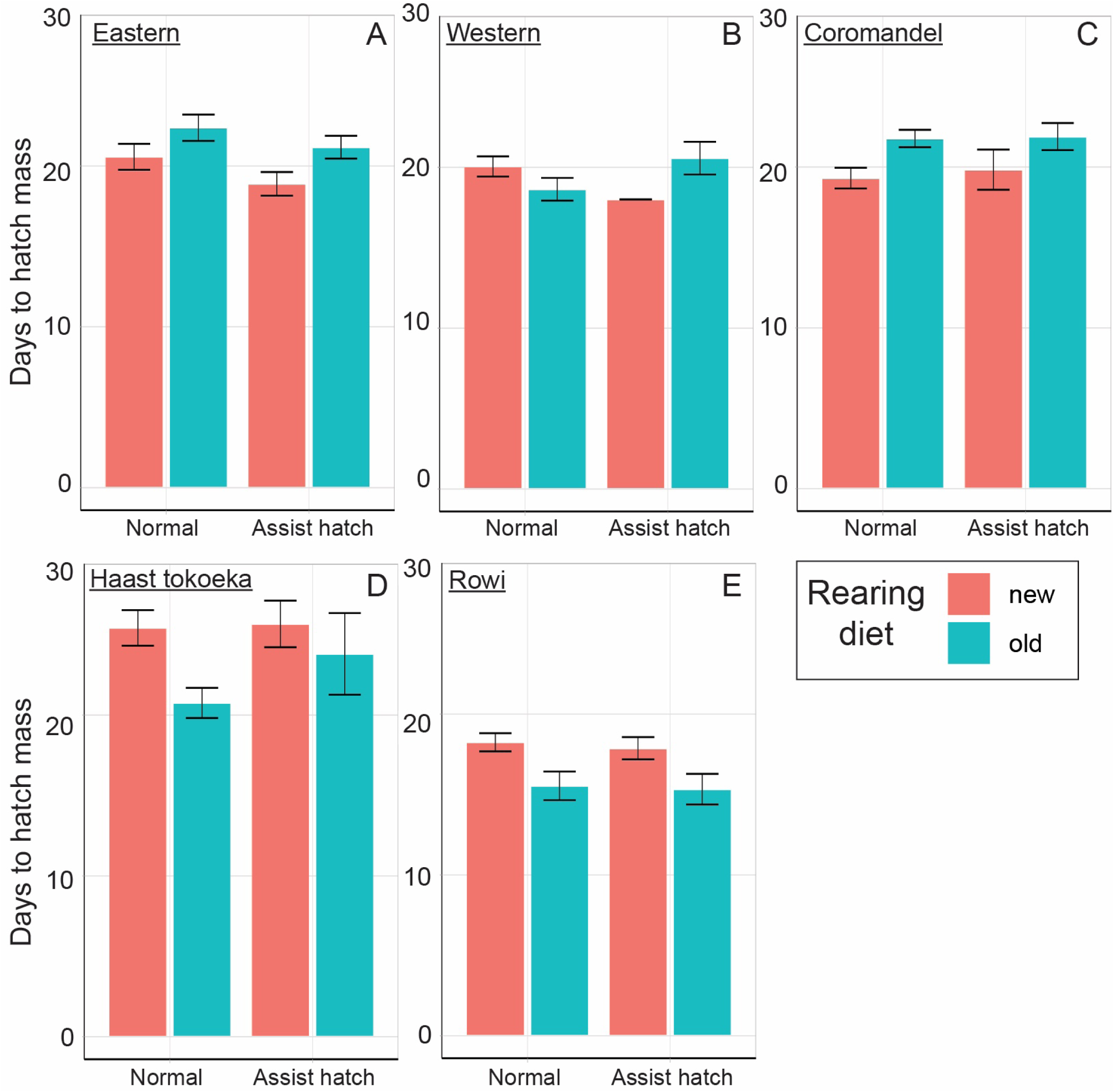
ANOVA showed no differences due to being an assist hatch chick in the number of days it took kiwi chicks to regain their hatch mass. Rearing diet is indicated for reference only - it was not statistically analysed. Assist hatch (AH) and normal hatch (NH) sample numbers: A) Coromandel *N* AH = 15, NH = 39; B) Eastern *N* AH = 33, NH = 48; C) Western *N* AH = 9, NH = 52, D) Haast *N* AH = 8, NH = 36, E) Ōkārito *N* AH = 14, NH = 45.

**Figure 9.**
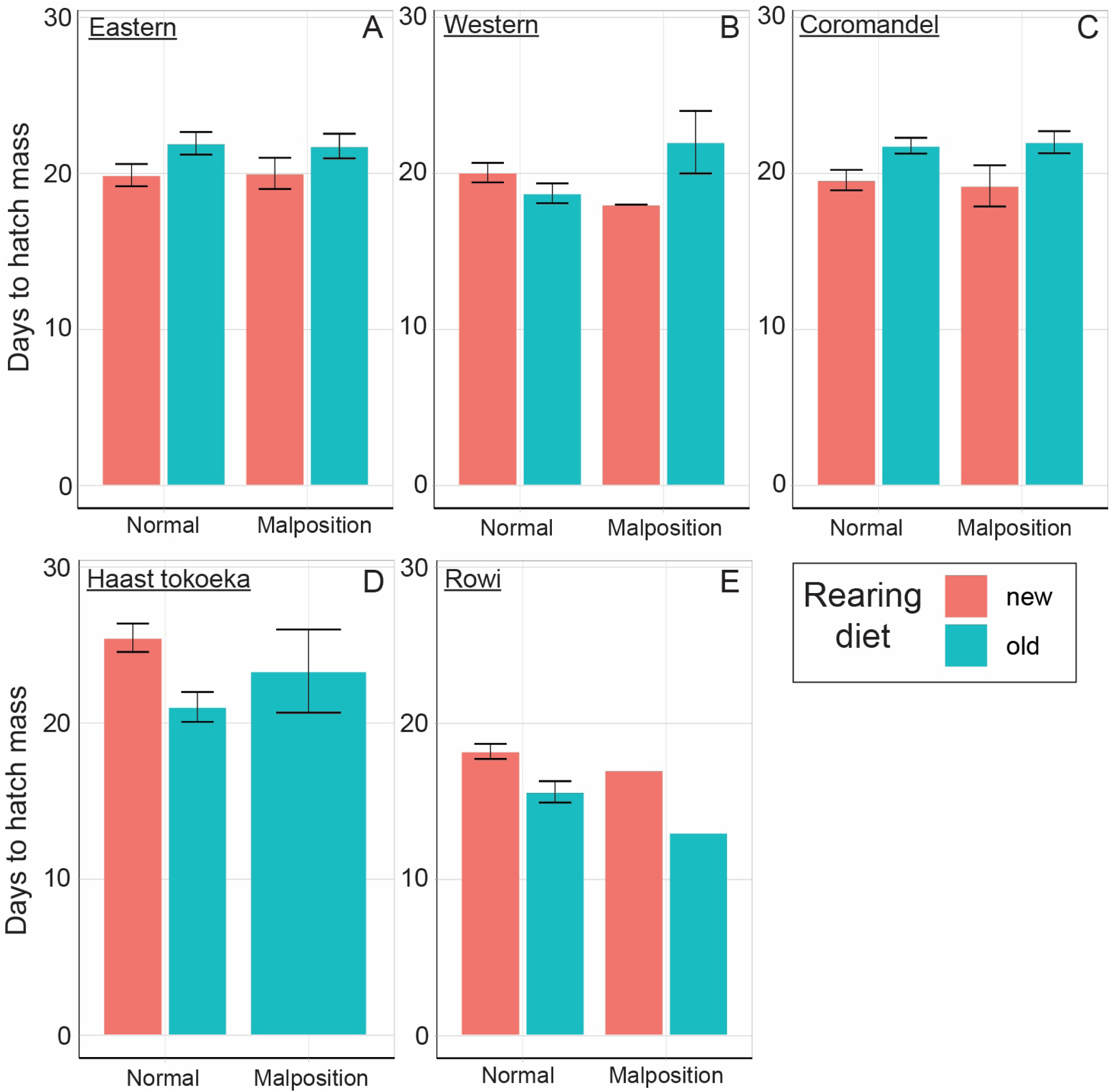
Being a malposition embryo did not impact the number of days it took kiwi chicks to regain their hatch mass. Chick rearing diet is shown here for reference only. Please note, there were only three malpositioned Haast tokoeka embryos recorded (D), and two malpositioned rowi embryos recorded (E). Malposition (Mal) and normal (Norm) embryo sample numbers: Coromandel *N* Mal = 9, Norm = 45, Eastern *N* Mal = 21, Norm = 60, Western *N* Mal = 5, Norm = 56, Haast *N* Mal = 3, Norm = 41, Ōkārito *N* Mal = 2, Norm = 57.

**Table 15.** Three ANOVA on *Apteryx* spp. chicks testing whether developmental time was significantly influenced by genetic group and, A) being an assisted-hatch chick, B) a malpositioned embryo, or C) being assist fed for more than four consecutive days. Chicks did differ in their time to regain hatch mass due to genetic group. Being assist-fed for more than four consecutive days also led to significantly longer developmental times.

| A. Variable | d.f. | Sum sq | Mean sq | F-value | P |
| --- | --- | --- | --- | --- | --- |
| Assist hatch | 1 | 0.0 | 0.42 | 0.03 | 0.862 |
| Genetic group | 4 | 958 | 239.45 | 17.46 | <0.0001 |
| Assist hatch*genetic group | 4 | 397 | 99.30 | 7.906 | 0.255 |
| Residuals | 289 | 3963 | 13.71 |  |  |

| **B. Variable** | **d.f.** | **Sum sq** | **Mean sq** | **F-value** | **P** |
| --- | --- | --- | --- | --- | --- |
| Malposition | 1 | 16 | 16.39 | 0.278 | 0.278 |
| Genetic group | 4 | 940 | 234.95 | 16.906 | <0.0001 |
| Malposition*genetic group | 4 | 22 | 5.54 | 0.398 | 0.810 |
| Residuals | 289 | 4016 | 13.90 |  |  |

| **C. Variable** | **d.f.** | **Sum sq** | **Mean sq** | **F-value** | **P** |
| --- | --- | --- | --- | --- | --- |
| Assist fed > 4 days | 1 | 1253.7 | 1253.7 | 118.841 | <0.0001 |
| Genetic group | 4 | 669.8 | 167.4 | 15.872 | <0.0001 |
| Assist fed > 4 days*genetic group | 4 | 22.4 | 5.6 | 0.531 | 0.713 |
| Residuals | 289 | 3048.8 | 10.5 |  |  |

Assist feeding chicks did however significantly impact their developmental time. Our ANOVA revealed chicks who were assist-fed for more than four days took significantly longer to regain their hatch mass than those who were not (*F*(1,289) = 118.8, *P* < 0.0001, Table 15c, Figure 10). This pattern was not genetic group specific. There was no significant interaction between genetic group and the influence of assist feeding on the time it took kiwi to reach their hatch mass (*assist fed*genetic group*: *F*(1,289) = 0.531 *P* = 0.7131, Table 15c, Figure 10).

**Figure 10.**
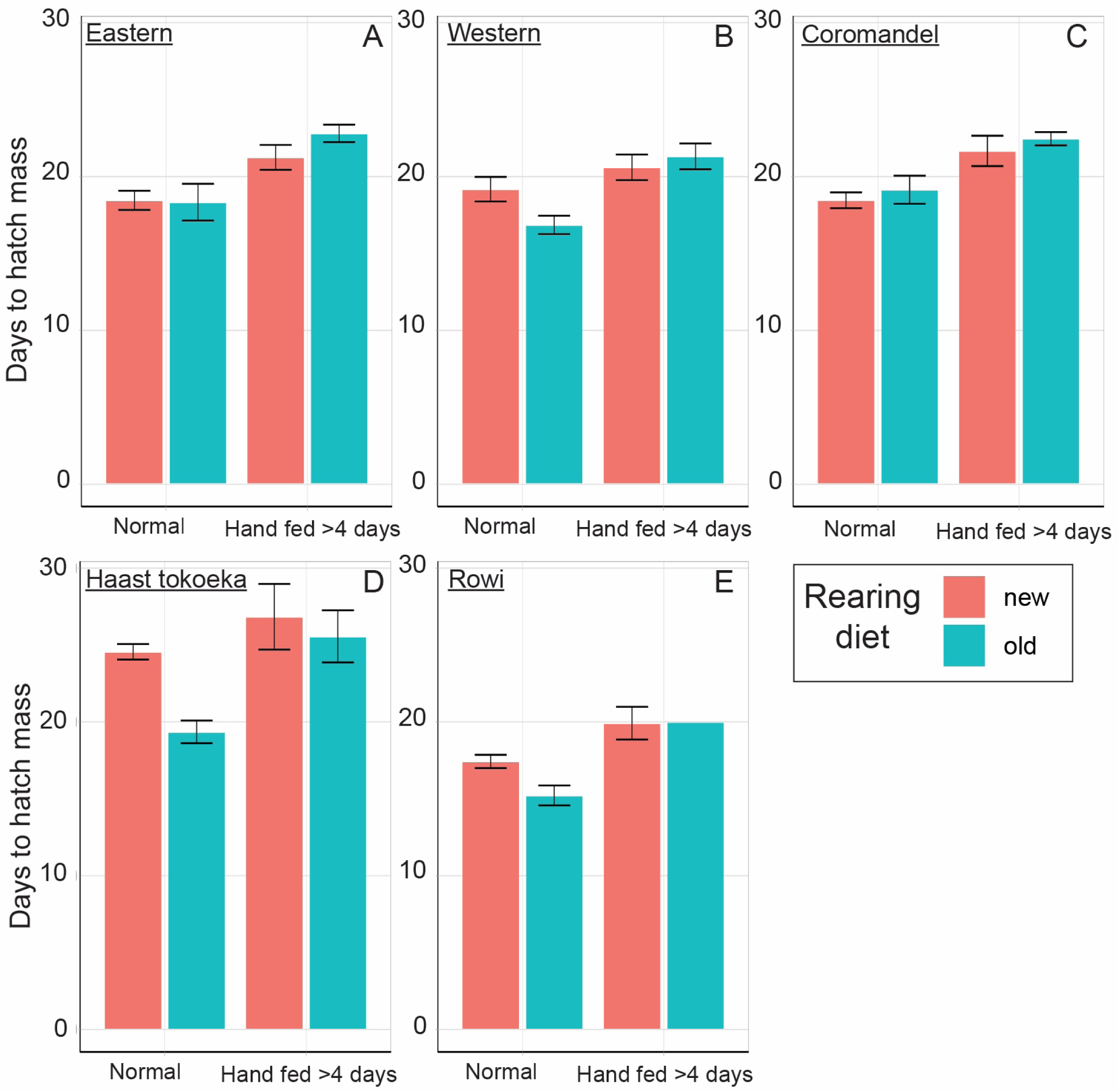
Assist feeding chicks did significantly impact the number of days it took chicks to regain their hatch mass. Chicks who were assist-fed for more than four days took significantly longer to regain their hatch mass than those who were not (compare the bars on the left of the panels to the right-hand bars). Chicks assist fed for more than four days on the old kiwi maintenance (teal bars) diet took longer to regain their hatch mass than chicks assist fed on the new diet (orange bars). Assist fed (AF) and not assist fed (NAF) and new diet (ND) and old diet (OD) sample numbers: Coromandel *N* AF = 34 (ND = 6, OD = 28), NAF = 20 (ND = 13, OD = 7), Eastern *N* AF = 58 (ND = 12, OD =46), NAF = 23 (ND = 11, OD =12), Western *N* AF = 29 (ND = 10, OD = 19), NAF = 32 (ND = 11, OD = 21), Haast *N* AF= 15 (ND = 6, OD = 9), NAF = 29 (ND = 9, OD = 20), Ōkārito *N* AF = 13 (ND = 12, OD = 1), NAF = 46 (ND = 28, OD = 18).

### Differences due to diet and assist feeding

Our ANOVA found a significant interaction between diet and assist feeding. Chicks assist fed for more than four days on the old diet took longer to regain their hatch mass than chicks assist fed on the new diet (*assist fed*diet*: *F*(1,289) = 8.636 *P* = 0.0036, Table 16, Figure 10).

**Table 16.**
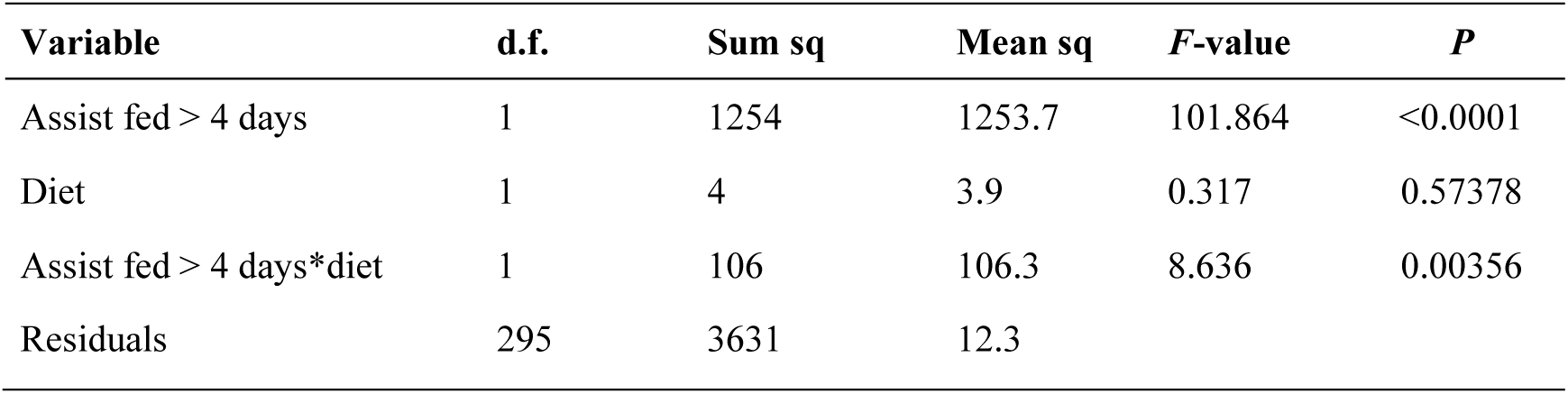
Analysis of variance showing chicks assist fed for more than four consecutive days took longer to regain their hatch mass than chicks that did not require sustained assist-feeding. Chicks assist-fed the old maintenance diet also had a longer developmental time than those assist-fed on the new ZAA Massey diet. Bonferroni adjusted alpha was 0.025.

| **Variable** | **d.f.** | **Sum sq** | **Mean sq** | **F-value** | **P** |
| --- | --- | --- | --- | --- | --- |
| Assist fed > 4 days | 1 | 1254 | 1253.7 | 101.864 | <0.0001 |
| Diet | 1 | 4 | 3.9 | 0.317 | 0.57378 |
| Assist fed > 4 days*diet | 1 | 106 | 106.3 | 8.636 | 0.00356 |
| Residuals | 295 | 3631 | 12.3 |  |  |

### Brown Kiwi (Apteryx mantelli) life history data Hatch mass differences due to sex, clutch number, genetic group, and egg number

Kruskal-Wallis analyses showed no significant relationship between hatch mass and sex in brown kiwi (χ^2^ = 2.2584, d.f., = 1, *P* = 0.1329, Figure 11a) or between hatch mass and clutch number (χ^2^ = 3.9201, d.f., = 1, *P* = 0.047, Figure 11b and c).

**Figure 11.**
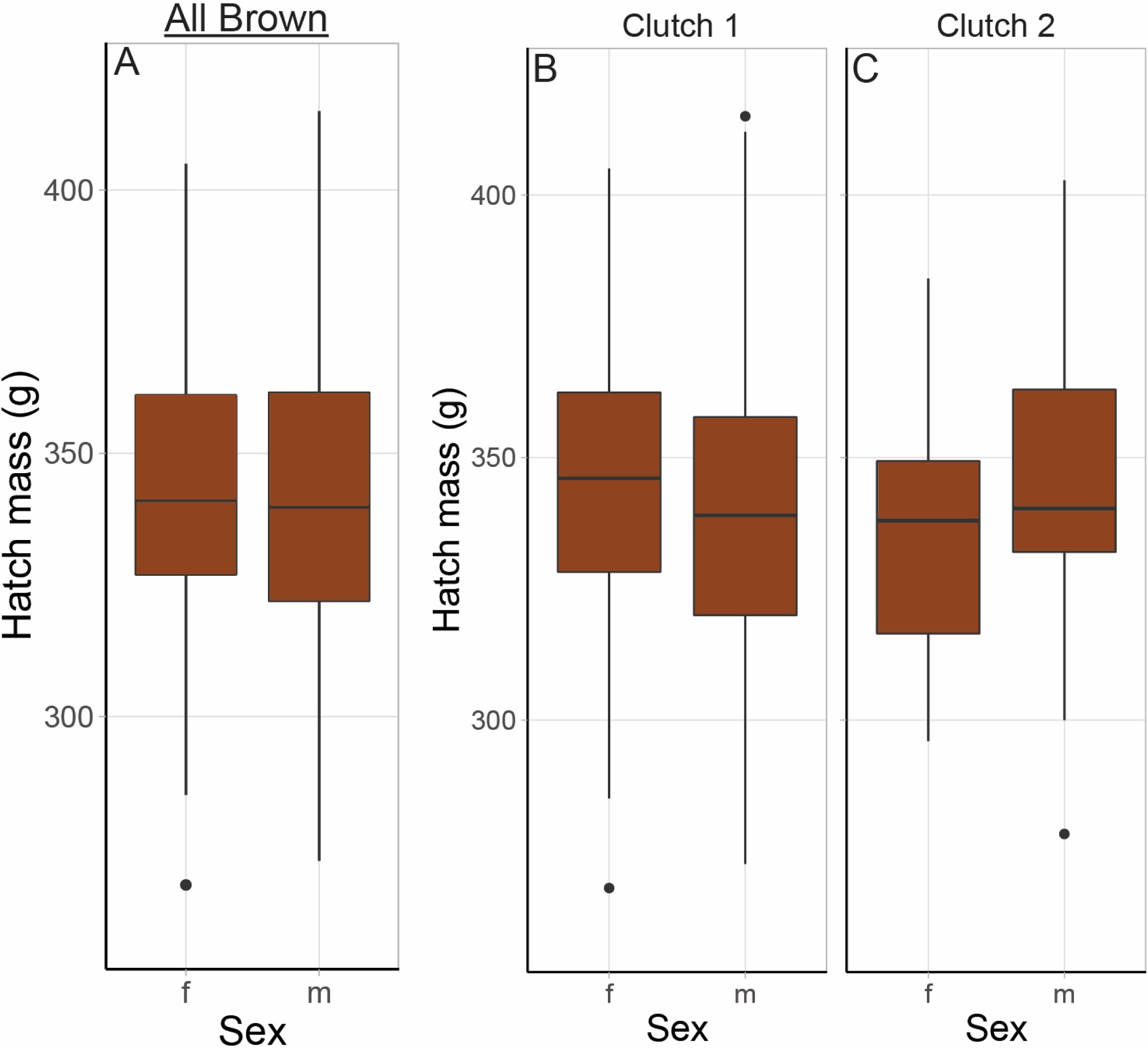
A) Box plot showing that there is no difference between females and males in mass at hatching in Brown kiwi (*A. mantelli*). Kruskal Wallis testing confirmed this statistically. Panels B) and C) depict the hatch mass of first and second clutch brown kiwi chicks of each sex. Kruskal Wallis analyses also found there were no differences in hatch mass between first and second clutch eggs. Boxes and whiskers, from bottom to top, show minimum value, first quartile, the median, third quartile and the maximum value. Sample numbers for female and male first (FC) and second clutch SC) eggs as follows: female *N* = 103 (FC = 74, SC = 29), male *N* = 89 (FC = 59, SC = 30).

Between the genetic provenance groups, hatch mass only approached significant difference (χ^2^ = 8.0859, d.f., = 2, *P* = 0.0175, Figure 12). Planned, Bonferroni corrected (α = 0.0071) post-hoc tests showed Coromandel chicks were significantly lighter at hatch than Western chicks (χ^2^ = 9.458, d.f., = 1, *P* = 0.0021, Figure 12), though as the primary analysis was not significant, this result is reported for completeness only. While Coromandel chicks also appeared to be lighter at hatch than Eastern chicks, this was not statistically supported (χ^2^ = 2.5951, d.f., = 1, *P* = 0.1072). There was also no difference in hatch mass between Eastern or Western chicks (χ^2^ = 1.1034, d.f., = 1, *P* = 0.2935). Note, while sex differences they are shown for completeness (Figure 12) we did not statistically compare sex differences in hatch mass among the genetic groups.

**Figure 12.**
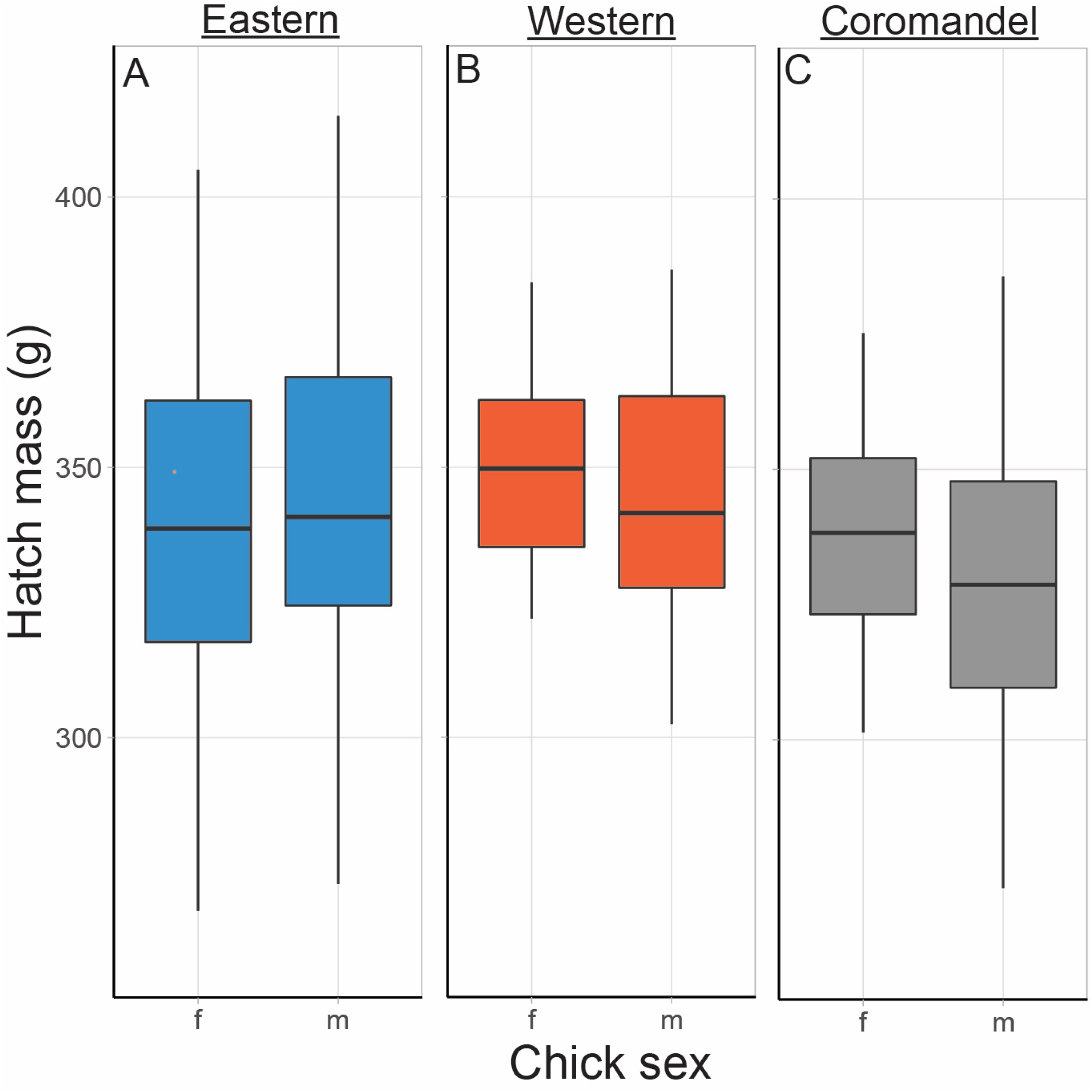
Box plots showing variation in hatch mass for *A. mantelli* from different genetic groups. Box and whiskers show the minimum, first quartile, median, third quartile and the maximum value. Note we did not analyse for interactions with sex, but data are shown for completeness. Sample *N* for each: Coromandel *N* = 52, Eastern *N* = 80, Western *N* = 60).

Hatch mass did not significantly differ between the first or second egg from within a clutch either (Kruskal-Wallis χ^2^ = 0.000243, d.f., = 1, *P* = 0.9876, Figure 13). Due to a lack of statistical power, we did not test for interactions between hatch mass and egg number, or clutch number and provenance group, however, these data are shown (Figure 14). There may be a biologically significant pattern in Coromandel birds (Figure 14c) whereby chicks from the second egg from the second clutch are lighter at hatch (Figure 14).

**Figure 13.**
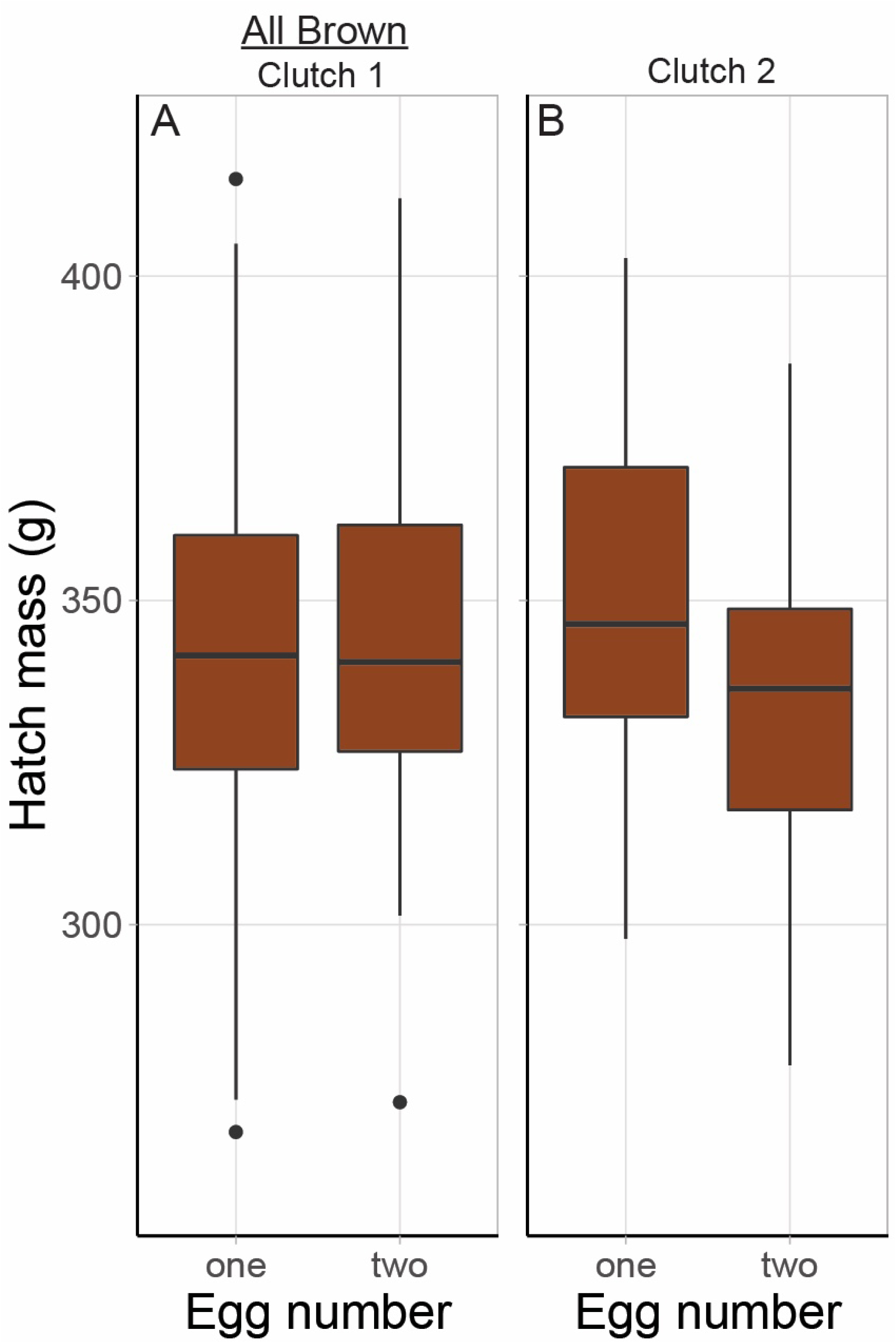
Box and whisker plots demonstrating that hatch mass does not differ between chicks from the first *vs*. second egg from within a clutch. Kruskal Wallis analyses verifies this. Plots show the minimum, first quartile, median, third quartile and the maximum value. *N* for each: first clutch *N* = 133 (1st egg = 73, 2nd egg = 60), second clutch *N* = 59 (1st egg = 24, 2nd egg = 35).

**Figure 14.**
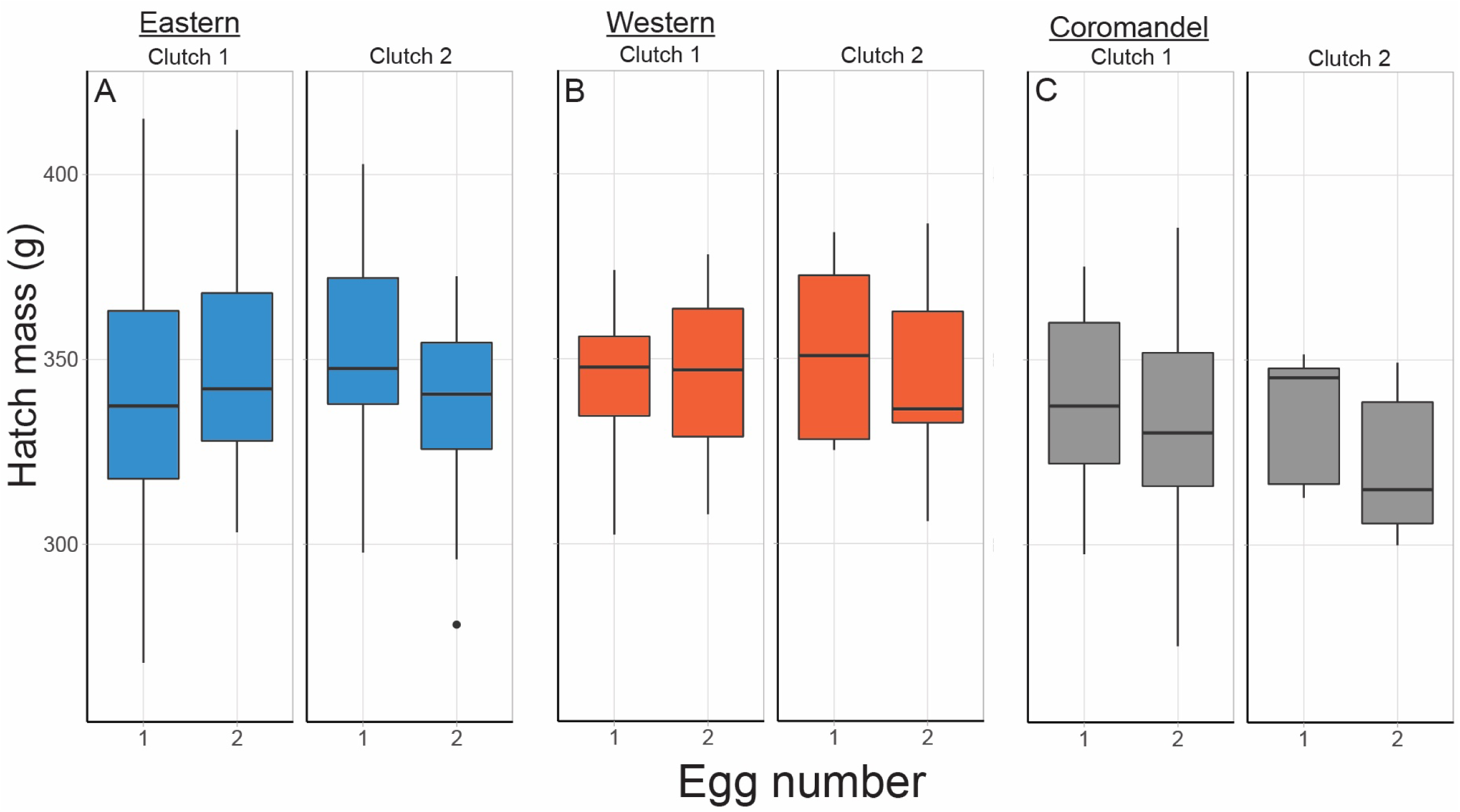
There are no apparent patterns of difference in hatch mass between chicks from egg 1 and egg 2 in Brown kiwi. While there may be an interesting trend for egg 2, second clutch Coromandel chicks, panel (C), to be lighter at hatch. Due to a lack of power, we did not attempt to statistically validate the pattern. Plots indicate the minimum value, first quartile, the median, third quartile and the maximum value. First and second clutch (C1 and C2) and first and second egg (E1 and E2) sample *N* were: Coromandel C1E1 *N* = 17, C1E2 *N* = 18, C2E1 *N* = 5, C2E2 *N =* 9, Eastern C1E1 *N* = 28, C1E2 *N* = 21, C2E1 *N* = 13, C2E2 *N =* 17, Western C1E1 *N* = 28, C1E2 *N* = 21, C2E1 *N* = 6, C2E2 *N =* 9.

### Egg size differences due to sex, clutch number, genetic group, and egg number

Unsurprisingly, Kruskal-Wallis tests found there was no significant relationship between the sex of brown kiwi chicks and the size of their eggs (χ^2^ = 3.1725, d.f., = 1, *P* = 0.07489, Figure 15), or between their egg size and clutch number (χ^2^ = 2.6586, d.f., = 1, *P* = 0.103, Figure 16.).

**Figure 15.**
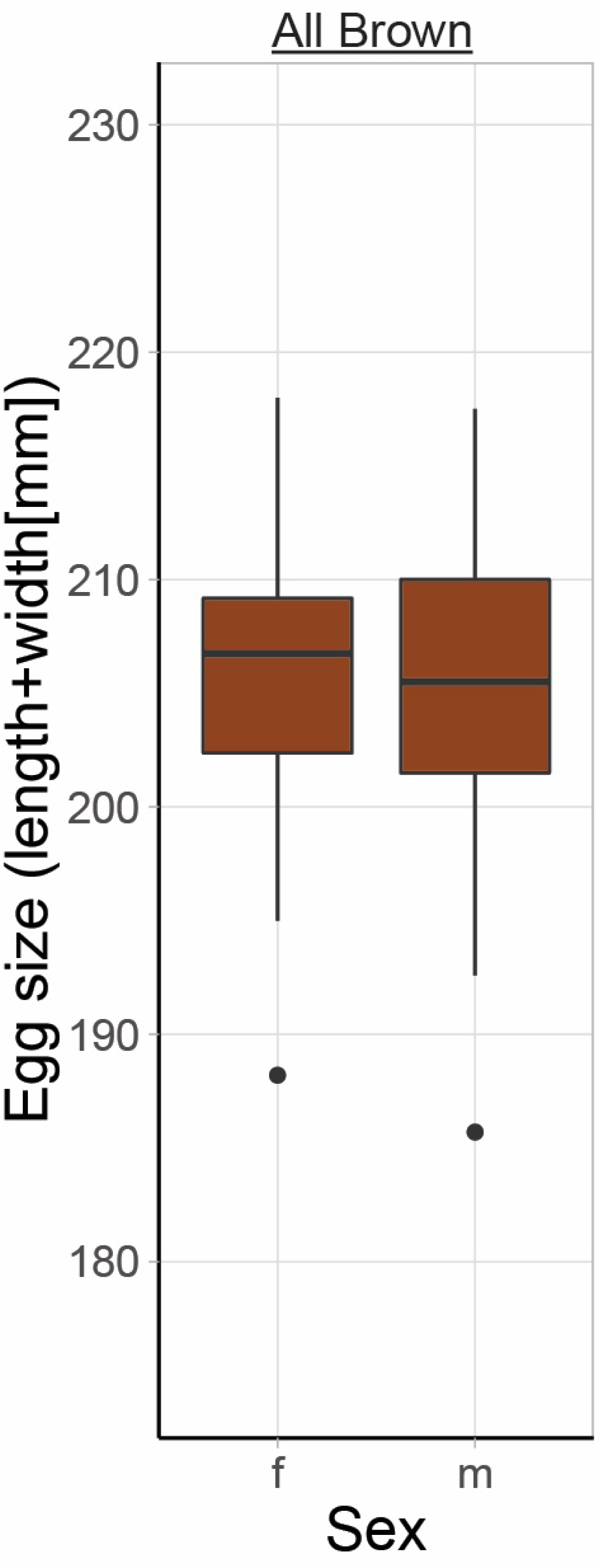
Kruskal Wallis analysis found no significant differences between the sexes of brown kiwi chicks in the size of their eggs. Box and whiskers show the minimum, first quartile, median, third quartile, and the maximum value. Female *N* = 103, male *N* = 89.

**Figure 16.**
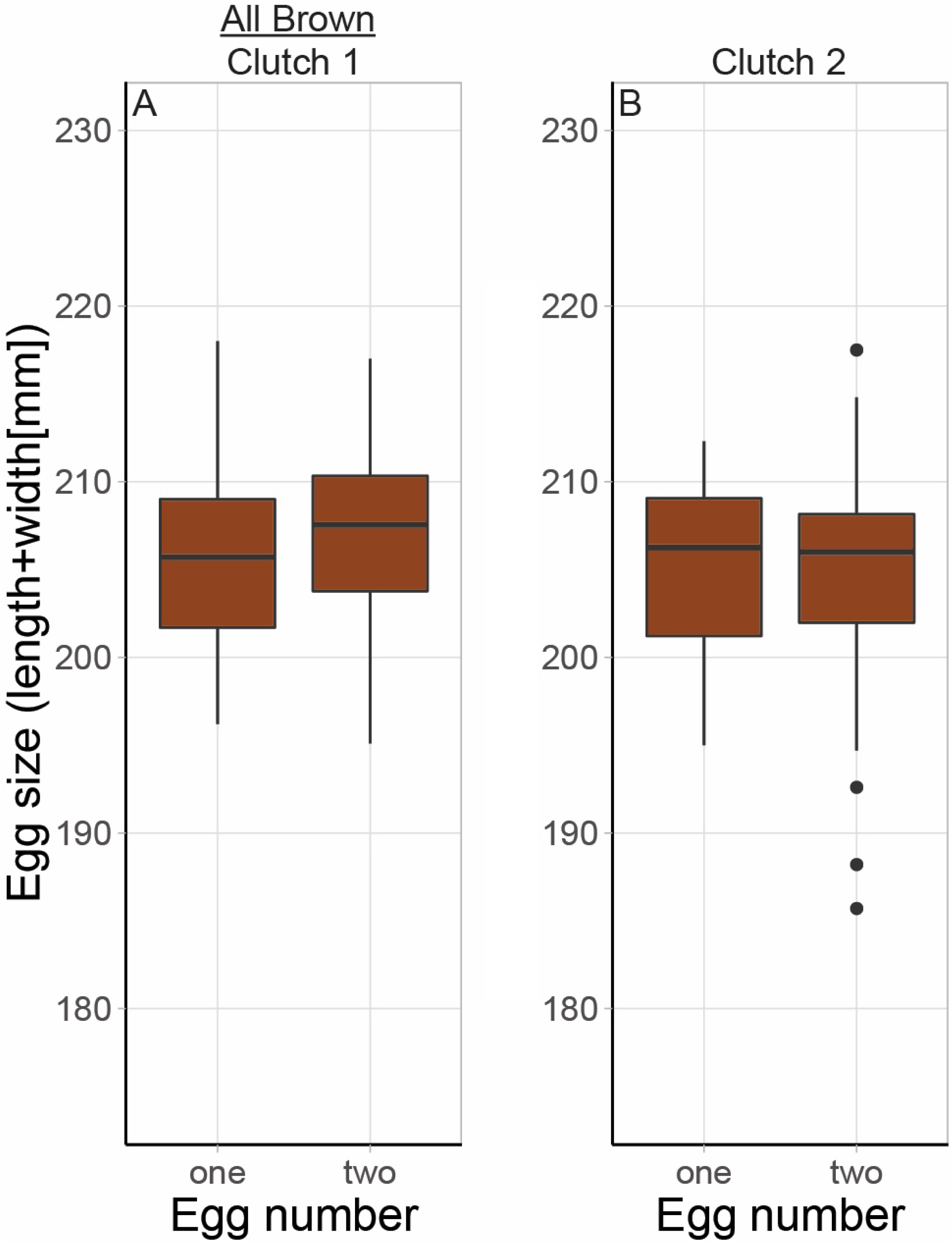
Box and whisker plots displaying that there was no relationship egg size and clutch number in *A. mantelli* eggs. Kruskal Wallis analyses statistically confirmed this pattern. Boxes show the first quartile, median, and third quartile. Whiskers show the minimum and maximum values. *N* for each: first clutch *N* = 133 (1st egg = 73, 2nd egg = 60), second clutch *N* = 59 (1st egg = 24, 2nd egg = 35).

Egg size did differ significantly among the kiwi chick genetic groups (χ^2^ = 11.615, d.f., = 2, *P* = 0.003, Figure 17) with post-hoc tests (α = 0.0071) showing Coromandel eggs being significantly smaller than Eastern (χ^2^ = 7.5205, d.f., = 1, *P* = 0.0061) and Western eggs (χ^2^ = 10.656, d.f., = 1, *P* = 0.0011). There were no significant differences between Eastern and Western eggs (χ^2^ = 0.22938, d.f., = 1, *P* = 0.632). Kruskal-Wallis analysis also showed that egg number within a clutch was not associated with differences in egg size (χ^2^ = 1.0298, d.f., = 1, *P* = 0.3102, Figure 18). As for hatchling mass, we plotted egg size partitioned by clutch number, egg number, and genetic group (Figure 18), and while we did not statistically analyse these sub-groupings, there are no obvious interactions.

**Figure 17.**
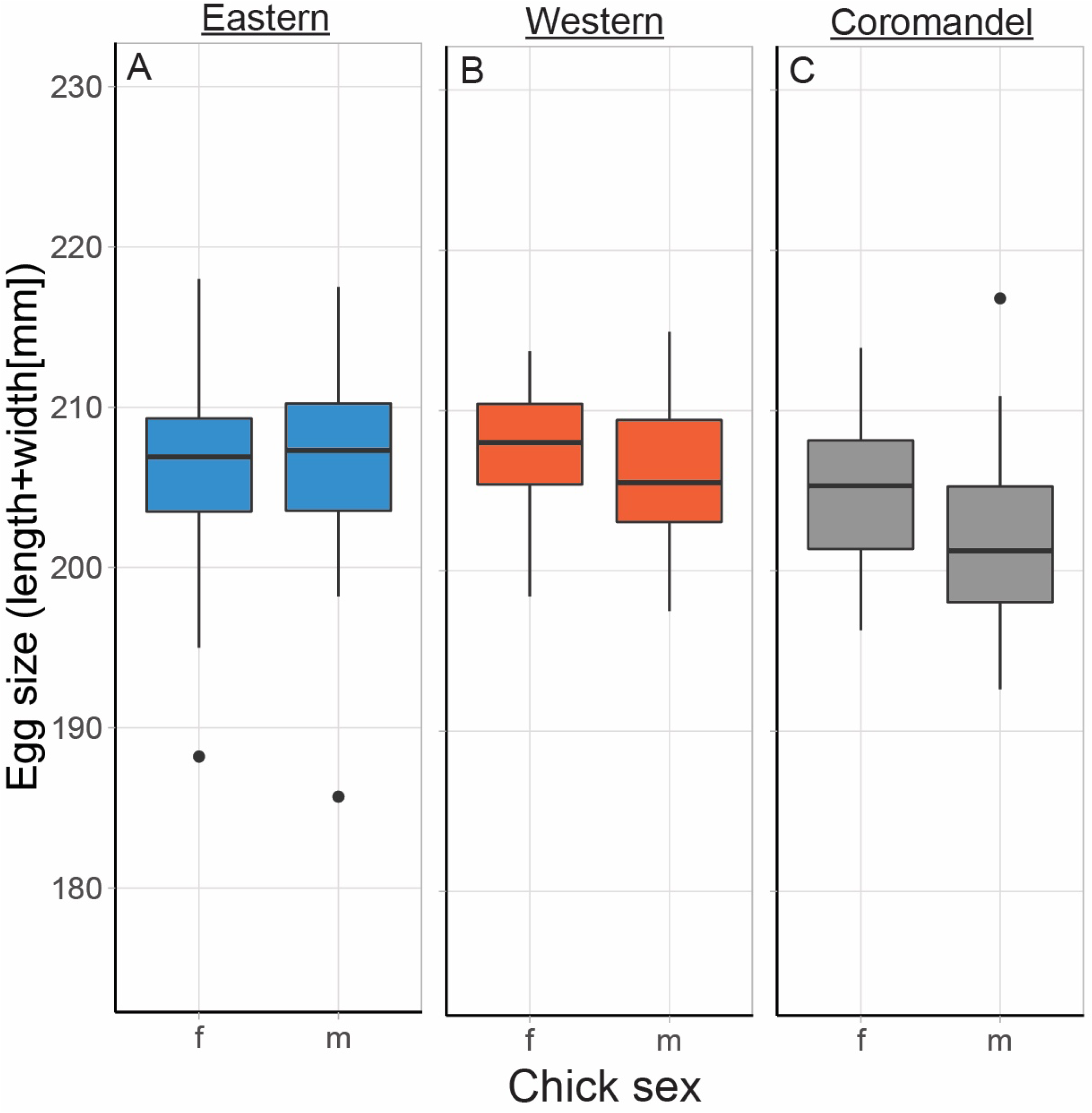
Egg size differed significantly among the Brown kiwi (*A. mantelli*) genetic groups analysed. Kruskal Wallis analyses showed Coromandel (boxes labelled C) eggs were significantly smaller than both Eastern (A) and Western eggs (B). Whiskers show the minimum and maximum values, and the boxes represent the first quartile, median, and third quartile. Note we did not analyse for interactions with sex, but data are shown. Sample *N* for each: Coromandel *N* = 51, Eastern *N* = 79, Western *N* = 60).

**Figure 18.**
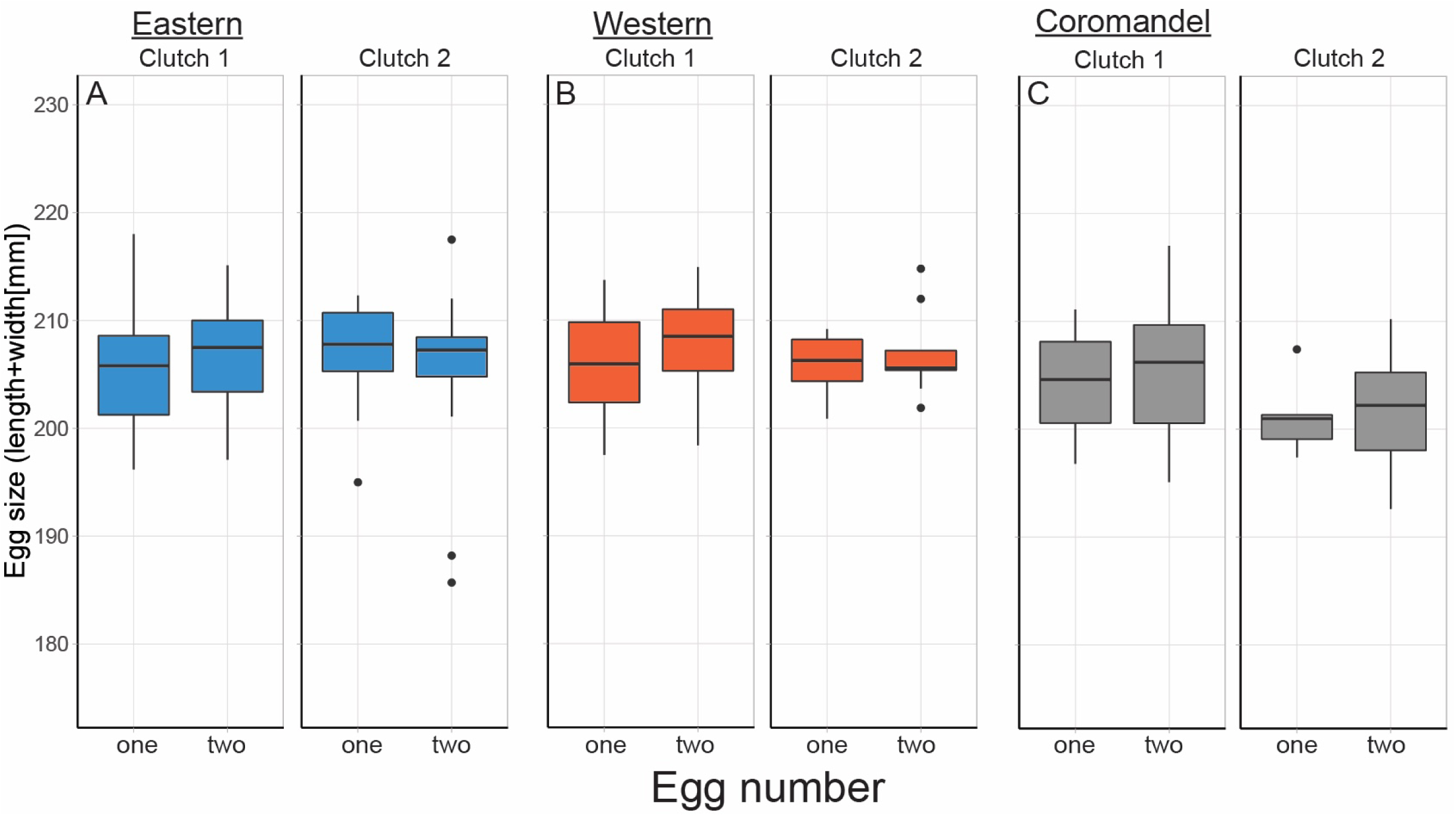
Box plots of egg size data for three genetic groups of Brown kiwi (*A. mantelli*). Kruskal Wallis analyses revealed egg number within a clutch was not associated with variation in egg size. While we did not formally analyse the data shown, there does not appear to be an interaction between egg number, egg size and genetic group. The boxes represent the first quartile, median, and third quartile. The whiskers show the minimum and maximum. First and second clutch (C1 and C2) and first and second egg (E1 and E2) sample *N* where: A) Eastern C1E1 *N* = 28, C1E2 *N* = 21, C2E1 *N* = 13, C2E2 *N =* 17; B) Western C1E1 *N* = 28, C1E2 *N* = 21, C2E1 *N* = 6, C2E2 *N =* 8; & C) Coromandel C1E1 *N* = 17, C1E2 *N* = 18, C2E1 *N* = 5, C2E2 *N =* 9.

### Differences in days to hatch mass (developmental time) due to clutch number and egg number

Kruskal-Wallis testing showed was no relationships between egg clutch number and the number of days it took chick to re-gain its hatch mass in Brown kiwi (χ^2^ = 0.206, d.f., = 1, *P* = 0.6498, Figure 19). Being the second egg laid in a clutch was not associated with developing more slowly (χ^2^ = 4.385, d.f., = 1, *P* = 0.0362 [Bonferroni adjusted α = 0.025], Figure 19).

**Figure 19.**
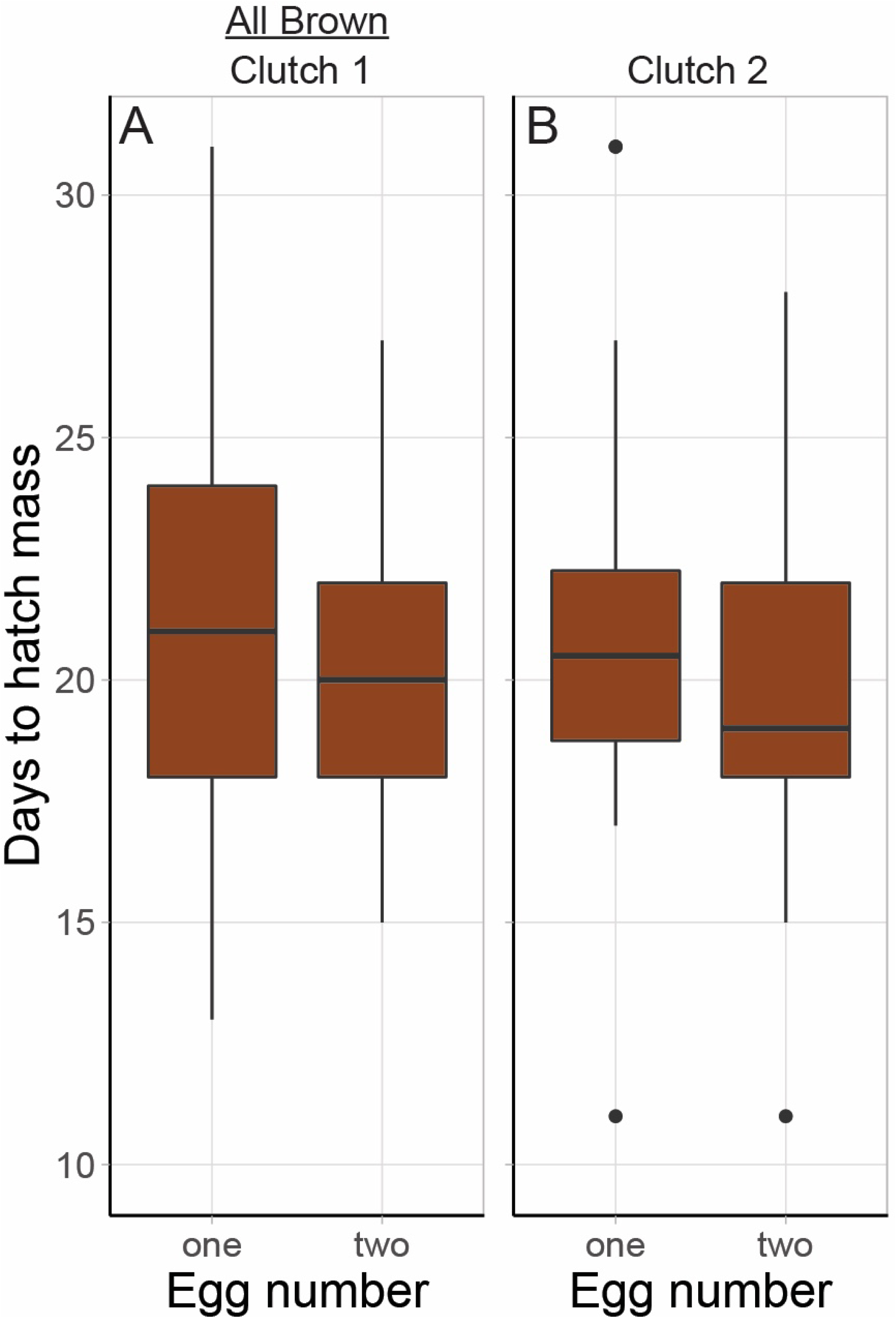
We identified no relationships between egg clutch number and the number of days it took a chick to re-gain its hatch mass in Brown kiwi. However, being the second egg laid in a clutch (i.e., egg number 2 from either clutch) was associated with growing more slowly, however this trend was not statistically supported. Box and whisker plots indicate the minimum value, first quartile, the median, third quartile and the maximum value. Sample numbers were: C1E1 *N* = 43, C1E2 *N* = 34, C2E1 *N* = 24, C2E2 *N =* 33.

### Sex, clutch and egg number

Chi-squared analysis found there was no significant association between a brown kiwi chick’s sex and its clutch number (χ^2^ = 0.045, d.f., = 1, *P* = 0.8312). Out of 130 first clutch eggs, 75 were female (55.6%) and 60 male (44.4%). Of the 57 second clutch eggs, 30 were female (52.6%) and 27 male (47.4%). While there was a slight trend for the first egg laid within a clutch to be female, there was no significant association between egg number (first or second within a clutch) and chick sex (χ^2^ = 0.00074, d.f., = 1, *P* = 0.9783). From the 98 eggs that were the first laid in their clutch 53 were female (54%) and 45 male (46%). Of the 94 eggs that were laid second within a clutch, there were 52 females (55.3%) and 42 males (44.7%).

### Relationships between sires, malpositioned embryos, hatch types, and hand feeding needs

Chi-squared test showed a significant relationship between the number of malpositioned embryos and kiwi genetic provenance groups (*χ*^2^ = 25.8, *P* = 0.0011, Table 17). There was a lot of variation in the incidence of malpositioned embryos among the kiwi genetic groups. The Eastern group had the most malpositioned embryos, with 17 malpositioned chicks of 79 eggs (incidence of 21.5%). Next was Coromandel with 9 malpositioned embryos from 54 eggs (16.67%), then Western (5 malpositioned from 65 eggs = 7.7%), then Haast tokoeka with 3 malpositioned chicks from 47 eggs (6.4%). Ōkārito brown kiwi (rowi) has the fewest, with only 3 of 57 eggs containing a malpositioned embyro (3.5%).

**Table 17.**
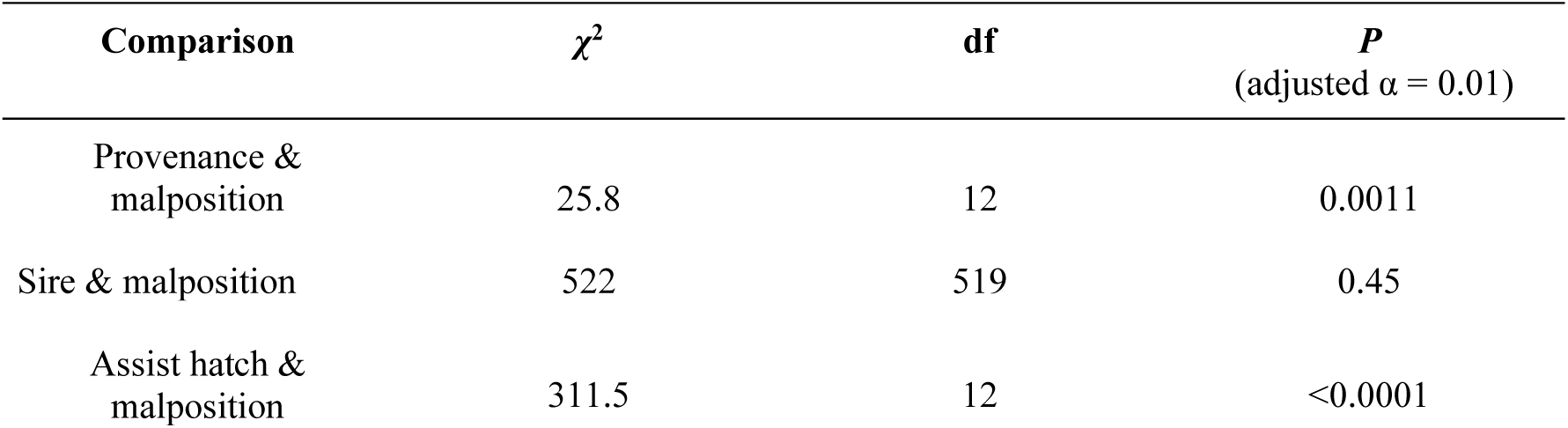
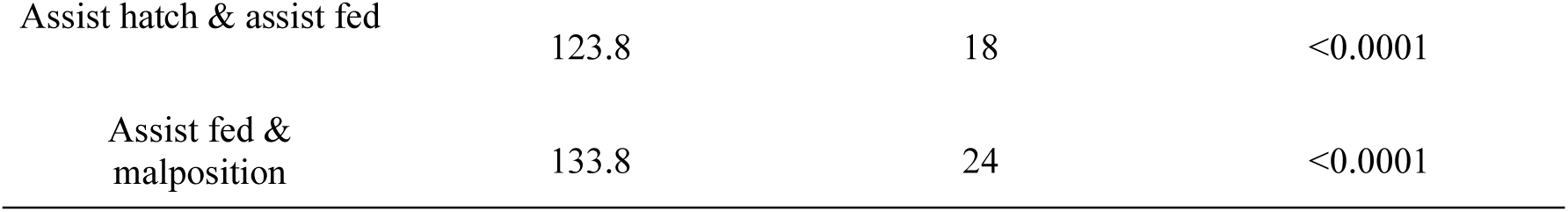
Multiple Chi-square tests of independence exploring data on malpositioned embryos, assist-feeding, and assist hatching, from chicks from five kiwi genetic groups (*Apteryx* spp.). There was no significant relationship between individual sires and their likelihood to produce malpositioned embryos. There was a significant association between genetic provenance group and the number of malpositioned embryos. For example, 21.5% of the Eastern genetic group eggs contained malpositioned embryos, while *A. rowi* (Ōkārito brown) kiwi only had malpositioned embryos in 3.5% of their eggs. Malpositioned embryos were also significantly more likely to require an assist hatch, and significantly more likely to require consecutive days of hand-feeding.

| Comparison | $\chi^2$ | df | $P$<br>(adjusted $\alpha = 0.01$) |
| --- | --- | --- | --- |
| Provenance & malposition | 25.8 | 12 | 0.0011 |
| Sire & malposition | 522 | 519 | 0.45 |
| Assist hatch & malposition | 311.5 | 12 | <0.0001 |
| Assist hatch & assist fed | 123.8 | 18 | <0.0001 |
| Assist fed & malposition | 133.8 | 24 | <0.0001 |

Despite Eastern sires producing more malpositioned embryos, chi-squared testing did not identify a relationship between individual sires, Eastern or otherwise, and the number of malpositioned chicks produced (*χ*^2^ = 522, *P* = 0.45, Table 17).

There was a significant relationship between being a malposition embryo and requiring an assisted hatch (*χ*^2^ = 311.5, *P* < 0.0001, Table 17). There were also significant relationships between being a malpositioned embryo and an assisted hatch and requiring more than four days on continuous hand-feeding (*malposition cf. assist feed*: *χ*^2^ = 133.8, *P* < 0.0001, Table 17; *assist hatch cf. assist feed*: *χ*^2^ = 123.8, *P* < 0.001, Table 17). The contingency table used for these analyses is shown in Table S7.

## DISCUSSION

### Growing up on the new *vs.* old kiwi maintenance diets

Our findings suggest kiwi, especially the South Island taxa, do have some genetically determined differences in their developmental rate and in how they cope metabolically when growing on foods of varying nutritional composition. Coromandel and Western chicks grew the most efficiently over time on the new diet. Eastern chicks were intermediate in their growth efficiency, while Rowi and then Haast tokoeka chicks grew the least efficiently (trait responses are summarised in Table 18). This means that rowi and Haast chicks had to eat more of the new diet each day to achieve body mass increases than the Coromandel, Western, and then Eastern chicks. Haast chicks also developed more slowly on the new diet compared to the other genetic groups, while rowi chicks developed the fastest.

**Table 18.** Summary of the main differences between the studied kiwi genetic groups in their nutritional responses to the new *vs.* old kiwi maintenance diets and their life history characters. When reading across the row, comparisons are given for within each diet type. For example, compare results for rowi “old” to Haast “old”, but not rowi “new” to rowi “old”.

| Response variable | Haast tokoeaka |  | Rowi |  | Coromandel |  | Western |  | Eastern |  |
| --- | --- | --- | --- | --- | --- | --- | --- | --- | --- | --- |
|  | new | old | new | old | new | old | new | old | new | old |
| Growth efficiency | lowest | low | low | lowest | highest | highest | highest | high | high | high-medium |
| Development time | slowest | slow | fastest | fastest | fast | slow | medium | medium | medium | slow |
| Food intake (inferred) | highest | high | high | highest | lowest | low | lowest | low | low | medium |
| Mass loss before feeding | highest |  | high |  | high |  | lowest |  |  | low |
| Hatch mass |  |  |  |  | lightest |  | intermediate |  |  | heavier |
| Egg size |  |  |  |  | smallest |  | larger |  |  | larger |
| Malpo embryos (rank 1 - 5) | 4 |  | 5 (least) |  | 2 |  | 3 |  | 1 (most) |  |

On the old diet, Coromandel chicks grew the most efficiently, with Western and Eastern chicks intermediate. Again, as on new diet, the South Island chicks grew the least efficiently. On the old diet, it was the rowi chicks that grew the least efficiently, then the Haast. However, rowi again developed faster compared to all the other genetic groups, especially when reared on the old diet. This means that again, rowi had to eat more of the old diet over less time, to maintain their fast development.

Of note are the differences in mass loss over the first eight days, as the chicks absorbed their yolk (Dzialowski & Sotherland 2004). Haast chicks lost the most mass, followed by Coromandel, rowi, Eastern, and then Western chicks. While these differences are interesting and warrant further investigation, they do not suggest losing more mass prior to feeding (i.e., metabolising a bigger yolk), sets a chick up to have a faster development rate or more efficient growth. The genetic group which grew the most efficiently, irrespective of diet - the Coromandel kiwi - lost nearly as much mass prior to feeding as the Haast and rowi, both of which grew inefficiency on both diets and showed opposite trends in their development time.

Overall, the new diet supported more similar growth efficiency outcomes between genetic groups, and better supported the potential stresses of ONE in the brooder room (for example, being hand-fed) than the old diet.

### Macronutrient differences between the diets and genetic group responses

The old and new maintenance diets differ in their energy density, macronutrient ratios, and in the levels and ratios of key micronutrients. Feeding on one or the other diet would present substantially different physiological challenges and outcomes. The new diet is more energy dense, offering almost twice as many calories (kcal) per unit intake than the old diet. This is because the new diet contains over three times as much fat. Over 50% of the new diet’s total energy comes from fat, compared to 27.4% of energy in the old. Protein provides 32% of the energy in the new diet, whereas in the old diet, which contained far more beef meat, 57.6% of total energy was from protein – a less energy dense macronutrient. The carbohydrate derived energy content of the new and old diets was comparable (17.5% and 15% respectively), suggesting any macronutrient related differences in performance between chicks reared on either food were driven by protein and/or fat.

The estimated wild brown kiwi (*A. mantelli* [from the Northland genetic group]) adult diet (Kleinpaste 1990; Potter et al. 2010) is most similar in fat composition to the new kiwi diet, offering 40.2% of total energy from fat. In terms of protein content, the wild diet, with protein providing 57.6% total energy, is more similar to the old maintenance diet where protein supplied 49.6% of the energy. How similar the estimated wild diet is to wild hatchling kiwi diets is unknown and needs investigation. Most kiwi diet studies have focused on adult wild diet (Reid et al. 1982; Colbourne & Powlesland 1988; Klienpaste 1990).

The South Island species chicks showed the strongest responses to the two diets, and it is likely the rowi and Haast chick diets require review. The rowi grew less efficiently than the North Island genetic groups on both the high-protein, low energy old diet, and the high-fat, high calorie new diet. Though despite the *inefficient* growth, rowi chicks developed the fastest. This combination of inefficient growth, yet fast development indicates rowi were required to eat more, of both diets, in less time, to support what may be a naturally fast developmental rate. If rowi are constrained to develop quickly, regardless of diet, the current ONE situation that requires them to consume large volumes of fixed nutrient ratio food in a short time would present physiological consequences. A similar issue is faced by Haast tokoeka chicks. Like the rowi, Haast chicks grew inefficiently on both diets. Though the Haast developed more slowly than the other chicks, especially on the new high-fat diet. Haast chicks were required to eat very high volumes of both foods over a protracted developmental period. Some Haast chicks for example, took nearly two months to reach release from the brooder room on the new diet. The health consequences for over-consumption of one or another macronutrient are well established. Over-consumption of protein can promote mitochondrial malfunction (Sanz et al. 2004; Ayala et al. 2007); illness due to toxic nitrogenous waste accumulation (Raubenheimer and Simpson 2009); kidney changes and damage (Walker et al. 1989; Sabat et al. 2004); major changes in innate immunity (Ponton et al. 2011); and a decrease in immune function, that continues the longer animals are forced to feed on high-percentage protein foods (Dzhumagaziev 1980). Concerning for ONE birds who were raised on the old diet (which was used for over 10 years) are recent findings that show negative trans-generational effects of high protein feeding by parents, for even short periods, increases inflammation and decreases B cells responsiveness in their offspring – even when those offspring are returned to an optimal diet (Gruber et al. 2018). The health and fitness costs associated with over-feeding on fat during development include chronic inflammation (Esposito & Giugliano 2006; Lee et al., 2010; Wang et al. 2012), which can impede immune function (Kiran et al. 2022), lead to auto-immune disease (Versini et al. 2014), cardiovascular disease and excess adiposity (Cottrell and Ozanne 2008), and cause negative trans-generational effects (Langley-Evans 2015).

While little work has been published on the specific macronutrient needs of kiwi, researchers have estimated the protein requirements for kiwi chicks, which are known to display exceptionally slow growth as older juveniles (McLennan et al. 2004; Heck & Woodward 2021). Using data from McLennan et al. (2004) from growing wild kiwi chicks, Sales (2006a) fitted Gompertz body growth equations and estimated their percentage protein requirements. Sales (2006a) estimated that a 10 day old, 313 g female kiwi chick would require 4.2 g of protein per day, and a 10 day old, 319 g male kiwi chick would require 4.0 g of protein per day. Most 10 day old ONE chicks (of all species) eat at least 15 grams of artificial diet per day (see SM1). When feeding on the old diet, a 10 day old chick eating 15 g of diet (wet weight) would have been ingesting 2.4 g of protein per day. If the requirements for males and females is averaged to 4.1 g of protein per day, this falls 1.7 g of protein short of the Sales (2006a) estimate for healthy growth. The new diet is only marginally better. A 10 day old chick on the new diet would intake 2.5 g of protein, 1.6 g short of the recommendation. The natural diet described in Potter et al. (2010) is only marginally better, offering 2.98 g of protein, 1.2 g short of the Sales (2006a) recommendation. It is possible that all these diets, which are all adult bird maintenance diets, are too low in protein to sustain the growth rates of very young chicks described by McLennan et al. (2004). For older chicks Sales (2006a) estimated a 454 g, 50 day old female would require 5.1 g of protein per day, and a 460 g male would require 4.6 g of dietary protein. By day 50, the majority of ONE chicks are eating at least 110 g of the artificial diet each day. A daily intake of 110 g on the old diet would provide 17.7 g of protein. While consuming 110 g of the new diet would provide 18.4 g of protein, and 110 g of the adult wild diet (Potter et al. 2010) provides 21.9 g protein. If the Sales (2006a) recommendations for 50 day old kiwi are accurate, then all of these diets would provide ample, if not too much, protein/g for older birds. Prier et al. (2013) have commented that captive diets may not accurately support the growth conditions for kiwi chicks, and we suggest that if the protein requirements determined by Sales (2006a) using the data from McLennan et al. (2004) are accurate, then all three diets are likely too low in protein for early development and become too high in protein later. We have anecdotal evidence from keeping ONE chicks, that as they approach release weight (∼ 800 - 900 kg) some chicks display bone and joint problems, indicative of too rapid growth, something an excessively high percentage protein diet could be contributing to (see Prier et al. [2013] for a discussion). Work in insects (de Carvalho & Mirth 2017; Rho & Lee 2022) and mice (Sørensen et al. 2008) has demonstrated that nutritional preferences, that are tightly linked to optimal performance, shift during development. Mirroring the estimates of Sales (2006a), work on these animals shows that the need for high percentage protein foods decreases over developmental time.

### Improving the captive chick diets macronutrition

The differences observed among the ONE kiwi chicks are likely due to each genetic group having subtly different macronutrient intake requirements and abilities to minimise the physiological costs of macronutrient over-nutrition. Animals, of all life stages, use innate nutritional wisdom” (*sensu* Simpson & Raubenheimer 2012) to regulate their diet’s macronutrient composition. They do this by selecting foods that achieve a “target” nutrient balance over time. For animals on fixed ratio captive diets, this is not possible. Straightforward work could determine the differing nutritional needs of growing chicks from each genetic group and help improve their captive diets. First, we should conduct nutritional choice experiments on chicks of different ages. There are well established protocols for conducting choice experiments with animals to measure their dynamic macronutrient intake targets [see work on ibis (Coogan et al. 2017), dogs (Hewson-Hughes et al. 2013), mice (Sørensen et al. 2008), moose (Felton et al. 2016), cockroaches (Raubenheimer & Jones 2006), flies (Lee et al. 2008), beetles (Rho & Lee 2022), and locusts (Raubenheimer & Simpson 1993)]. These simple experiments should begin on captive kiwi (of all ages and life stages) without delay. These choice experiment methods are straight-forward, intuitive, and given the correct suite of foundation diet ingredients, low-cost. They could be easily implemented by captive rearing facilities, arguably for any species that is non-habituated to particular foods. These investigations should be complimented by research on chick’s wild diet. Traditional feacal analyses (e.g., Colbourne & Powlesland 1988; Klienpaste 1990) would be a good starting point for kiwi chicks. Environmental DNA from faeces could also be analysed using high-throughput illumina-flow techniques that would allow for total dietary species number and abundance estimates from faecels (De Barba et al. 2014; Deiner et al. 2017). Isotope analysis of body tissues and environmental food sources could also occur (Wang et al. 2023), and the findings from the continuing work of San Juan et al. (2021) on the consequences of captive *vs*. wild diets on kiwi gut microbiota should be incorporated. These wild-diet discoveries should guide the food substrate choices used in the feeding experiments.

Given the scale of ONE, and the behavioural, morphological and evolutionary consequences on inappropriate captive rearing protocols (Alberts 2007; Passos et al. 2017; Brown et al. 2021) the natural kiwi chick diets, and their region-specific nutritional needs should become a research priority. It is known that when species are nutritionally mis-matched to their optimal, dynamic, natural diet, individuals can over- or under-investment in life history traits, especially those associated directly with adult fitness (Lee et al. 2008; Solon-Biet et al. 2015; Gray et al. 2018). It is concerning that the potential impact of the new diet on ONE chicks (and historically the old diet), are not being monitored for, or measured. Currently, ONE chicks are known to have lower post-translocation survival than wild-reared chicks (Jahn et al. 2022). Working out how to minimise the potential physiological and evolutionary impact of ONE is critical.

### Eco-physiological explanations for genetic group differences in nutritional responses

Species differ in how well they can mitigate costs when confined to inappropriate diets. Nutritional ecology conceptualises “nutritional generalists” and “nutritional specialists” (Machovsky-Capuska et al. 2016). Generalists species occur in climatic and nutritionally variable ecological environments, and have evolved to better maintain homeostasis when feeding across a broad range of nutritionally and temporally variable foods than specialists. Nutritional specialists tend to evolve in homogenous food environments and, over time, are less well able to tolerate foods with a nutritional composition far from their evolutionarily familiar “target” (Raubenheimer & Simpson 2018). While conjectural, it is possible rowi can better minimise costs associated with high-protein, and especially, high-fat feeding than Haast chicks who may be more “specialist”. Haast may need to take a slower staged approach to metabolising excessive protein, and especially fat.

Interestingly, Coromandel chicks grew the most efficiently of all genetic groups, on both diets. While all these patterns warrant further investigation, perhaps Coromandel chicks are the most “generalist” of all genetic groups. It is true that Coromandel ecological district experiences drier summers, more frequent heavy rain events with more flooding, more frequent summer droughts than the West Coast ecological districts where rowi and Haast come from (McEwen 1987). The West Coast also has more stable temperatures year round and higher floristic and vegetation uniformity than the Coromandel (McEwen 1987). Differences in soil type, rainfall consistency and therefore soil penetrability could also drive adaptive differences among kiwi (Kleinpaste 1990).

### Micronutrient differences between the diets

The new kiwi maintenance diet provides better overall micro-nutritional support to growing chicks than the old diet (Table 19). The well-established calcium (Ca) to phosphorus (P) ratio recommended for growing hatchling and juvenile birds is 1Ca:1P to 2Ca:1P (Pathak 2021). Compared to the old diet, which offers only 1C:13.5P, the new kiwi diet offers a better level of calcium to phosphorus at 1Ca:3.9Ph, though this is likely to still be too high in phosphorus to facilitate optimal absorption and utilisation of either micro nutrient (see discussion in Table 19), as high-phosphorus intake can actively inhibit calcium absorption, even if dietary calcium intake is otherwise adequate (Takeda et al. 2014; Manopriya et al. 2022). The new diet also offers substantially higher levels of several carotenoids than the old diet (Table 19), including cryptoxanthin, which optimises bone growth (Burri et al. 2016). The levels of essential omega-6 and omega-3 polyunsaturated fatty-acids in the new diet, are also an improvement over those from the old kiwi diet. Linoleic acid (an omega-6) and Alpha-linolenic acid (an omega-3) are important metabolite precursors (Marangoni et al. 2020) and are critical in nerve and brain development (Schuchardt et al. 2010). Lower Linoleic acid (an omega-6 fatty-acid) to Alpha-linolenic acid (an omega-3) ratio nutritional sources are known to promote better neurone development (Schuchardt et al. 2010), and to lower inflammation (Bork et al. 2019). The new kiwi diet contains more than twice as much of both of these fatty-acids than the old diet due to the addition of canola and corn oils to the diet recipe. The new kiwi maintenance diet’s ratio of omega-6:omega-3 of 1.6 to 1 also more closely approaches the general 2:1 to 4:1 recommended ratio for animals than the old diet, at 1.3 omega-6 to 1 omega-3 (though both diets offer the fatty-acids at an acceptable ratio [Alagawany et al. 2019]). In addition to refining the macronutrient requirements for each age-stage and kiwi taxa, a major improvement to the kiwi diet for growing chicks (and also laying kiwi hens) would be to reduce dietary phosphorus and possibly omega-6 fatty acid levels (Ponnampalam et al. 2021), by reducing the reliance on beef meat as the major protein source in the kiwi diet recipe. A practical way to do this, which would also improve behavioural outcomes (Mellor et al. 2009) and preparedness for release to the wild (Sales 2006b), would be the inclusion of more live insect food, or insect derived foods into the kiwi diet. The nutritional value of insects is being increasingly recognised and quantified as human consumption increases, and insects are becoming more available (Hartmann et al. 2018; Higa et al. 2021). Black Solider Fly (*Hermetic illucens*), the larvae of which contain the ideal 2Ca to 1P ratio and are 50% protein and 15% fat (Shah et al. 2022) are readily cultivated (Higa et al. 2021) and would be an ideal candidate with which to begin feeding trials in kiwi.

**Table 19.**
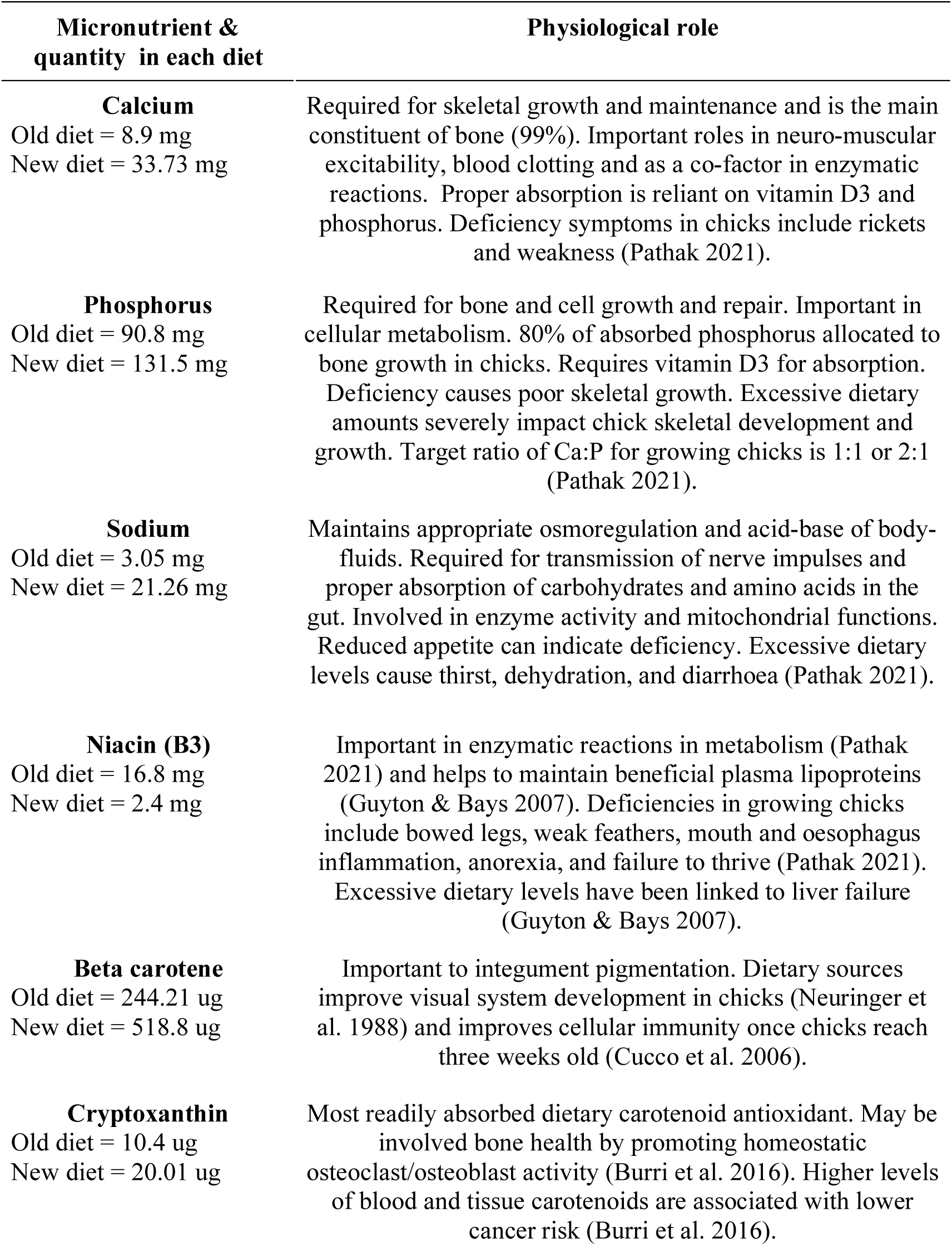
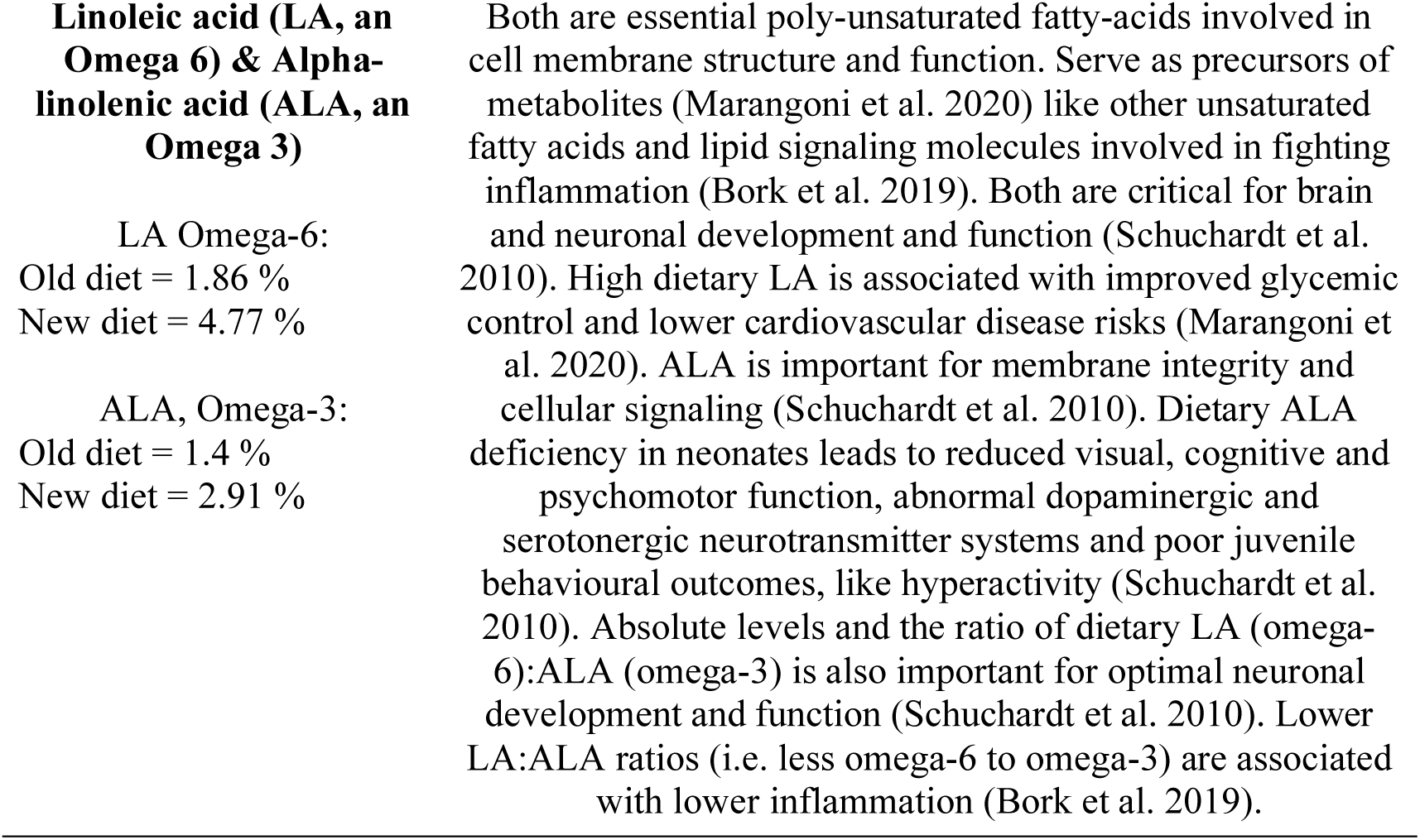
Summary of the main physiological roles of micronutrients that had substantially differed levels in the new Massey ZAA kiwi maintenance diet compared to the old kiwi maintenance diet.

### Other Operation Nest Egg metrics and analysed life history traits

We found no relationship between hatch mass or egg size and the number of days it took kiwi chicks reach release from the brooder room, irrespective of diet, sex, or genetic group. Therefore, from an ONE management perspective, keepers do not need to make allowances for any particular group to remain in the brooder room for more or less time. Relevant to big picture ONE planning however, is our finding that chicks who were malpositioned as embryos, and were therefore more likely to require an assisted hatch, were also more likely to require extended periods of hand-feeding in the brooder room. All three of these occurrences necessitate that keepers conduct intensive, “interventionist” work. Malpositioned embryos require more candling and monitoring (sometimes through X-rays) than normal presenting embryos, and assist hatches and hand-feeding are delicate, skillful, time-consuming activities. Our comparisons identified that Eastern genetic group eggs were more likely to have malpositioned chicks (followed by Coromandel, Western, Haast, and rowi), and while no particular Eastern sires were identified as producing more or less maplositioned embryos, our finding indicate that some breeding pairs could be removed from ONE. While our current analysis was coarse, individual males with family histories of malpositioned eggs (or other congenital health issues) could be readily identified from NKH ONE records. Given the likely stress experienced by the malpositioned embryo, and burden of care on keepers (and cost of care), it may be advisable to identify specific males who produce high numbers of problem offspring, remove their ONE transmitters, and focus conservation effort on other birds.

Within Brown kiwi (*A. mantelli*) we identified no differences among the first and second clutch, or the first and second egg, or sex of the embryo in egg size, chick hatch mass, or chick development time. This suggests that kiwi, unlike species of crested penguins (Davis et al. 2022), gulls (Saino et al. 2010), or wrens (Bowers et al. 2014), have not evolved to apply even subtle brood reduction or sex-specific nutritional allocation strategies to their offspring. Instead, hens appear to invest nutritional resources equally in all their offspring, regardless of time of year, and regardless of the short duration between the first and second egg (see Fuller 1990). This is an impressive feat for Brown kiwi hens, who must secure adequate resources to sustain production of up to four eggs a season. Perhaps related to this demand on hens, is our discovery that Coromandel birds have the smallest eggs, lose nearly as much weight post-hatch as Haast tokoeka, and have a tend to have smaller chicks than the other genetic groups. While we need to extract and analyse more records to confirm some of these patterns, the results support our conjectural ideas about the potential nutritional environment instability of habitats in the Coromandel. In Emu (*Dromaius novaehllandiae*), another ratite, smaller eggs hatch chicks with larger yolks. This is due to less overall maternal resource investment in the eggs, as better provisioned hens lay larger eggs which generate larger hatchlings with less residual yolk (Dzialowski and Sotherland 2004). Greater maternal investment in yolk, rather than over-all hatchling size is thought to be a strategy to increase hatchling survival under food scarcity in precocial birds (see Dzialowski and Sotherland 2004; Dawson and Clark 1996; Anderson and Alisauskas 2001). The more unstable climate of the Coromandel could be driving subtle life history differences in Coromandel kiwi, whereby chicks have evolved greater growth efficiency and hens, greater thrift with respect to their maternal investments. These patterns could be investigated by comparing NKH Coromandel ONE records to regional climatic records.

## ACKNOWLEDGEMENTS

Thank you to the NKH, Richard Benton and the West Coast Wildlife Centre. David Raubenheimer for early discussions on approaches to data extraction. Marla Sokolowski and Matthew Renner for comments on the manuscript. Matthew Renner and Anne Gray for their enduring support, encouragement, and Douglas care.

## Supplementary tables for Gray et al. 1716

**Table S1.** Micronutrient composition for kilogram of insectivore mix “Kiwi Pre-mix” manufactured by Biovac New Zealand. This mix was added to both the old and new kiwi maintenance diets at the rate of 2 g pre-mix / 100 g of prepared diet.

| Micronutrient | Levels / kg | Unit |
| --- | --- | --- |
| Vitamin A (Retinol) | 735000 | iu |
| Vitamin D3 | 62000 | iu |
| Vitamin E | 5300 | iu |
| Vitamin K | 375 | mg |
| Thiamin (B1) | 356 | mg |
| Riboflavin (B2) | 250 | mg |
| Pyridoxine (B6) | 250 | mg |
| Vitamin B12 | 2500 | mg |
| Vitamin C | 4580 | mg |
| Panthothenic Acid (B5) | 1350 | mg |
| Folic Acid (folate pre-cursor) | 80 | mg |
| Biotin (B7) | 20 | mg |
| Niacin (B3) | 1750 | mg |
| Inositol | 10500 | mg |
| Choline | 30000 | mg |
| Manganese | 2250 | mg |
| Iron | 2000 | mg |
| Copper | 135 | mg |
| Zinc | 2000 | mg |
| Iodine | 37.5 | mg |
| Cobalt | 50 | mg |
| Selenium | 15 | mg |

**Table S2.** Generalised linear mixed model comparing the growth efficiency (mass change per unit daily food intake) of *A. mantelli* “Coromandel” kiwi chicks raised on the old *vs.* new kiwi maintenance diets.

| Fixed effects | Estimate | Std. Error | df | t-value | P |
| --- | --- | --- | --- | --- | --- |
| (Intercept) | 0.9409 | 0.1079 | 2.967e+01 | 8.717 | <0.0001 |
| Days | 0.03059 | 0.004932 | 3.411e+01 | 6.201 | <0.0001 |
| Diet*days | 0.0006209 | 0.001963 | 1.064e+03 | 0.316 | 0.752 |

**Table S3.** Generalised linear mixed model comparing the growth efficiency (mass change per unit daily food intake) of *A. mantelli* “Western” kiwi chicks raised on the old *vs.* new kiwi maintenance diets.

| Fixed effects | Estimate | Std. Error | df | t-value | P |
| --- | --- | --- | --- | --- | --- |
| (Intercept) | 0.8470 | 0.09924 | 3.118e+01 | 8.535 | <0.0001 |
| Days | 0.03765 | 0.004853 | 0.3794e+01 | 7.758 | <0.0001 |
| Diet*days | -0.006046 | 0.001986 | 1.107e+03 | -3.045 | 0.00238 |

**Table S4.** Generalised linear mixed model comparing the growth efficiency (mass change per unit daily food intake) of *A. mantelli* “Eastern” kiwi chicks raised on the old *vs.* new kiwi maintenance diets.

| Fixed effects | Estimate | Std. Error | df | t-value | P |
| --- | --- | --- | --- | --- | --- |
| (Intercept) | 0.9475 | 0.07865 | 3.534e+01 | 12.046 | <0.0001 |
| Days | 0.03012 | 0.003489 | 4.354e+01 | 8.634 | <0.0001 |
| Diet*days | -0.004101 | 0.001657 | 1.590e+03 | -2.474 | 0.0135 |

**Table S5.** Generalised linear mixed model comparing the growth efficiency (mass change per unit daily food intake) of Ōkārito kiwi chicks (*A. rowi*) raised on the old *vs.* new kiwi maintenance diets.

| Fixed effects | Estimate | Std. Error | df | t-value | P |
| --- | --- | --- | --- | --- | --- |
| (Intercept) | 0.9082 | 0.1031 | 3.299e+01 | 8.896 | <0.0001 |
| Days | 0.03007 | 0.004024 | 3.852e+01 | 7.474 | <0.0001 |
| Diet*days | -0.01243 | 0.003169 | 1.069e+03 | -3.924 | <0.0001 |

**Table S6.**
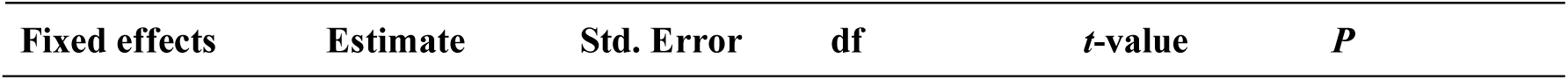
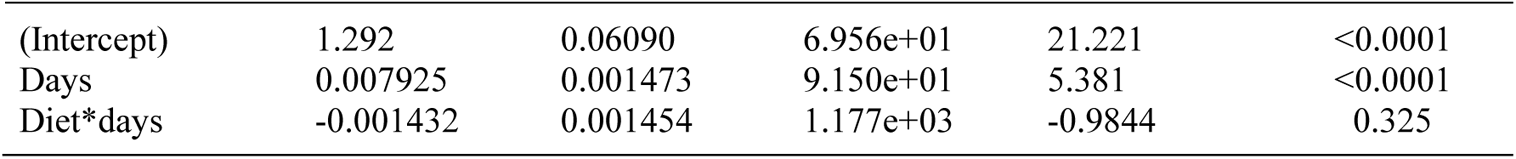
Generalised linear mixed model comparing the growth efficiency (mass change per unit daily food intake) of Haast tokoeka kiwi chicks (*A. australis* “Haast”) raised on the old *vs.* new kiwi maintenance diets.

**Table S7.**
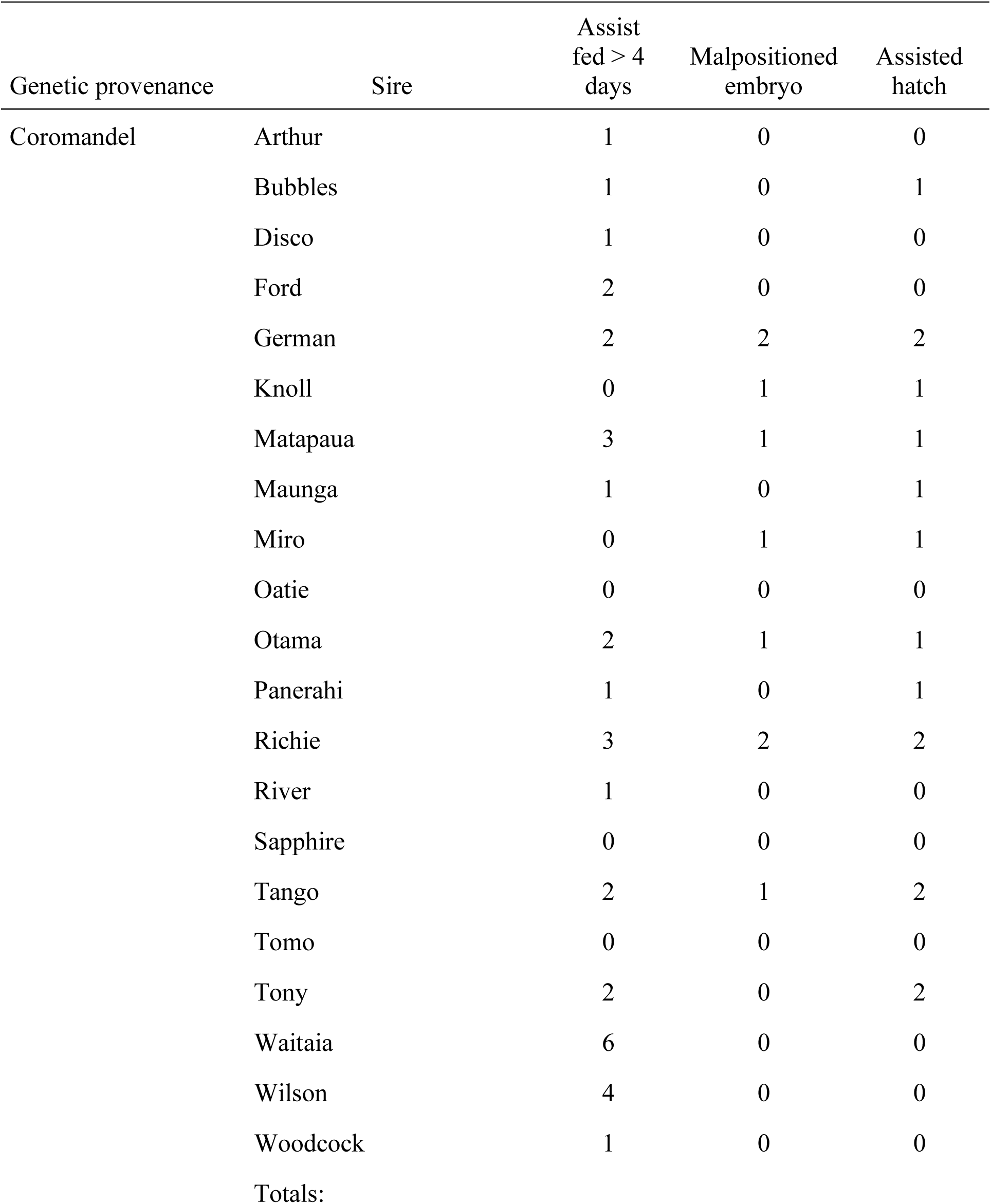
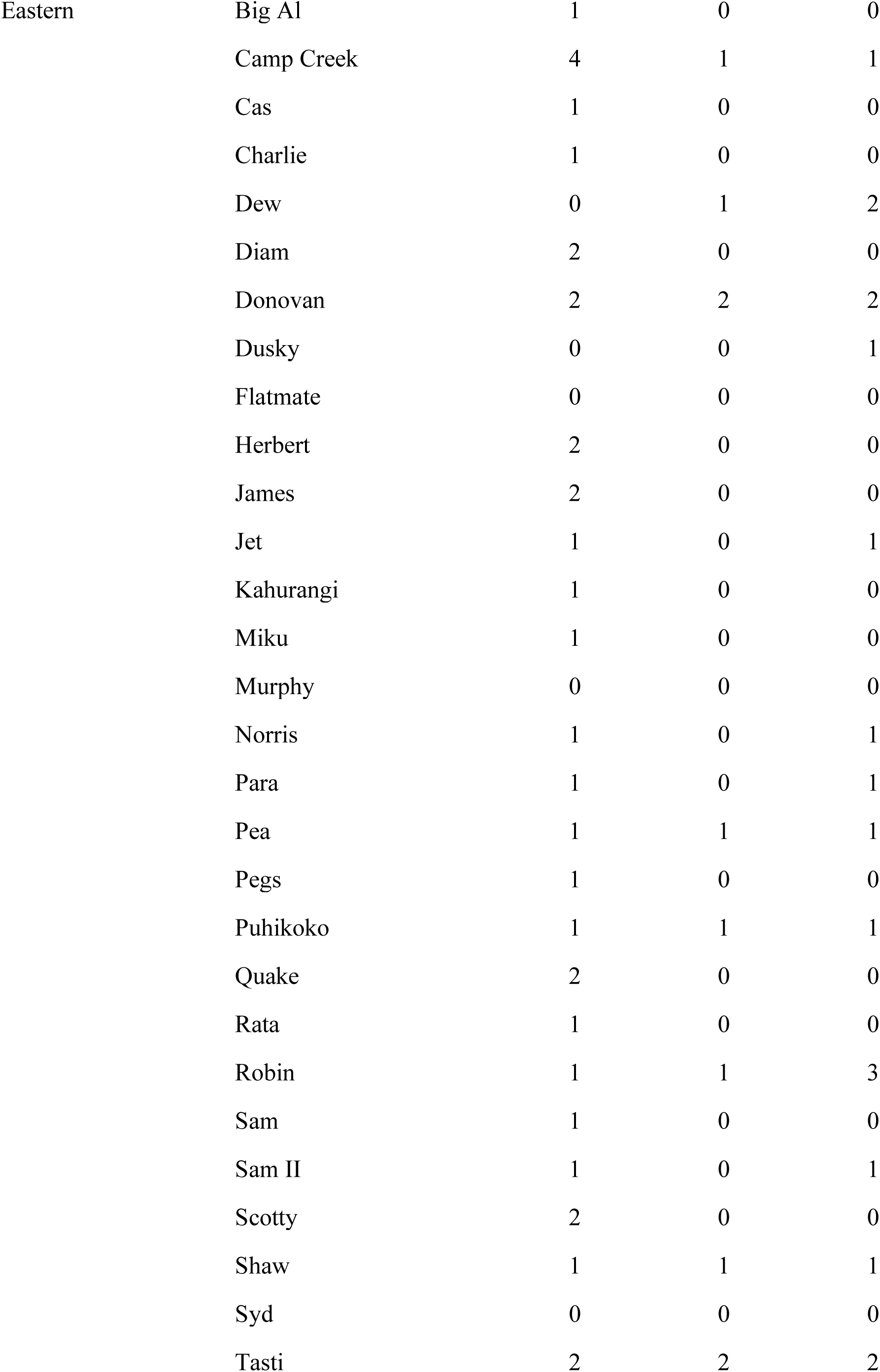
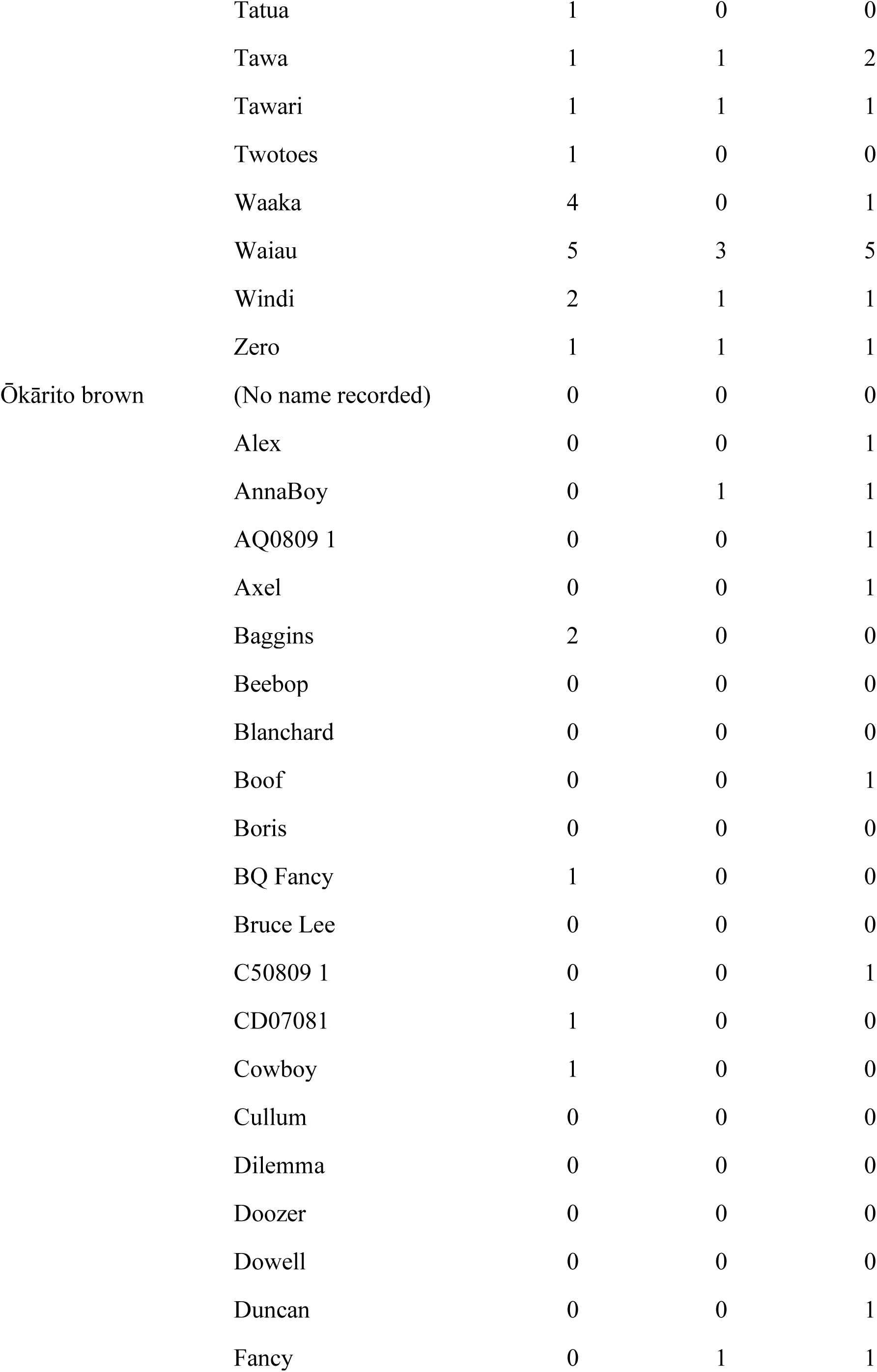
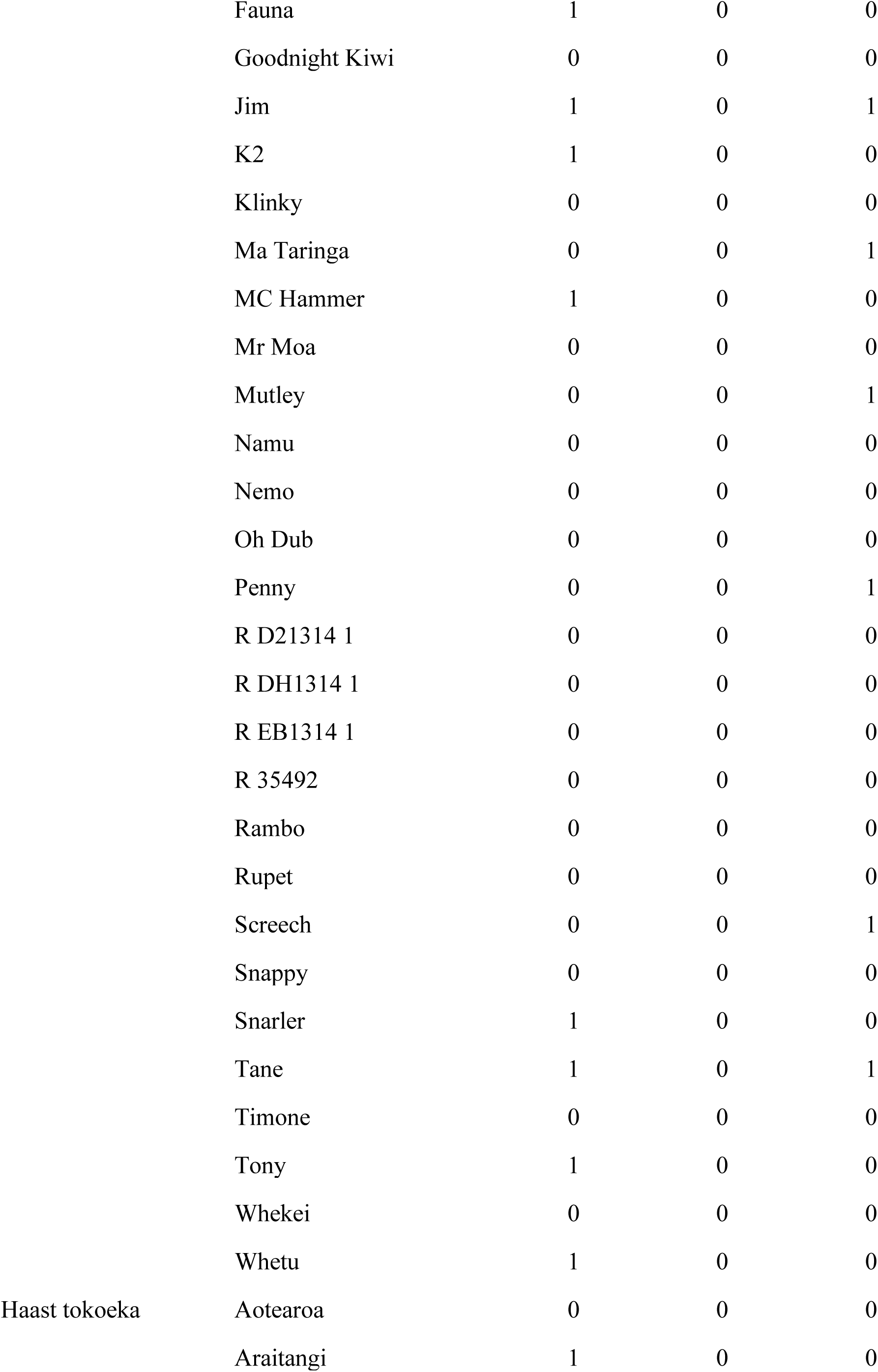
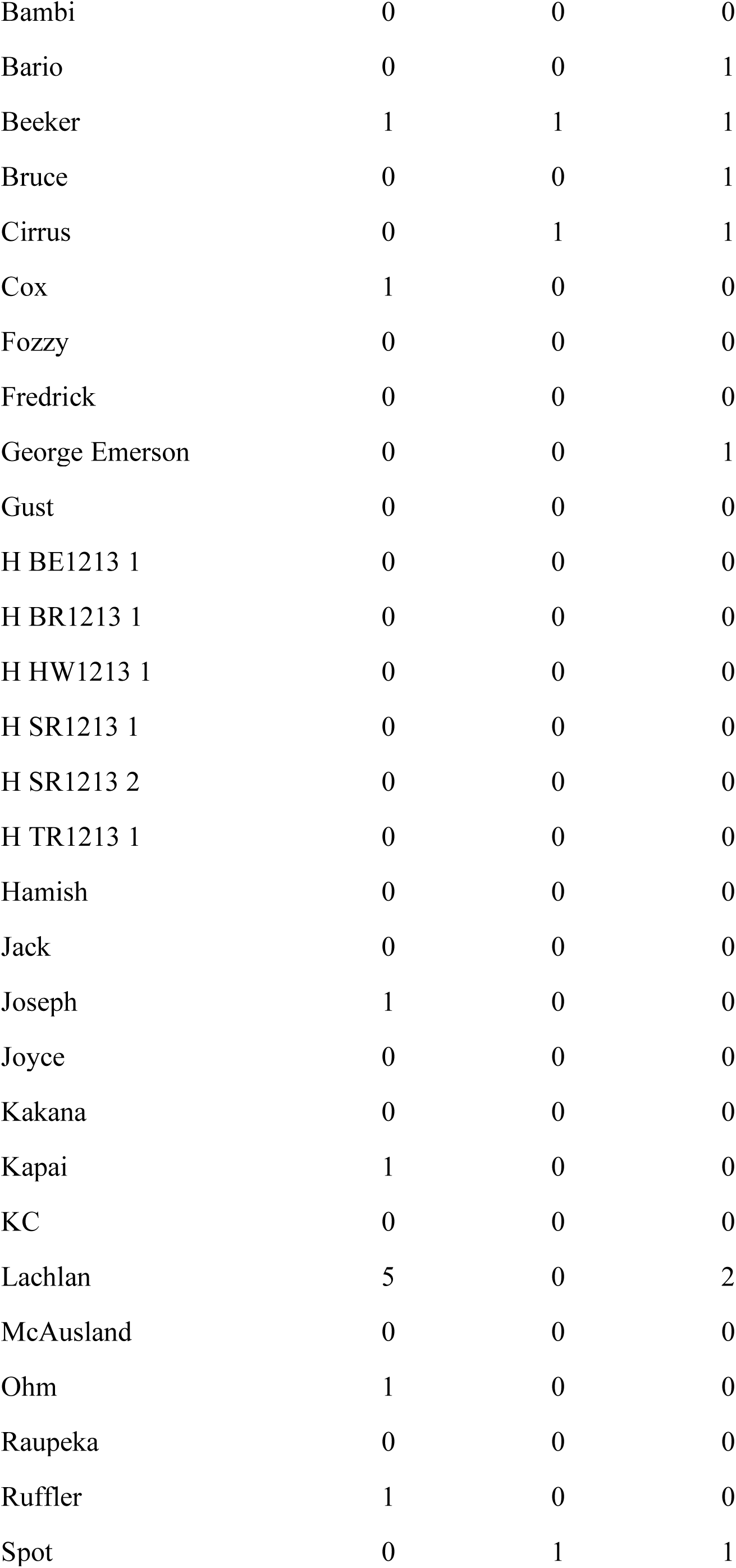
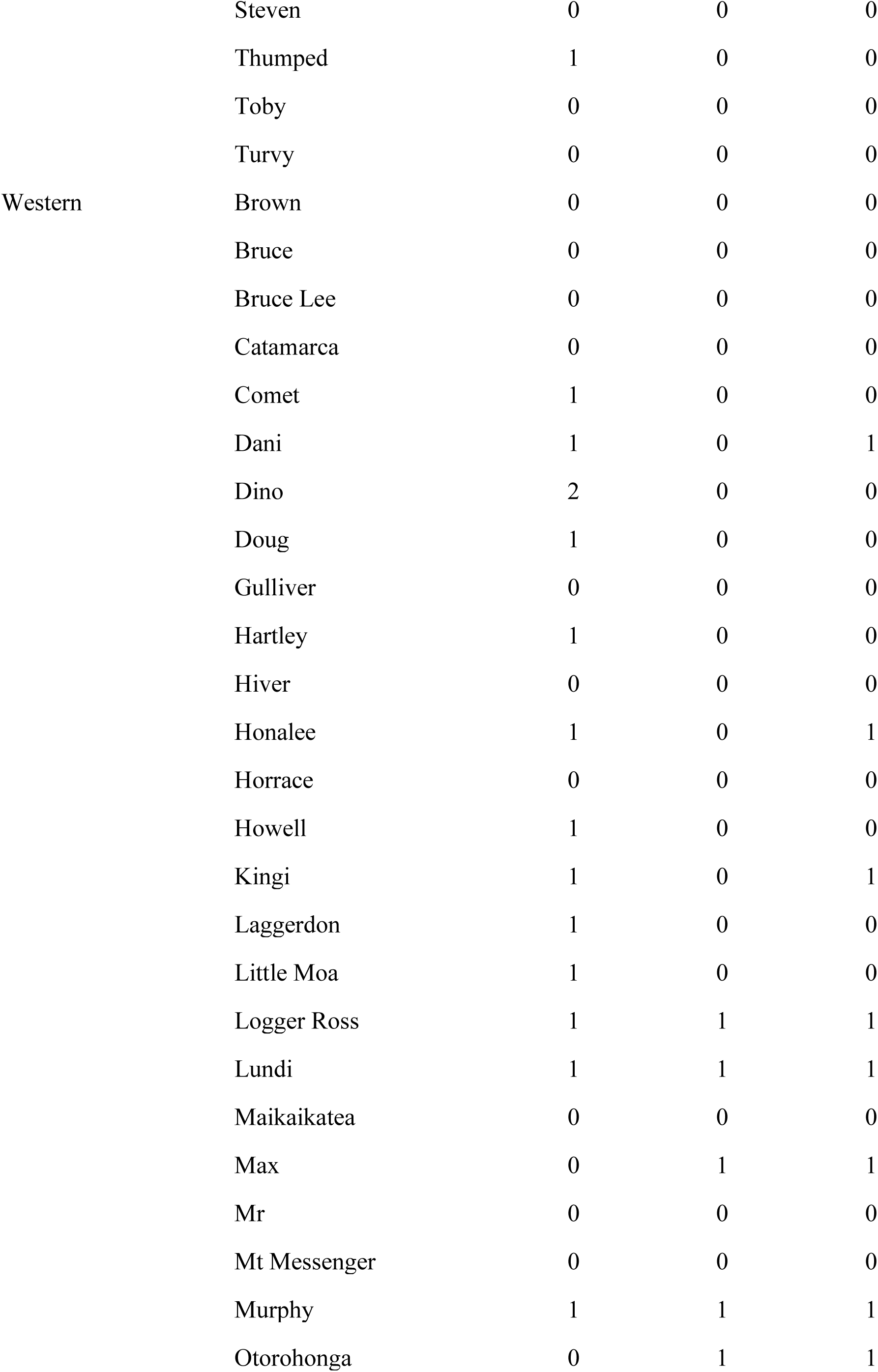
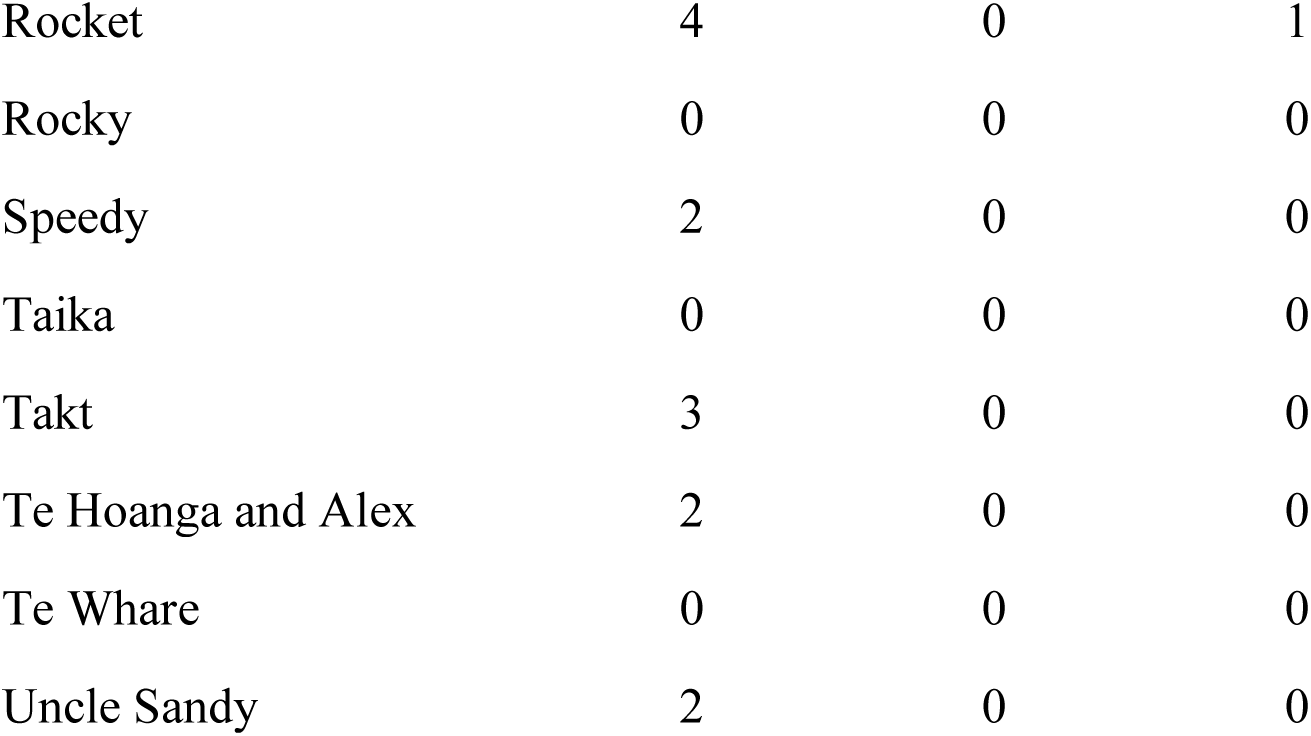
Contingency table showing number of assist fed, malposition, and assisted hatch chicks from kiwi sires of each genetic group. The count data were analysed using Chi-square tests.

## Notes

### Competing Interest Statement

The authors have declared no competing interest.

